# Genetically Engineered Biomimetic Nanozymes Reprogram Immune Niches to Intercept Colitis-Carcinoma Transition

**DOI:** 10.64898/2026.03.12.709998

**Authors:** Qi Sun, Wanling Liu, Sijie Li, Xiwen Chen, Hanjie Zhang, Dongze Mo, Pinwen Zhou, Xiaomiao Cui, Di Huang, Bing Xia, Jingjing Zhang, Xinying Wang, Xiaoyu Wang, Hui Wei

## Abstract

Colitis-associated colorectal cancer (CAC) arises from chronic inflammatory niches characterized by persistent oxidative stress and dysregulated immune cell recruitment. Current anti-inflammatory therapies provide only transient symptom relief and fail to prevent malignant progression due to their inability to simultaneously mitigate oxidative injury and immune chemotaxis. Our analysis of clinical samples revealed markedly elevated C-X-C motif chemokine ligand 2 (CXCL2) in intestinal tissues from patients with inflammatory bowel disease (IBD) and CAC, implicating the CXCL2-CXCR2 axis in driving excessive neutrophil and macrophage infiltration and fostering tumorigenesis. Herein, we report a biomimetic nanozymes (PB@ECM) that reprogram inflammatory immune niches to intercept CAC progression. PB@ECM integrates a Prussian blue nanozyme core with genetically engineered macrophage membranes, enabling concurrent scavenging of reactive oxygen species through superoxide dismutase- and catalase-like activities and sequestration of CXCL2 via membrane-displayed CXCR2 receptors. In murine models of colitis and CAC, PB@ECM significantly alleviated intestinal inflammation, suppressed neutrophil and macrophage infiltration, and effectively inhibited colitis-carcinoma transition. By disrupting the pathological crosstalk between oxidative stress and immune chemotaxis, this work establishes a biomimetic nanozyme strategy for preventing inflammation-driven carcinogenesis.

## Introduction

Chronic inflammation has long been recognized as a central driver of cancer initiation and progression^1–6^. Persistent inflammatory microenvironments sustain excessive production of reactive oxygen species (ROS) and proinflammatory mediators, leading to DNA damage, genomic instability, and oncogenic mutations^6, 7^. Concurrently, prolonged infiltration of innate immune cells, particularly neutrophils and macrophages, further accelerates tumorigenesis through continuous secretion of cytokines such as interleukin-1β (IL-1β) and tumor necrosis factor-α (TNF-α), which activate oncogenic signaling pathways, promote epithelial proliferation, and suppress apoptosis. Despite this well-established causal link, effective strategies to intercept the transition from chronic inflammation to cancer remain scarce, largely because current therapeutic paradigms focus on symptomatic inflammation control while overlooking key immune cell-driven malignant transformation^8, 9^.

Colitis-associated colorectal cancer (CAC) represents a prototypical and clinically devastating model of inflammation driven carcinogenesis^10^. Emerging from long-standing inflammatory bowel diseases (IBD)^11^, including ulcerative colitis^12^ and Crohn’s disease^13^, CAC follows a chronic inflammation dominated oncogenic trajectory that is distinct from sporadic colorectal cancer and is associated with poor prognosis^14, 15^. Existing IBD therapies, such as nonsteroidal anti-inflammatory drugs and biologics, primarily suppress inflammatory symptoms and often provide only transient benefit. In patients with sustained disease activity, extensive immune cell infiltration, and repeated inflammatory insults, merely attenuating inflammation is insufficient to halt malignant transformation. These limitations underscore an unmet need for therapeutic strategies that go beyond inflammation relief and actively disrupt immune cell-driven mechanisms propelling chronic inflammation toward cancer.

Chemokine-mediated immune cell recruitment has emerged as a central yet underexplored contributor to inflammation-driven carcinogenesis^16, 17^. Through analysis of clinical samples and interrogation of public transcriptomic databases, we identified the C-X-C motif chemokine ligand 2 (CXCL2)-CXCR2 signaling axis as a key driver of immune cell recruitment during the IBD-to-CAC transition. CXCL2, abundantly secreted by inflamed intestinal tissues, promotes excessive infiltration of neutrophils and macrophages via CXCR2 engagement, establishing a feed-forward loop that sustains tissue injury and fosters a tumor-promoting immune niche^18, 19^. We therefore hypothesized that effective interception of colitis-to-cancer progression requires simultaneous suppression of inflammation-associated oxidative stress and blockade of CXCL2-CXCR2 driven immune cell recruitment.

Here, we report a genetically engineered biomimetic nanozyme that integrates these two functions into a single therapeutic platform (Figure 1). By coating genetically engineered macrophage membranes (ECM) onto Prussian blue nanozymes (PB), we developed PB@ECM to simultaneously scavenge inflammatory ROS and neutralize CXCL2. PB, an FDA-approved material for metal detoxification, exhibits intrinsic superoxide dismutase (SOD)-, catalase (CAT)-, and hydroxyl radical-scavenging activities, enabling efficient mitigation of oxidative stress in inflamed tissues^20, 21^. Meanwhile, the ECM is engineered to overexpress CXCR2, allowing selective sequestration of CXCL2 and suppression of excessive neutrophil and macrophage infiltration. While PB alone effectively alleviates colitis by reducing oxidative injury, it fails to prevent progression from colitis to colorectal cancer. In contrast, PB@ECM synergistically integrates antioxidant and immunomodulatory functions, thereby not only relieving intestinal inflammation but also intercepting the inflammation-to-cancer transition. Using murine models of colitis and CAC, we demonstrate that PB@ECM markedly attenuates inflammatory injury, reduces innate immune cell infiltration, and effectively prevents the development of colitis-associated colorectal cancer (Figure 1). Together, this work establishes a biomimetic nanozyme strategy for reprogramming inflammatory immune niches to block inflammation-driven carcinogenesis.

**Figure 1.**
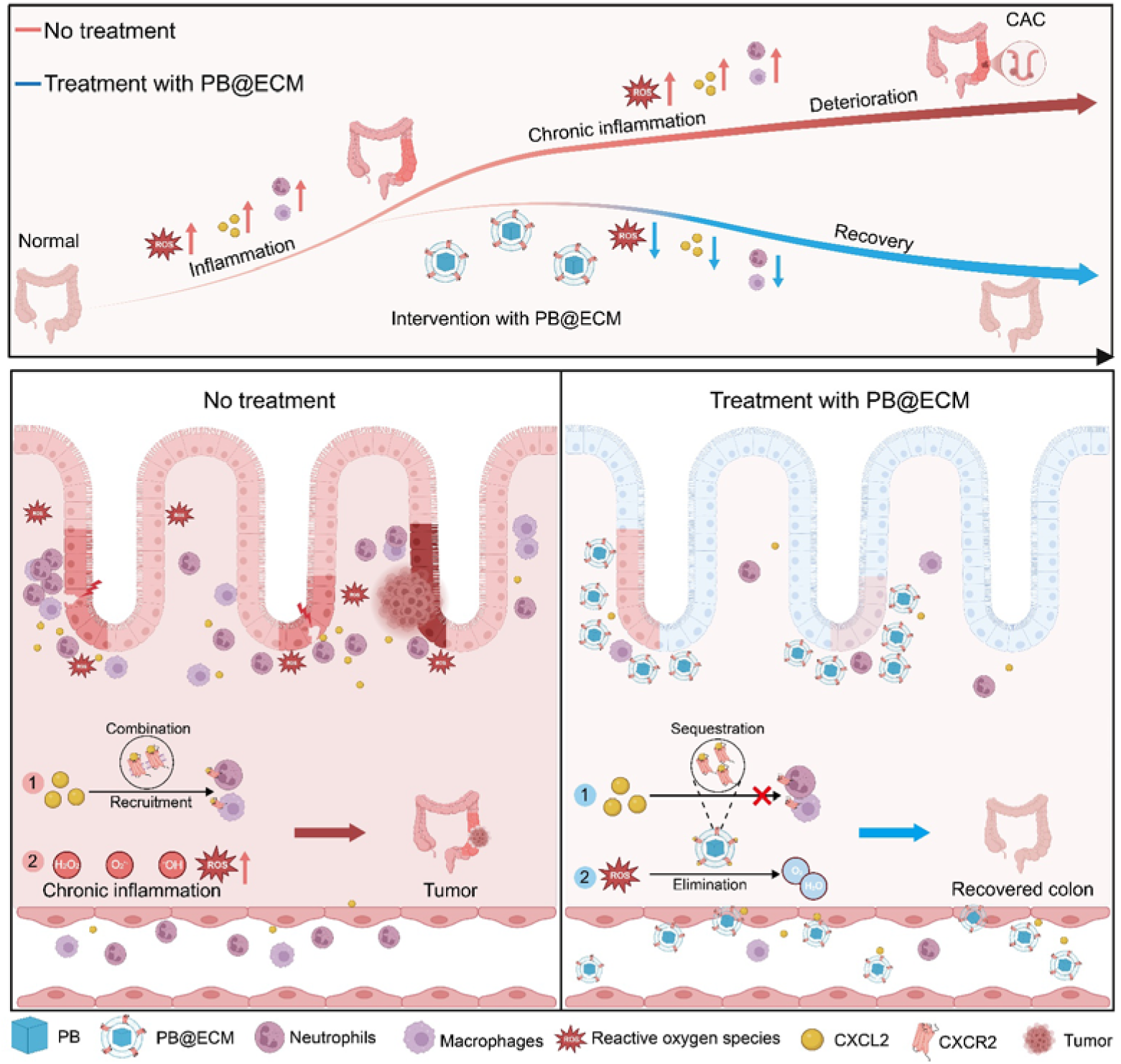
Schematic illustration of PB@ECM nanoformulation for intercepting transition from chronic colitis to colorectal cancer. Without proper treatment, chronic inflammation in IBD elevates ROS and releases chemokines that recruit immune cells, ultimately driving malignant progression to colorectal cancer. With dual functional PB@ECM treatment, the macrophage membrane coating with overexpressed CXCR2 enables precise targeting of colonic inflammation sites. The PB core scavenges ROS, while ECM, *via* CXCR2, sequesters CXCL2 and inhibits the CXCL2-CXCR2 axis-mediated recruitment of neutrophils and macrophages. Together, PB@ECM reduces ROS-mediated damage and suppresses recruitment mediated pro-inflammatory response, thereby alleviating the malignant transformation from chronic colitis to colorectal cancer.

## Results and Discussion

### CXCL2 Overexpression and Immune Cell Infiltration in IBD and CAC Patients

Alleviating oxidative stress alone is insufficient to prevent chronic inflammation from progressing to cancer. Instead, multiple therapeutic actions targeting pathological processes are required to halt this deterioration. To characterize the pathological environment at lesion sites in IBD and CAC, we performed transcriptome sequencing on colon tissues from patients with IBD, CAC, and healthy control (Figure 2A, Supplementary Tables 1 and 2). Principal component analysis revealed distinct transcriptomic profiles between the normal and diseased groups (Supplementary Figures 1 and 2). Pro-inflammatory genes, including *IL-1*β and *IL-1*α, were markedly upregulated in both IBD and CAC tissues (Supplementary Figures 1D and 2D). Notably, *CXCL2* expression was significantly increased in both IBD and CAC patients compared to healthy controls (Figures 2B, C and Supplementary Figure 2E). Consistent with these transcriptomic findings, the enzyme linked immunosorbent assay (ELISA) confirmed elevated CXCL2 secretion in colonic tissues from patients with IBD (Figure 2D) and CAC (Figure 2E). Gene Ontology (GO) analysis further indicated enhanced biological processes related to chemotaxis and migration of neutrophil and macrophage at lesion sites (Supplementary Figure 2F).

**Figure 2.**
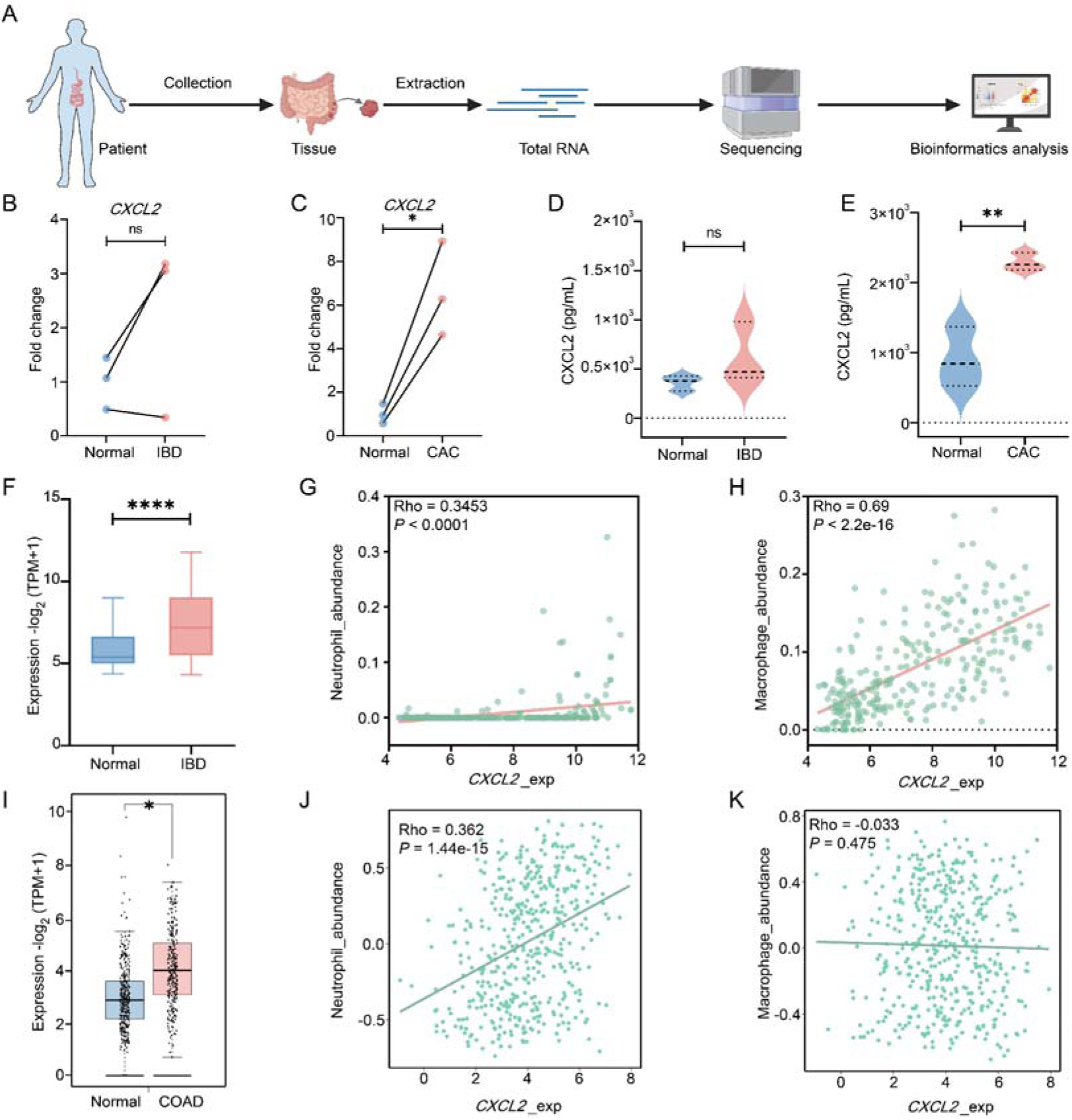
Correlation between CXCL2 expression and immune cell infiltration in IBD and CAC patients. **A**, Flowchart of transcriptome sequencing for colonic tissues from patient. **B**, Gene expression of *CXCL2* in colon tissues from normal and IBD patients. **C**, Gene expression of *CXCL2* in colon tissues from normal and CAC patients. **D**, ELISA measurement of CXCL2 levels in colon tissues from normal and IBD patients (n = 3). **E**, ELISA measurement of CXCL2 levels in colon tissues from normal and CAC patients (n = 3). **F**, Gene expression of *CXCL2* in colon tissues from normal and IBD patients (Normal = 99, IBD = 265). **G**, Correlation analysis examining relationships between neutrophil and *CXCL2* in IBD patients (n = 265). **H**, Correlation analysis examining relationships between macrophage and *CXCL2* in IBD patients (n = 265). **I**, Gene expression of *CXCL2* in colon tissues from normal (n = 349) and COAD (n = 275) patients. **J**, Correlation analysis examining relationships between neutrophil and *CXCL2* in CAC patients (n = 459). **K**, Correlation analysis examining relationships between macrophage and *CXCL2* in CAC patients (n = 459). NS, no significant difference, \**P* < 0.05, \*\**P* < 0.01, \*\*\*\**P* < 0.0001.

To validate these observations and reveal correlation between CXCL2 and immune cell infiltration, we conducted further analysis using the disease database. Transcriptome data of IBD patients (GSE66407, Gene Expression Omnibus database) showed elevated neutrophil infiltration and upregulated *CXCL2* expression in inflamed colonic tissue (Figure 2F and Supplementary Figure 3A). Correlation analysis confirmed positive associations between *CXCL2* expression and the infiltration of neutrophil and macrophage in IBD (Figure 2G, H). Similarly, analysis of colon adenocarcinoma (COAD) patients’ samples from The Cancer Genome Atlas Program (TCGA) database using Gene Expression Profiling Interactive Analysis (GEPIA)^22^ revealed high *CXCL2* expression and significant neutrophil infiltration, both associated with unfavorable relapse-free survival (Figure 2I and Supplementary Figure 3B). TISIDB^23^ analysis also revealed a strong positive correlation between *CXCL2* expression and neutrophil infiltration in COAD patients (Figure 2J), whereas macrophage infiltration showed a weak negative correlation (Figure 2K). Together, these results indicated that *CXCL2* was positively correlated with neutrophil and macrophage infiltration during inflammation, but became more closely associated with neutrophil infiltration in tumors. Moreover, *CXCL2* expression positively correlated with *CXCR2* expression in both IBD and COAD tissues (Supplementary Figure 3C, D), highlighting the involvement of CXCL2-CXCR2 axis across disease stages.

Mechanistically, CXCL2 binds to CXCR2 on neutrophils and macrophages, recruiting them to sites of inflammation and tumors^19, 24–28^. These immune cells release ROS and proinflammatory cytokines, which in turn exacerbate inflammation and promote tumorigenesis. Together, these findings suggested CXCL2-drive immune cell infiltration may contribute to the malignant transformation of chronic colitis to colorectal cancer. Therefore, targeting CXCL2 could represent a promising strategy to suppress immune cell infiltration and interrupt the inflammation-to-cancer transition in colonic diseases.

### Preparation and Characterization of PB@ECM

Motivated by the above findings, we designed a biomimetic catalytic nanoformulation, termed “PB@ECM”, by cloaking Prussian blue (PB) nanozymes with engineered cell membranes displaying CXCR2 (Figure 3A). This design enables competitive sequestration of CXCL2, the above-identified chemokine responsible for recruiting neutrophil and macrophage during inflammation and tumorigenesis. To generate the CXCR2-displaying membrane, the RAW264.7 macrophages were transduced with lentiviral vectors encoding CXCR2, together with Zoanthus green fluorescence protein (ZsGreen) and a 3 × Flag for selection and visualization (Figure 3B and Supplementary Figure 4). Following monoclonal screening, a stable CXCR2-overexpressing cell line (CXCR2-RAW) was established and validated by fluorescence microscopy, quantitative reverse transcription polymerase chain reaction (RT-qPCR), and flow cytometry (Figure 3C-E). Engineered cells membranes were isolated from CXCR2-RAW cells by lysis and differential centrifugation. In parallel, PB was synthesized according to established protocol^29^, exhibiting a uniform cubic morphology (Supplementary Figures 5 and 6) and ferricyanide structure (Figure 3F). Subsequently, the ECM and PB were coextruded through 800 nm polycarbonate membranes to obtain PB@ECM. As a control, PB coated with native RAW264.7 cell membrane (PB@CM) was similarly prepared. High resolution transmission electron microscope (HRTEM) identified the lattice spacings of 0.257 nm corresponding to the (400) planes of PB (Figure 3G). High-angle annular dark-field scanning TEM (HAADF-STEM) and the corresponding energy-dispersive spectroscopy (EDS) elemental mappings of a PB nanocube confirmed a length size of approximately 150 nm and an even distribution of Fe, C, N, O, and K elements (Figure 3H). TEM images of ECM vesicles exhibited a rounded cup-shaped structure, whereas PB@ECM showed a typical core-shell structure, in which the cubic PB core was surrounded with a membrane shell (Figure 3I and Supplementary Figures 7-9). Dynamic light scattering (DLS) analysis indicated that the hydrodynamic diameter of PB@ECM (200.8 ± 1.1 nm) was slightly larger than that of un-coated PB nanocubes (191.0 ± 2.1 nm; Figure 3J and Supplementary Figure 10). The zeta potential of PB@ECM (−15.0 ± 0.1 mV) was comparable to that of ECM vesicles (−13.3 ± 0.1 mV), but less negative than that of uncoated PB cores (−33.2 ± 0.6 mV) (Figure 3K). Powder X-ray diffraction (PXRD) patterns of PB and PB@ECM exhibited the typical characteristic peaks of ferricyanide (Figure 3F), which confirmed that the PB retained their crystalline structure post-coating. X-ray photoelectron spectroscopy (XPS) analysis of Fe showed comparable profiles between PB and PB@ECM, indicating no alteration in PB’s chemical state upon membrane coating (Supplementary Figure 11). SDS-polyacrylamide gel electrophoresis (SDS-PAGE) and Western blot analysis further revealed that membrane proteins, including CXCR2, were preserved in PB@ECM (Figure 3L, M), implying successful membrane integration and functional retention from the parental CXCR2-RAW cells. Collectively, these results demonstrated the successful fabrication of PB@ECM, combining the catalytic activity of PB with the chemokine-sequestering capability of CXCR2-displaying membranes.

**Figure 3.**
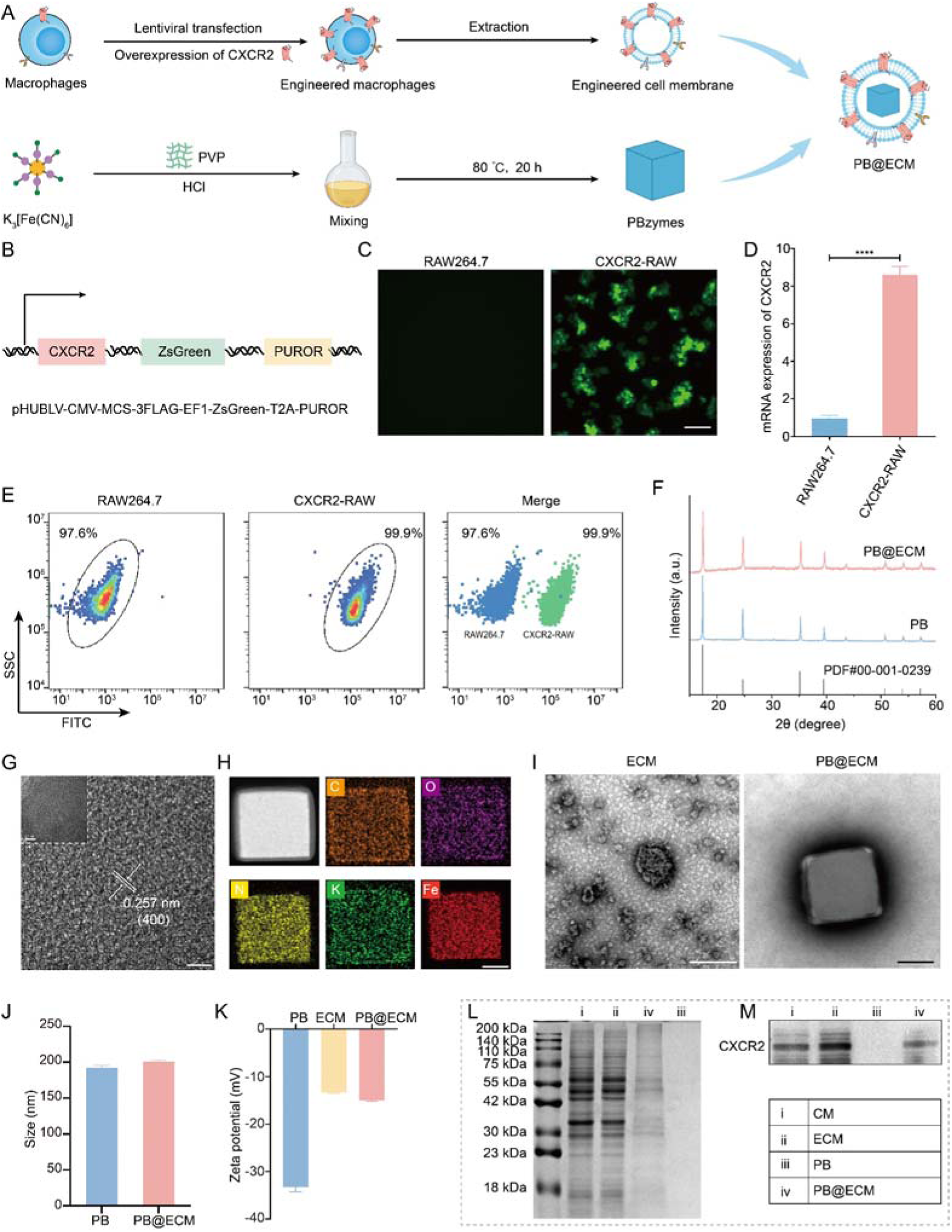
Characterization of PB@ECM. **A**, Schematic illustration of preparation of PB@ECM. **B**, Scheme of retrovirus vector encoding mouse CXCR2. ZsGreen, green fluorescence protein; PUROR, puromycin resistance gene encoding puromycin N-acetyltransferase. **C**, Representative fluorescence images of RAW264.7 and CXCR2-RAW cells, showing establishment of RAW264.7 cells stably expressing mouse CXCR2 on cell membranes (CXCR2-RAW). Left, normal RAW264.7 cells; Right, CXCR2-RAW cells. Scale bar, 50 μm. **D**, RT-qPCR analysis of *CXCR2* expression in RAW264.7 and CXCR2-RAW cells (n = 3). **E**, Representative flow cytometry analysis of green fluorescence protein in RAW264.7 and CXCR2-RAW cells. **F**, XRD patterns of PB and PB@ECM. Standard card, PDF#00-001-0239. **G**, HRTEM image of PB. Scale bar, 2 nm. Inset, low-resolution TEM image. Scale bar, 10 nm. **H**, EDS images of PB. Scale bar, 50 nm. **I**, TEM images of ECM and PB@ECM. Scale bar, 100 nm. **J**, Hydrodynamic diameters of PB and PB@ECM (n = 3). **K**, Zeta potentials of PB, ECM, and PB@ECM (n = 3). **L**, Western blot of CXCR2 in CM, ECM, PB, and PB@ECM. **M**, SDS-PAGE of CM, ECM, PB, and PB@ECM. CM, cell membrane of RAW264.7 cells; ECM, engineered cell membrane of CXCR2-RAW cells. \*\*\*\**P* < 0.0001.

### Enzyme-Like Catalytic Activity of PB@ECM

PB exhibits potent ROS-scavenging capabilities through multiple enzyme-like activities^20^. In this study, we systematically investigated the antioxidant activity of PB@ECM against representative ROS implicated in colitis pathogenesis, including superoxide anion (O_2_^•−^), hydrogen peroxide (H_2_O_2_), and hydroxyl radicals (^•^OH). Next, we evaluated the enzyme-like activities of PB@ECM to determine whether they were affected by ECM coating. To evaluate SOD-like activity, we employed dihydroethidium (DHE) fluorescence assays. PB@ECM treatment led to a concentration-dependent decrease in O_2_^•−^ levels, indicating a superior O_2_^•−^ scavenging ability (Figure 4A and Supplementary Figure 12A). These results were further confirmed using a commercial SOD assay kit (Supplementary Figure 12B). CAT-like activity was assessed using a dopamine oxidation assay, wherein the decomposition of H_2_O_2_ generates O_2_ that promotes the oxidative polymerization of dopamine. A significant increase in absorbance confirmed that PB@ECM efficiently catalyzed H_2_O_2_ breakdown in a dose-dependent manner (Figure 4B). Additionally, we employed electron paramagnetic resonance (EPR) spectroscopy to assess ^•^OH scavenging using 5,5-dimethyl-1-pyrroline N-oxide (DMPO) as a spin-trapping agent. A marked reduction in the EPR signal of the DMPO/^•^OH adducts was observed upon treatment with PB@ECM, confirming its promising ^•^OH clearing ability (Figure 4C). Importantly, the ECM coating did not compromise the intrinsic catalytic activity of PB (Supplementary Figure 13), indicating that the ROS-scavenging capability was well preserved after surface coating. Taken together, these results demonstrated that broad-spectrum ROS-scavenging activity of PB@ECM and underscored its potential for alleviating oxidative stress in colitis-associated inflammatory environments.

**Figure 4.**
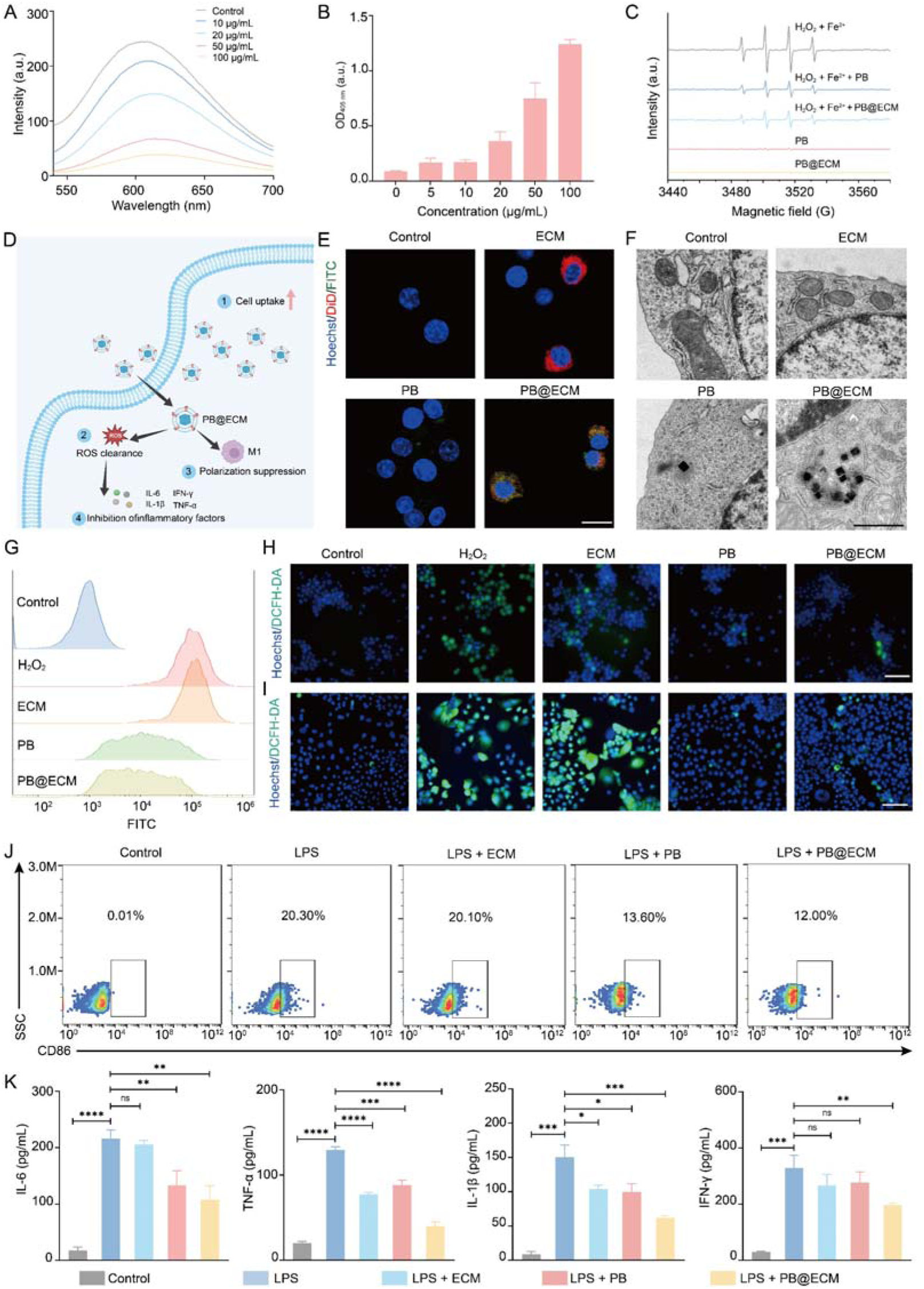
PB@ECM scavenges ROS and protects cells from oxidative stress. **A**, Fluorescence spectra of reaction of HE with X and XO in the absence and presence of PB@ECM. **B**, Quantitative bar chart of dopamine-enabled assay for CAT-like activity of PB@ECM (n = 3). **C**, Scavenging effect of PB and PB@ECM on ^•^OH by EPR. **D**, Scheme of cellular effects of PB@ECM. **E,** Representative fluorescence images showing uptake of ECM, PB, and PB@ECM by RAW264.7 cells. ECM, DiD (red); PB, FITC (green); nuclei, Hoechst (blue). Scale bar, 20 μm. **F**, Representative TEM images showing location of PB and PB@ECM in RAW264.7 cells. Scale bar, 1 μm. **G**, Flow cytometry analysis of intracellular ROS levels in RAW264.7 cells treated with H_2_O_2_, H_2_O_2_ + ECM, H_2_O_2_ + PB, and H_2_O_2_ + PB@ECM. **H**, Confocal fluorescence images of ROS levels in RAW264.7 cells after indicated treatments. **I**, Confocal fluorescence images of ROS levels in Caco-2 cells after indicated treatments. ROS, DCFH-DA (green); nuclei, Hoechst (blue). Scale bar, 100 μm. **J**, Macrophage polarization of CD86 in RAW264.7 cells after indicated treatments, revealed by flow cytometry. **K**, Levels of typical inflammatory cytokines of RAW264.7 cells after indicated treatments (n = 3), including IL-6, TNF-α, IL-1β, and IFN-γ. NS, no significant difference, \**P* < 0.05, \*\**P* < 0.01, \*\*\**P* < 0.001, \*\*\*\**P* < 0.0001.

### Intracellular ROS Scavenging and Anti-Inflammatory Capabilities of PB@ECM

The biocompatibility and cellular uptake of PB@ECM were evaluated to assess their therapeutic potential. Cytotoxicity assays demonstrated that PB@ECM possessed negligible toxicity toward RAW 264.7 macrophages up to 100 μg/mL, comparable to pristine PB (Supplementary Figure 14). Based on the activity test and cell membrane of PB@ECM, we will next investigate its cellular functions (Figure 4D). To investigate cellular internalization, fluorescently labeled FITC-PB and FITC-PB@ECM were synthesized and subsequently incubated with RAW264.7 and Caco-2 cells (Supplementary Figures 15 and 16). Confocal microscopy and TEM analysis revealed enhanced uptake of PB@ECM compared to PB (Figure 4E, F, and Supplementary Figures 17 and 18), likely due to the ECM coating facilitating membrane interaction^30^.

To assess the cytoprotective potential of PB@ECM under oxidative stress, we established an *in vitro* inflammation model by treating RAW264.7 and Caco-2 cells with H_2_O_2_. Following PB@ECM treatment, intracellular ROS level was significantly reduced, as confirmed by DCFH-DA staining via both confocal microscopy and flow cytometry (Figure 4G-I), suggesting that PB@ECM efficiently scavenged ROS and alleviated oxidative stress. At the organelle level, TEM imaging revealed that PB@ECM preserved mitochondrial morphology under oxidative stress, preventing typical damage features such as swelling, matrix brightening, and cristae fragmentation (Supplementary Figure 19). Moreover, PB@ECM exhibited robust anti-inflammatory effects. Flow cytometric analysis indicated that PB@ECM inhibited M1 polarization of macrophage (Figure 4J), and ELISA measurements showed a marked reduction in pro-inflammatory cytokine secretion (Figure 4K). Collectively, these results demonstrated that PB@ECM effectively scavenged ROS, protected cells from oxidative damage, and modulated inflammatory response *in vitro*, supporting their promise for therapeutic applications in intestinal inflammation.

### PB@ECM Sequesters CXCL2 and Inhibits Neutrophil and Macrophage Chemotaxis

To evaluate the CXCL2-sequestering capability of PB@ECM, we incubated CXCL2 with PB@CM or PB@ECM in the absence or presence of the CXCR2 antagonist SB225002. Only PB@ECM effectively sequestered to CXCL2 (Figure 5A), owing to the expression of CXCR2 on the engineered membranes. This sequestration was diminished in the presence of SB225002, confirming the receptor-ligand specificity (Figure 5A). We next assessed whether CXCL2 sequestration could functionally suppress chemokine-driven immune cell migration using a transwell assay. Neutrophils were freshly isolated from murine bone marrow (Figure 5B). Neutrophils or RAW264.7 macrophages were placed in the upper chamber, while the lower chamber contained CXCL2 alone or in combination with PB, PB@CM, PB@ECM, or PB@ECM + SB225002 (Figure 5C). After 24 h, the number of migrated cells was quantified to assess chemotactic inhibition. PB@ECM markedly reduced both neutrophil and macrophage migration (Figure 5E-G), while PB and PB@CM had minimal effects. Notably, the inhibitory effect of PB@ECM was partially abrogated by SB225002, further substantiating that the anti-migratory activity of PB@ECM was mediated by competitive CXCL2 binding through CXCR2 receptors on the engineered membrane (Figure 5E-G). To further examine the chemotactic modulation, a scratch-wound assay was preformed to investigate the impact of PB@ECM on macrophage motility. PB@ECM significantly inhibited CXCL2-induced RAW264.7 cells migration (Figure 5H, I), while PB, PB@CM, and PB@ECM with SB225002 showed negligible effects, indicating the essential role of CXCL2-CXCR2 interactions. Together, these results established PB@ECM as an effective chemokine scavenger capable of sequestering CXCL2 and attenuating neutrophil and macrophage chemotaxis. Beyond its intrinsic ROS-scavenging properties, the dual-functional nanoplatform PB@ECM offered a promising strategy for disrupting inflammatory cell recruitment within CXCL2-enriched microenvironments.

**Figure 5.**
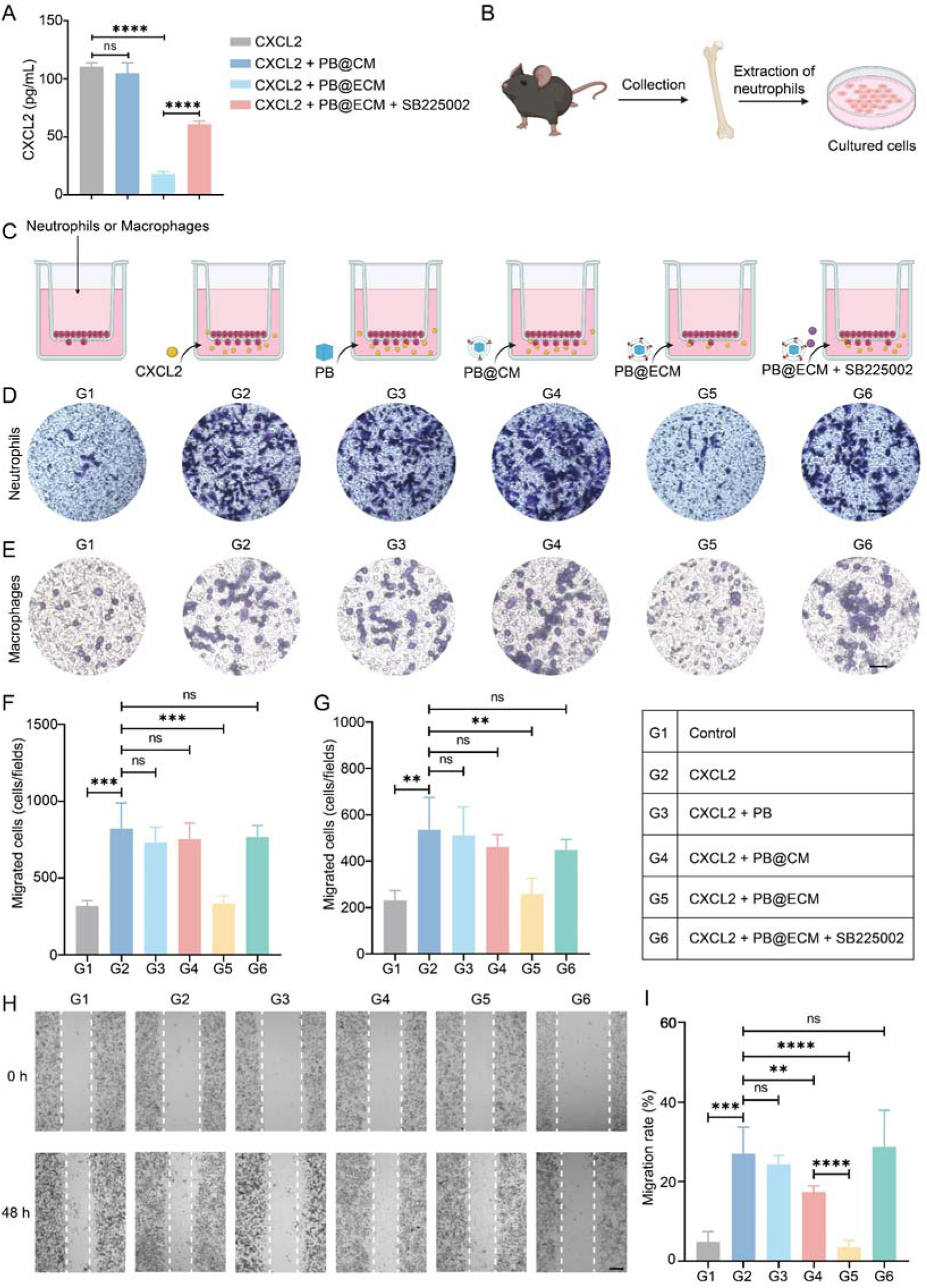
*In vitro* inhibition of neutrophil and macrophage migration by PB@ECM *via* CXCL2-CXCR2 axis. **A**, Content of CXCL2 in supernatant after incubation of CXCL2 solution with PB@CM, PB@ECM, and PB@ECM + SB225002, measured by ELISA (n = 3). **B**, Schematic illustration of neutrophil extraction from mouse bone marrow. **C**, Schematic illustration of transwell experiment. Representative images (**D**) and quantification (**F**) of migratory neutrophil cells stained with crystal violet at 24 h after incubation with CXCL2, CXCL2 + PB, CXCL2 + PB@CM, CXCL2 + PB@ECM, and CXCL2 + PB@ECM + SB225005, determined by transwell assay (n = 5). Scale bar, 50 μm. Representative images (**E**) and quantification (**G**) of migratory RAW264.7 cells stained with crystal violet at 24 h after incubation with CXCL2, CXCL2 + PB, CXCL2 + PB@CM, CXCL2 + PB@ECM, and CXCL2 + PB@ECM + SB225005, determined by transwell assay (n = 5). Scale bar, 50 μm. Representative images (**H**) and quantification (**I**) of migratory RAW264.7 cells at 48 h after incubation with CXCL2, CXCL2 + PB, CXCL2 + PB@CM, CXCL2 + PB@ECM, and CXCL2 + PB@ECM + SB225005 (n = 5). Scale bar, 200 μm. NS, no significant difference, \*\**P* < 0.01, \*\*\**P* < 0.001, \*\*\*\**P* < 0.0001.

### Therapeutic effects of PB@ECM for colitis

Given the potent ROS-scavenging capacity and chemotaxis-inhibitory properties of PB@ECM, we next assessed its therapeutic efficacy in a murine IBD model. Acute colitis was induced in C57BL/6 mice by administering 2.5% (w/v) DSS (dextran sodium sulfate) in drinking water for 7 days, followed by tail vein injections of PBS, ECM, PB, or PB@ECM on days 8 and 10, as outlined in Figure 6A. Body weight changes, a key indicator of colitis progression, were monitored throughout the study (Figure 6B). DSS-induced colitis led to a 23.5% reduction in body weight by day 11, whereas healthy controls displayed continuous weight gain. Notably, treatment with PB@ECM significantly ameliorated weight loss, with only a 10.5% reduction, compared to 15.0% in the PB-treated group (Figure 6B). Additionally, PB@ECM treatment effectively mitigated DSS-induced colon shortening, a hallmark of intestinal inflammation, outperforming PB and ECM groups (Figure 6C and D). Histological analysis further corroborated these findings. Haematoxylin and eosin (H&E) staining revealed that DSS treatment caused marked disruption of the colonic architecture and epithelial integrity. Treatment with PB@ECM substantially mitigated these pathological changes, showing superior tissue preservation and reduced inflammatory cell infiltration compared to other treatment groups (Figure 6E). These findings indicated the superior capacity of PB@ECM in alleviating DSS-induced systemic symptoms. ELISA results revealed that PB@ECM significantly lowered the levels of key proinflammatory cytokines, including IL-6, TNF-α, IL-1β, and IFN-γ (Figure 6F). Furthermore, PB@ECM-treated mice exhibited markedly reduced neutrophil and macrophage infiltration in colonic tissues compared to other treatment groups, demonstrating the chemotaxis-inhibitory properties of PB@ECM by sequestering CXCL2 (Figure 6G and H). These findings suggested that the superior therapeutic efficacy of PB@ECM arose from its combined ability to attenuate inflammation and suppress immune cell accumulation in the colon.

**Figure 6.**
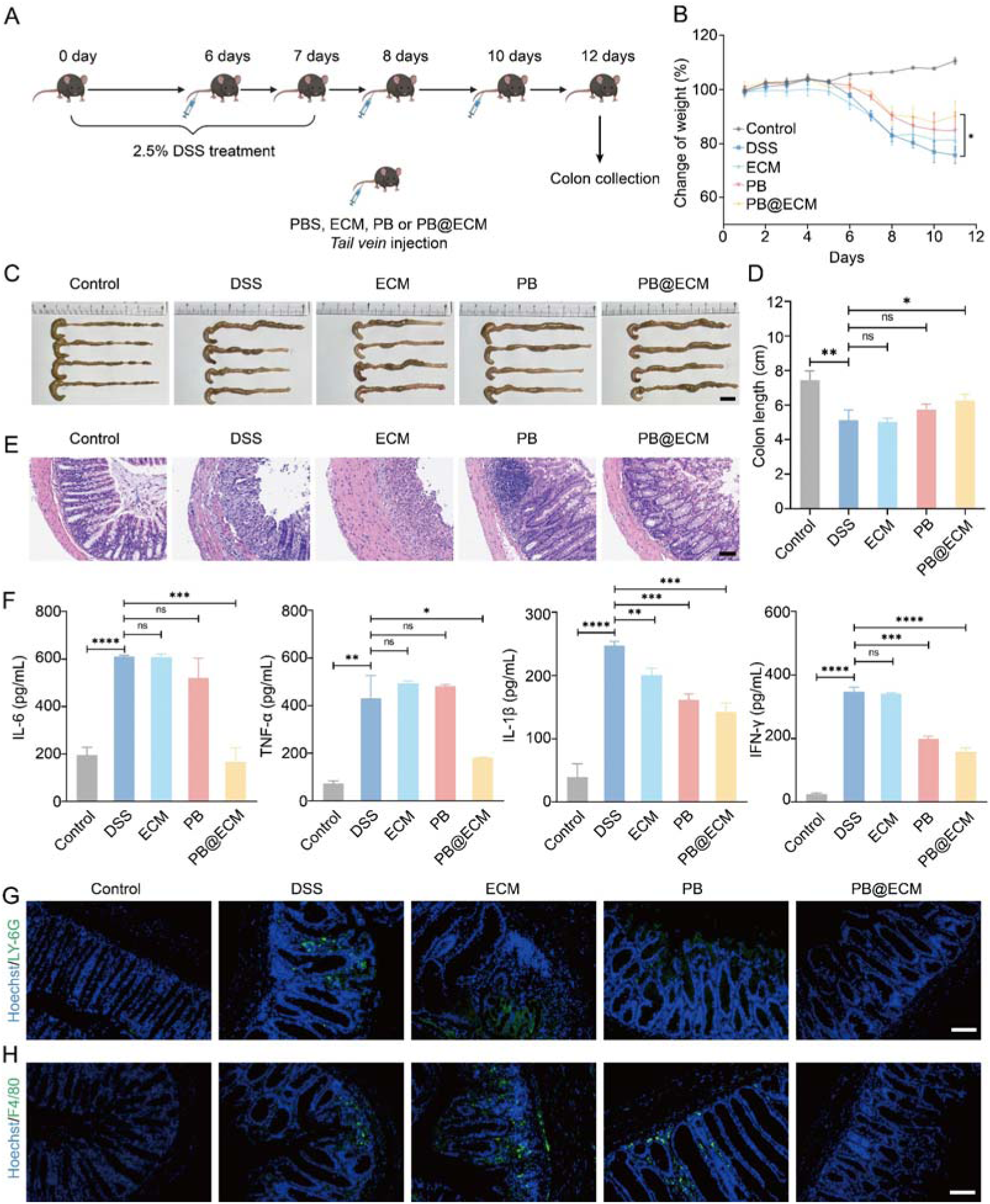
Anti-inflammatory activity and inhibition of neutrophil and macrophage infiltration by PB@ECM in DSS-induced mice. **A**, Schematic illustration of DSS-induced mouse model. **B**, Daily body weight changes of mice for 11 days. **C**, Photographs of colons and (**D**) corresponding colon lengths after indicated treatments (n = 4). Scale bar, 1 cm. **E**, H&E-stained colonic sections after indicated treatments. Scale bar, 100 μm. **F**, Levels of typical inflammatory cytokines in colons on day 12 after indicated treatments by ELISA (n = 3). Representative confocal images of neutrophil (**G**) and macrophage (**H**) in colon tissues. Neutrophil, Ly-6G (green); macrophage, F4/80 (green); nuclei, Hoechst (blue). Scale bar, 100 μm. NS, no significant difference, \**P* < 0.05, \*\**P*< 0.01, \*\*\**P* < 0.001, \*\*\*\**P* < 0.0001.

### PB@ECM Inhibits Colitis to Colon Cancer Progression

CAC develops from chronic IBD, driven by sustained oxidative stress and immune cell infiltration. IBD lesions feature elevated ROS and CXCL2 levels, which together promote epithelial damage and recruit neutrophils and macrophages, accelerating the inflammation-to-cancer transition. Our *in vitro* studies confirmed that PB@ECM catalytic nanoformulation both scavenged ROS and disrupted CXCL2 activity, highlighting their potential to impede CAC progression.

To evaluate the therapeutic benefits of PB@ECM in CAC, we first assessed its targeting ability *in vivo*. After establishing the CAC model using AOM/DSS (AOM, azoxymethane), FITC-PB and FITC-PB@ECM were intravenously injected on day 29, and colons were harvested 6 h post-injection for fluorescence analysis (Supplementary Figure 20A, B). As shown in Supplementary Figures 20C, D, and 21, FITC-PB@ECM exhibited significantly higher accumulation in the colon compared with FITC-PB (11.88 × 10^9^ *vs.* 3.99 × 10^9^ (p/s)/(μW/cm^2^)), attributed to the inflammatory targeting capability of the ECM. The biodistribution analysis further showed predominant clearance *via* the liver and kidney (Supplementary Figure 22).

The therapeutic efficacy of PB@ECM was then evaluated across three stages of disease progression. In the early stage (Figure 7A), no significant differences in colon length or visible tumor formation were observed among the ECM, PB, and PB@ECM groups (Figure 7B, Supplementary Figures 23A, and 24). However, histological analysis revealed markedly alleviated colonic inflammation in the PB@ECM group (Figure 7C), and cytokine profiling further showed that PB@ECM-treated mice had significantly lower levels of IL-6, TNF-α, IL-1β, and IFN-γ compared to the AOM/DSS, ECM, and PB groups (Supplementary Figure 23B), indicating its efficacy in mitigating early inflammatory injury. Additionally, reduced neutrophil and macrophage infiltration was observed by immunofluorescence analysis, accompanied by decreased CXCL2 levels in the colon (Figure 7D, E and Supplementary Figure 23C-E).

**Figure 7.**
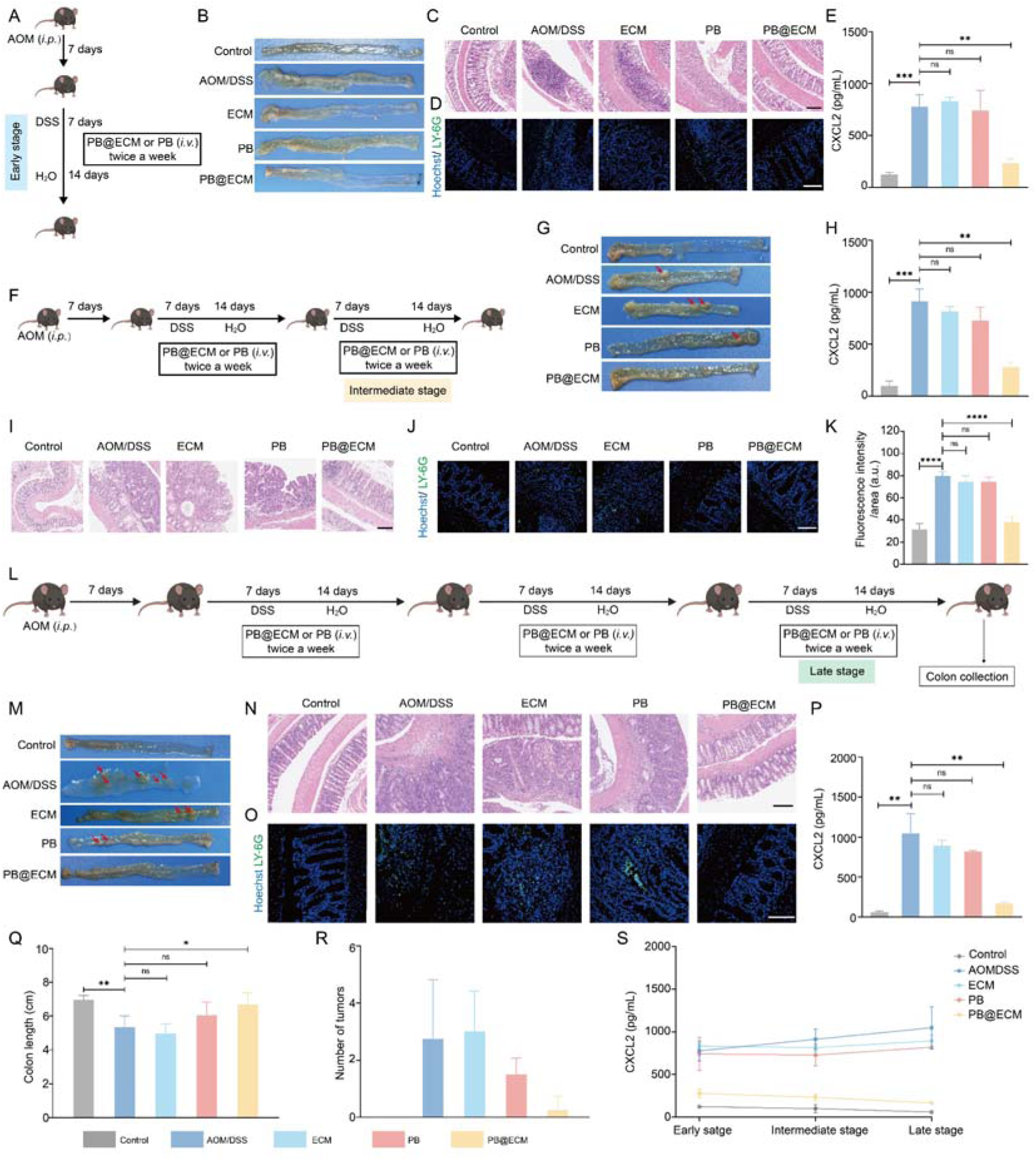
Treatment of CAC mice with PB@ECM. **A**, Schematic illustration for PB@ECM treatment of AOM/DSS mice (early stage). **B**, Representative photographs of colons from Control, ECM, PB, and PB@ECM groups at early stage. **C**, Representative H&E-stained colonic sections from Control, AOM/DSS, ECM, PB, and PB@ECM groups at early stage. Scale bar, 200 μm. **D**, Representative confocal images of neutrophil in colon tissue at early stage. Neutrophil, Ly-6G (green); nuclei, Hoechst (blue). Scale bar, 100 μm. **E**, ELISA of CXCL2 in colons at early stage (n = 3). **F**, Schematic illustration for PB@ECM treatment of AOM/DSS mice (intermediate stage). **G**, Representative photographs of colons from Control, ECM, PB, and PB@ECM groups at intermediate stage. **H**, ELISA of CXCL2 in colons at intermediate stage (n = 3). **I**, Representative H&E-stained colonic sections from Control, AOM/DSS, ECM, PB, and PB@ECM groups at intermediate stage. Scale bar, 200 μm. **J**, Representative confocal images of neutrophil in colon tissue at intermediate stage. Neutrophil, Ly-6G (green); nuclei, Hoechst (blue). Scale bar, 100 μm. **K**, Quantitative analysis of Ly-6G immunofluorescence staining (n = 4). **L**, Schematic illustration for PB@ECM treatment of AOM/DSS-induced CAC mice. **M**, Representative photographs of colons from Control, ECM, PB, and PB@ECM groups at late stage. **N**, Representative H&E-stained colonic sections from Control, AOM/DSS, ECM, PB, and PB@ECM groups at late stage. Scale bar, 200 μm. **O**, Representative confocal images of neutrophil in colon tissue at late stage. Neutrophil, Ly-6G (green); nuclei, Hoechst (blue). Scale bar, 100 μm. **P**, ELISA of CXCL2 in colons at late stage (n = 3). **Q**, Colon lengths in indicated groups at late stage (n = 5). **R**, Number of tumors in indicated groups at late stage (n = 5, n (AOM/DSS) = 4). **S**, Changes of CXCL2 at three different stages (n = 3). NS, no significant difference, \**P* < 0.05, \*\**P* < 0.01, \*\*\**P* < 0.001.

In the intermediate stage (Figure 7F), although colon length remained comparable across groups (Supplementary Figures 25A and 26A), PB@ECM-treated mice did not develop tumor areas, unlike the AOM/DSS, ECM and PB groups (Figure 7G and Supplementary Figure 26B). Cytokine profiling further showed that PB@ECM-treated mice had significantly lower levels of IL-6, TNF-α, IL-1β, and IFN-γ compared to the other groups (Supplementary Figure 25B). Histological analysis revealed markedly inhibited tumor formation in the PB@ECM group compared to the AOM/DSS, ECM, and PB groups (Figure 7I), indicating its efficacy in mitigating colitis to tumorigenesis. Importantly, neutrophil and macrophage infiltration was substantially mitigated in the PB@ECM group due to the reduced CXCL2 levels, which also maintained lower cytokine levels (Figure 7H, J, K, and Supplementary Figure 25B-D), suggesting that PB@ECM effectively limited disease progression from colitis to tumor.

By the late stage (Figure 7L), the therapeutic advantages of PB@ECM became increasingly pronounced. PB@ECM treatment significantly attenuated colon shortening (Figure 7Q and Supplementary Figure 27A) and markedly reduced tumor burden relative to ECM, PB, and untreated groups (Figure 7R and Supplementary Figure 27B). Notably, some PB@ECM-treated mice exhibited complete absence of detectable tumors (Figure 7M and Supplementary Figure 27B). ELISA results of PB@ECM group confirmed sustained suppression of proinflammatory cytokines in the colon (Supplementary Figure 28A), while H&E staining revealed minimal mucosal injury and leukocyte infiltration (Figure 7N). Immunofluorescence analysis further demonstrated significantly reduced infiltration of neutrophil and macrophage (Figure 7O and Supplementary Figure 28B-D), in line with decreased CXCL2 expression (Figure 7P). Remarkedly, CXCL2 levels in the PB@ECM group remained consistently low across all three disease stages, with no significant difference compared to the healthy controls (Figure 7S). Together, these findings highlighted the dual therapeutic effects of PB@ECM in attenuating inflammation and limiting immune cell infiltration, thereby effectively impeding the colitis-to-cancer transition.

Further histological analysis of major organs (heart, liver, spleen, lung, and kidney) revealed no signs of systemic toxicity across all treatment groups (Supplementary Figures 29, 30, and 31), supporting the favorable biosafety profile of PB@ECM.

### Transcriptomic Profiling of PB@ECM**-**Treated CAC Mouse Colons

To elucidate the mechanisms underlying the therapeutic effects of PB@ECM, we performed transcriptomic analysis on colonic tissues from late-stage CAC mice. Principal component analysis revealed distinct transcriptomic profiles among the Control, AOM/DSS, PB, and PB@ECM groups (Figure 8A), indicating marked transcriptional reprogramming upon treatment. Venn diagram analysis further illustrated differentially expressed gene patterns across key comparisons, including AOM/DSS *vs.* Control, PB@ECM *vs.* PB, and PB@ECM *vs.* AOM/DSS groups (Supplementary Figure 32). Specifically, volcano plots identified 973 up-regulated and 861 down-regulated genes in AOM/DSS group compared to Control group (Supplementary Figure 33A), while PB@ECM treatment resulted in 637 up-regulated and 1299 down-regulated genes relative to the PB group (Supplementary Figure 33B). Notably, PB@ECM significantly altered 2030 transcripts compared to the AOM/DSS group, including 830 up-regulated and 1200 down-regulated genes, based on the filtering criteria of |fold change| ≥ 2.0 and P < 0.05 (Figure 8B).

**Figure 8.**
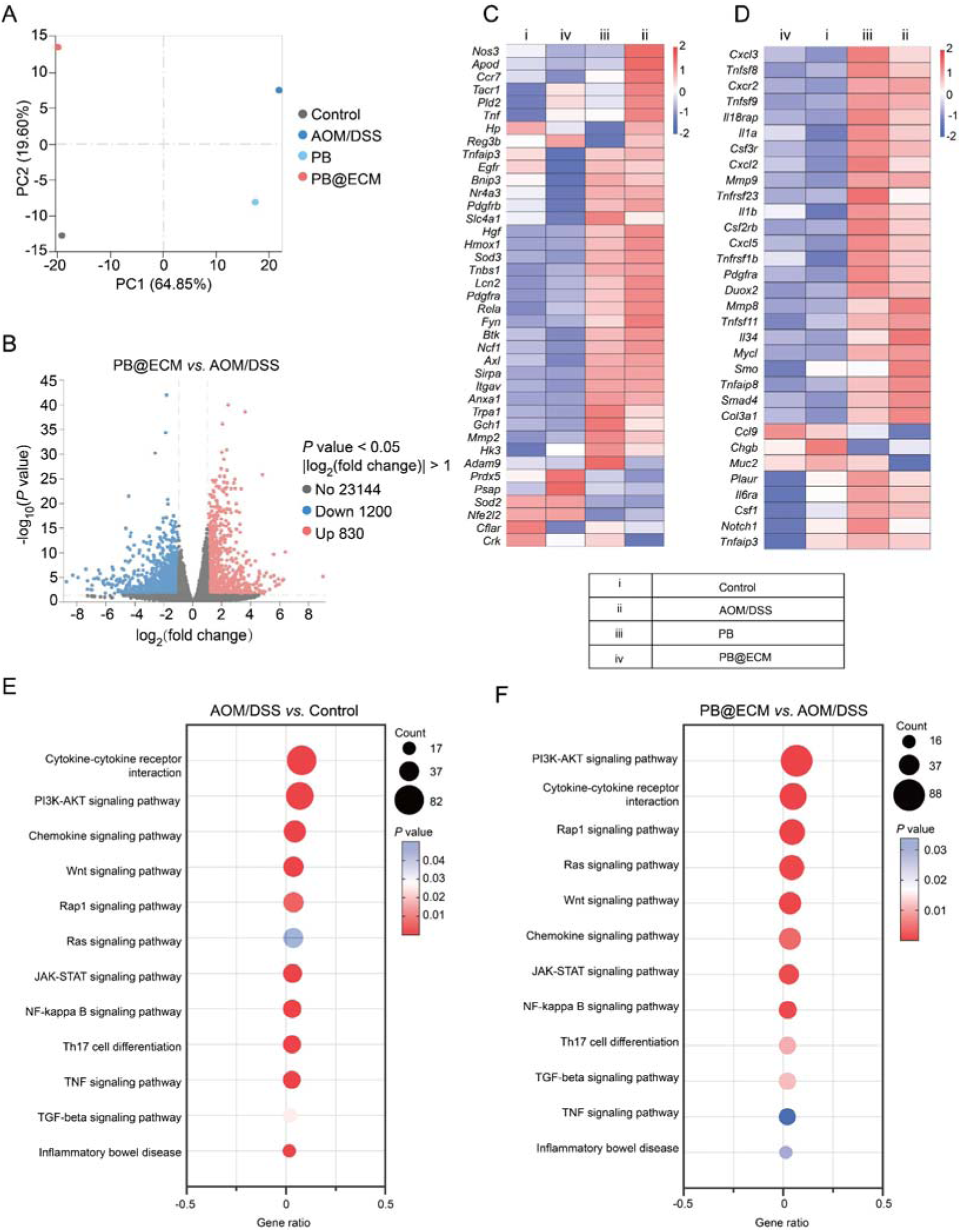
Transcriptome sequencing analysis of treatment with PB@ECM. **A**, Principal component analysis showed diversity of different groups (n = 3). **B**, Volcano plots of differential expression analysis (DEGs) in AOM/DSS and PB@ECM groups (|fold change| ≥ 2.0; *P* adjust < 0.05; up-regulated genes: red; down-regulated genes: blue). Heatmap of DEGs related to ROS (**C**) and tumor occurrence (**D**). KEGG pathways enriched up-regulated genes in AOM/DSS group compared to Control group (**E**) and down-regulated genes in PB@ECM-treated group versus AOM/DSS group (**F**). PI3K-AKT, Phosphatidylinositol 3-Kinase/Akt pathway; JAK-STAT, Janus kinase–signal transducers and activators of transcription; Rap1, Ras-proximate-1; Wnt, Wingless/Integration; TNF, Tumor necrosis factor; NF-kappa B, Nuclear factor kappa-light-chain-enhancer of activated B cells.

A heatmap of selected genes revealed a prominent down-regulation of pathways associated with ROS response, inflammatory signaling, and tumorigenesis following PB@ECM treatment (Figure 8C, D). Kyoto Encyclopedia of Genes and Genomes (KEGG) pathway enrichment analysis further demonstrated that PB@ECM treatment significantly suppressed pathways associated with immune activation and oncogenic pathways, including cytokine-cytokine receptor interaction, chemokine signaling, PI3K-AKT, Rap1, Wnt, JAK-STAT, and NF-κB signaling cascades (Figure 8E, F). Consistently, gene set enrichment analysis (GSEA) demonstrated a down-regulation of tumor-promoting signaling pathways upon PB@ECM treatment, including PI3K-AKT, TGF-β, and TNF (Supplementary Figure 34). GO analysis further indicated that PB@ECM down-regulated genes involved in leukocyte migration and activation, such as those regulating neutrophil chemotaxis, macrophage activation, and general cell migration (Supplementary Figure 35). These transcriptomic changes reflected PB@ECM’s capacity to sequester CXCL2 and thereby limiting recruitment of neutrophil and macrophage, the effect also supported by ELISA measurement and immunofluorescence staining (Figure 7O, P, and Supplementary Figure 28B). Collectively, these findings suggested that PB@ECM conferred therapeutic benefit by dampening inflammation, mitigating oxidative stress, and inhibiting immune cell infiltration, and pro-tumor signaling, ultimately halting the progression of colitis into colorectal cancer.

## Conclusion

Nanozymes have emerged as powerful platforms for biomedical applications owing to their catalytic robustness and multifunctionality^31–33^. Antioxidant nanozymes, in particular, have shown promise in treating inflammation-driven diseases by mitigating excessive ROS^34^, including IBD^35–37^ and cardiovascular diseases^38, 39^. However, chronic inflammatory microenvironments are governed not only by oxidative stress but also by persistent immune cell infiltration and chemokine signaling, which together accelerate tissue damage and malignant transformation. The inability of single-functional nanozymes to simultaneously address these interconnected pathological cues has limited their efficacy in intercepting inflammation-to-cancer progression.^40, 41^.

In this study, we report PB@ECM, a biomimetic nanozyme that integrates catalytic antioxidant activity with targeted chemokine sequestration to reprogram inflammatory immune niches. The PB core efficiently scavenges ROS via SOD- and CAT-like activities, while the genetically engineered macrophage membrane coating selectively neutralizes CXCL2 through overexpressed CXCR2, thereby suppressing excessive neutrophil and macrophage recruitment. By concurrently attenuating oxidative injury and immune chemotaxis, PB@ECM disrupts the self-amplifying inflammatory cascade that drives colitis-associated colorectal cancer. Compared with PB or ECM alone, PB@ECM exhibits synergistic therapeutic efficacy, markedly reducing inflammation, immune infiltration, and tumor burden across multiple stages of disease progression.

Importantly, PB@ECM leverages clinically relevant components, as PB is FDA-approved and cell membranes are inherently biocompatible^42, 43^, underscoring its translational potential. Future studies should focus on optimizing administration routes, validating efficacy and safety in large animal models, and benchmarking against established anti-inflammatory therapies. Overall, this work establishes a dual-action biomimetic nanozyme paradigm for intercepting inflammation-driven carcinogenesis. Beyond colitis-associated colorectal cancer, the modular integration of enzyme-mimicking cores with engineered membranes offers a broadly applicable strategy for reprogramming pathological immune microenvironments and preventing inflammation-associated malignancies.

## Acknowledgments

We thank Mr. Jiang Du for help in BET testing and Dr. Gen Wei for help in EPR testing. This work was supported by the National Natural Science Foundation of China (W2512074 and 22374071), the Key Program of Nanozyme Laboratory in Zhongyuan (NLZ-KP2024NIC06), the Natural Science Foundation of Jiangsu Province (BK20241895), the Jiangsu Provincial Key R&D Program (BE2022836), the State Key Laboratory of Analytical Chemistry for Life Science (5431ZZXM2501), the Fundamental Research Funds for the Central Universities (2025300292), the PAPD Program, Open Funds of NMPA Key Laboratory for Biomedical Optics (20240001), and the International Expansion and Enhancement Program by Nanjing University International Affairs Office.

