## supporting information for "Genetically Engineered Biomimetic Nanozymes Reprogram Immune Niches to Intercept Colitis-Carcinoma Transition"

### Materials

K_3_[Fe(CN)_6_] was obtained from Xi Long Scientific Co, Ltd. (China). Polyvinylpyrrolidone (PVP, Mw~55000) was purchased from Sigma-Aldrich (USA). HCl was purchased from Nanjing Chemical Reagent Co, Ltd. Trypsin, fetal bovine serum (FBS), Dulbecco's modified eagle medium (DMEM), and antibiotics (Penicillin-Streptomycin Solution) were obtained from BioChannel Biotechnology Co, Ltd. 5,5-dimethyl-1-pyrroline N-oxide (DMPO) was purchased from Dojindo (Japan). 30% H_2_O_2_ was obtained from Yong Hua Chemical Co., Ltd. Xanthine oxidase (XO) was purchased from Sigma-Aldrich. Xanthine (X) was obtained from Shanghai Yuanye Bio-Technology Co., Ltd. Hydroethidine (HE) was purchased from Shanghai Macklin Biochemical Technology Co., Ltd. 2′,7′-dichlorofluorescin diacetate (DCFH-DA) was purchased from MedChemExpress. SB225002 was obtained from Selleck Chemicals. Dextran sodium sulfate (DSS) and azoxymethane (AOM) were obtained from MP Biomedicals (USA). Ultrapure water was produced using a Milli-Q water purification system.

### Characterization

High-angle annular dark-field scanning TEM (HAADF-STEM) and their corresponding energy-dispersive spectroscopy (EDS) elemental mappings were performed on an FEI Titan3 G2 60-300 TEM equipped with an acceleration voltage of 300 kV and an EDS detector, respectively. Transmission electron microscopy (TEM) imaging was performed on a JEM-1400 microscope (JEOL, Japan) at an acceleration voltage of 80 kV. Powder X-ray diffraction (XRD) patterns were measured at a scanning rate of 5°/min using a diffractometer (Rigaku Ultima III, Japan or Bruker D8 Advance, Germany) with a Cu Kα radiation. Ultraviolet-visible absorption spectra were measured on a Spectra Max M2e microplate reader (Molecular Devices, USA). Electron paramagnetic resonance (EPR) spectra were recorded on a Bruker A300 spectrometer (X-band). Scanning electron microscopy (SEM) measurements were performed on a microscope at 15 kV (Merlin Compact, Carl Zeiss, Germany). X-ray photoelectron spectra (XPS) were collected using a Nexsa G2 (Thermo Scientific, USA). Hydrodynamic size and Zeta potentials were measured on a Nano sizer ZS90 (Malvern). The Brunauer–Emmett–Teller (BET) surface area data were obtained at 77 K on a Kubo X1000 (Biaode, China) using N_2_ adsorption-desorption isotherms. Images of H&E-stained tissues were photographed using a KF-PRO-005 scanner (KFBIO). Gene heatmap, KEGG pathway, and gene ontology (GO) analyses were performed *via* standard transcriptome analysis.

**Synthesis of PB**

The synthesis of PB was performed as previously described ^1^. K_3_[Fe(CN)_6_] (0.8 mM) was dissolved in 80 mL water, and then 6 g polyvinylpyrrolidone (PVP, Mw~55000) was added under magnetic stirring to obtain a clear solution. Next, HCl (0.01 M, 80 mL) was mixed with the above clear solution under magnetic stirring. The mixture was then incubated at 80 °C for 20 h. After the reaction, the precipitates were collected by centrifugation and washed three times each with ethanol and water. The synthesized PB samples were freeze-dried and stored for further use.

**Synthesis of PB@ECM**

0.5 mL of PB solution (1 mg/mL) was mixed with 0.5 mL of ECM (~6 × 10^6^ cells/mL) in PBS (pH 7.4), and then co-extruded through a 0.8 μm polycarbonate membrane with a miniextruder (Avanti Polar Lipids, USA) for 13 times to get the PB@ECM nanocomposites. The samples were stored at -80 ℃ for further use.

### Synthesis of FITC-PB@DiD-ECM for cell uptake and *in vivo* targeting

K_3_[Fe(CN)_6_] (696 mg) and PVP (8 g) were dissolved in hydrochloric acid solution (1 M) under magnetic stirring to obtain a clear yellow solution. Then the yellow solution was placed in an oven at 80 ℃ for 20 h to obtain a blue solution. The blue solution was centrifuged and washed several times with water to obtain porous Prussian blue (PB).

Certain amounts of FITC (1 mg/mL) were added to porous PB solution. After magnetic stirring for 12 h in the dark, the mixed solution was centrifugated (11,000 rpm, 10 min) and washed 5 times with water until the supernatant was free of FITC. Then, 0.5 mL of FITC-labeled PB solution (1 mg/mL) was mixed with 0.5 mL of DiD-labeled ECM (ECM, engineered macrophage membranes; ~6 × 10^6^ cells/mL) in PBS (pH 7.4). The mixture was then co-extruded through a 0.8 μm polycarbonate membrane with a mini-extruder (Avanti Polar Lipids, USA) for 13 times to get the FITC-PB@DiD-ECM nanocomposites. The samples were stored at −80 ℃ for further use.

### Biological TEM characterization of ECM, PB, and PB@ECM

For TEM analysis, a drop of the ECM or nanoparticle suspension was applied to a copper grid and incubated for 5 min, followed by washing with deionized water. Subsequently, the grid was negatively stained with a drop of 1% uranyl acetate for 8 min, and excess stain was removed using absorbent paper. The sample was visualized using a TEM (JEM-1400, JEOL).

### Coomassie blue staining

The proteins of cell membrane, PB@ECM, and PB were separated by 10% glycine SDS-PAGE. The resulting gels were stained with Coomassie brilliant blue (Meryer (Shanghai) Chemical Technology Co. Ltd., China).

### Measurement of SOD-like activity

HE, an O_2_^•^^−^ fluorescence probe, was used to assess the amount of O_2_^•−^. The SOD-like activity of PB and PB@ECM was evaluated by measuring the amount of O_2_^•−^ with the HE probe. Typically, different concentrations of PB and PB@ECM, xanthine (0.5 mM) and xanthine oxidase (0.05 U/mL), were mixed in Tris-HCl buffer (0.1 M, pH 7.4) at 37 °C in 30 min. HE (0.5 mg/mL) was added to the solution for another 30 min reaction. Then, the fluorescence spectra were recorded, with excitation and emission wavelengths of 470 and 610 nm, respectively.

The SOD-like activity of PB and PB@ECM was also assessed by using a SOD assay kit. First, 20 μL of the sample was mixed with 200 μL of 2-(4-iodophenyl)-3-(4-nitrophenyl)-5-(2,4- disulfophenyl)-2H-tetrazolium, monosodium salt. Next, 20 μL of the enzyme working solution was added to the mixture and cocultured at 37 °C for 20 min. Finally, the absorbance at 450 nm was recorded by using a microplate reader.

### Measurement of CAT-like activity

In a typical measurement^2^, the reaction system in the 96-well plate contained 5 μL of PB or PB@ECM, 90 μL of Tris buffer (10 mM, pH 8.5), 100 μL of dopamine (2 mg/mL), and 5 μL of H_2_O_2_ (0.8 M). After adding the solution into the plate, the plate was quickly transferred to an anaerobic equipment and incubated at 37 °C for 30 min. Finally, the absorbance at 405 nm was recorded by using a microplate spectrophotometer.

### Measurement of hydroxyl radical scavenging activity via EPR

EPR spectra were used to measure the hydroxyl radicals (^•^OH) scavenging activity. Hydroxyl radicals were generated by the reaction of H_2_O_2_ and Fe^2+^, which can be captured by DMPO. The sample contained a final concentration of H_2_O_2_ (0.5 mM), Fe^2+^ (5 μM), nanozyme (0.1 mg/mL), and DMPO (22.5 mg/mL) in PBS buffer (pH = 7.4). After adding Fe^2+^ to start the reaction for 2 min, the nanozyme was added to react for 60 s, and the EPR spectra were immediately recorded.

**Cell culture**

The mouse macrophage cell line RAW264.7 and the human colon cancer cell line Caco-2 were purchased from the Cell Bank of Type Culture Collection of Chinese Academy of Science (Shanghai, China). The RAW264.7 cells and Caco-2 cells were maintained in Dulbecco's modified eagle medium (DMEM) supplemented with 10% (v/v) fetal bovine serum (FBS) and 1% (v/v) antibiotics. All the cells were maintained at 37 °C in a humidified incubator with 5% CO_2_.

**Genetically engineered macrophages**

A lentivirus vector encoding CXCR2 fused at the C-terminal region with a ZsGreen-tag (pHBLV-m-CXCR2-3xflag-ZsGreen-PURO, Supplementary Table 3) and the lentivirus were obtained from Hanbio Biotechnology. RAW264.7 cells were infected with the lentivirus and subsequently incubated in DMEM supplemented with 10 μg/mL puromycin to get a cell line stably expressing m-CXCR2-ZsGreen. The cells stably expressing m-CXCR2-ZsGreen were named CXCR2-RAW cells. CXCR2-RAW cells were cultured in DMEM containing 10% FBS, 1% antibiotics, and 10 μg/mL puromycin.

**Cell membrane extraction**

Cell membranes extraction from CXCR2-RAW cells were performed as previously described^3^. Briefly, cells were washed with 1 × PBS (pH 7.4), detached using 1 × PBS (pH 7.4), and collected by centrifugation at 1.5 × 10^3^ rpm for 5 min. The precipitated cell pellets were repeatedly washed with 1 × PBS (pH 7.4) and resuspended in 1 mL of lysing buffer (20 mmol/L Tris-HCl, 10 mmol/L KCl, 2 mmol/L MgCl_2_, and 1 mini tablet of protease inhibitor without EDTA). After 5 min incubation, the suspension was centrifuged at 6 × 10^3^ rpm for 5 min to collect the supernatant. Then, 250 μL of lysing buffer was added to resuspend the precipitation. The suspension was centrifuged at 6 × 10^3^ rpm for 6 min to obtain additional supernatant. The combined supernatants were centrifuged at 2 × 10^4^ rpm for 25 min in an ultracentrifuge (Beckman Coulter, USA) and then supernatants were centrifuged at 1 × 10^5^ rpm for 35 min to obtain the cell membrane. The membrane samples were stored at −80 ℃ for further use.

**Study approval**

Collection and measurement of human samples were approved by the Jinling Hospital (2023DZKY-001-01) and informed consent was obtained from all participants.

**Animals**

Male C57BL/6 mice (20-22 g) were purchased from Beijing Vital River Laboratory Animal Technology Co., Ltd. All mice were maintained in specific pathogen free conditions, with an ambient temperature of 24 ± 2 °C, air humidity of 40-70%, and a 12 h dark/12 h light cycle. All animal procedures were performed under the guidelines approved by the Institutional Animal Care and Use Committee of Science and Technology Ethics Committee of Nanjing University (IACUC-2309002).

For the colitis model, mice were housed in groups of five mice per cage and acclimatized for 1 week before inclusion in the study. Mice received 2.5% (w/v) dextran sodium sulfate (DSS) supplemented in the sterilized drinking water for 7 days, followed by water. Healthy control mice received only water. Then, PBS, 200 μL of ECM solution, 10 mg/kg of PB and PB@ECM were administered *via* intravenous injection on predetermined days. Changes in bodyweight were assessed daily over the 11-day experimental period. On the last day of the experiment, mice were sacrificed, and the entire colon and other organs were excised. Colon length was measured and gently washed with physiological saline. Then, two pieces of the distal sections (0.5 cm in length) were used for histological assessment and immunofluorescence staining.

For the colitis associated colon cancer model, mice were housed in groups of five mice per cage and acclimatized for 1 week before inclusion in the study. Mice were intraperitoneally (*i.p.*) injected with 10 mg/kg azoxymethane (AOM) 7 days prior to the first 2.5% DSS-induced colitis period. The early, intermediate, late stages colitis periods were induced by giving 2.5% DSS in drinking water for 7 days, followed by 14 days of water. Healthy control mice received only water. Changes in bodyweight were assessed every couple of days over the experimental period. Mice were sacrificed and the entire colon and other organs were excised at the end of each water. Colon length was measured and gently washed with physiological saline. Then, the distal part of the 3 cm colon was made into Switzerland roll. These Switzerland rolls were used for histological assessment and immunofluorescence staining.

**RT-qPCR**

Total RNA was isolated from RAW264.7 and CXCR2-RAW264.7 cells using TRIzol reagent and reverse-transcribed into cDNA using a cDNA reverse transcription kit (Servicebio, no. G3337). The relative mRNA levels of genes were examined via RT-qPCR assay using a StepOne™ Real-Time PCR System (Thermo Fisher, USA). *β-Actin* was used as an internal control. Primer sequences are listed in Supplementary Table 4.

**Western blot analysis**

RAW264.7 cells, CXCR2-RAW cells, cell membranes, and PB@ECM were lysed using IP Lysis Buffer containing protease inhibitor cocktail (1:100 dilution) and phosphatase inhibitors (1:50 dilution) (Beyotime Biotechnology, Shanghai, China) on ice for 30 min. The proteins were isolated using 10% glycine SDS-PAGE and transferred to PVDF membranes. Proteins were detected by incubation with CXCR2 antibodies at 4°C overnight, followed by horseradish peroxidase (HRP) goat anti-rabbit secondary antibodies (Beyotime Biotechnology, Shanghai, China) for 1 hour. The blots were visualized using an enhanced chemiluminescence (ECL) kit (Servicebio, no. G2014) on a Tanon 5200 Multi Chemiluminescent Imaging System (Tanon, China). The information of antibodies is shown in Supplementary Table 5.

**Immunofluorescence analysis**

To detect the neutrophil and macrophage levels in colon tissues, paraffin-embedded colon sections were deparaffinized and hydrated. The sections were boiled in citrate buffer (Servicebio, China) for antigen recovery. Then, the sections were blocked with 5% Bovine serum albumin (BSA) for 1 h and incubated with primary antibodies against F4/80 and Ly-6G at 4 °C overnight. Fluorescence labeled secondary antibodies were applied at room temperature for 1 h. Hoechst (Beyotime) was used to label the nucleus. Immunofluorescence was observed by a laser confocal microscope (Leica SP8 STED 3X, Germany). The information of antibodies used was shown in Supplementary Table 5.

**Cytotoxicity assay of PB and PB@ECM**

Cytotoxicity was evaluated using RAW264.7 cells. RAW264.7 cells were incubated with a 96-well plate at 37 °C for 24 h, and the cell number of each well was 8 × 10^3^. Then, different concentrations (5, 10, 20, 50, and 100 μg/mL) of PB and PB@ECM were added to each well plate for another 24 h. Finally, cell viability was evaluated by CCK-8 assay.

The cell viability was calculated using the equation

$\text{V}\text{iability}\text{=}\frac{\text{O}_{\text{t}}\text{-}\text{O}_{\text{b}}}{\text{O}_{\text{c}}}\text{×100\%}$	(1)

where O_t_ is the absorbance of the testing samples, O_b_ is the background absorbance of the materials, and O_c_ is the absorbance of the control groups.

**ROS-scavenging activity of PB and PB@ECM**

The intracellular ROS level was monitored using DCFH-DA as a fluorescence probe through flow cytometry and laser scanning confocal microscopy. DCFH-DA has excitation and emission wavelengths at 488 and 525 nm, respectively. Specifically, RAW264.7 cells were incubated with a 6-well plate at 37 °C for 24 h, and the cell number of each well was 1 × 10^5^. Then, different concentrations (50 and 100 μg/mL) of PB and PB@ECM were added to each well plate for another 2 h. After washing three times with PBS (pH 7.4), 150 μM H_2_O_2_ was added and incubated for 30 min to stimulate cells to produce intracellular ROS. After that, DCFH-DA (0.01 mM) in DMEM was added to each well plate and incubated for 30 min, followed by washing. Cells were collected and analyzed using a flow cytometer (CytoFLEX, Beckman Coulter) and a fluorescence microscope (Leica DMi8, Germany). FlowJo v.10 was used for data analysis.

**Intracellular uptake assay**

RAW264.7 and Caco-2 cells were seeded onto round coverslips at 5 × 10^3^ cells per well in 24-well plates. After 24 h, ECM, PB, and PB@ECM were added. Following another 24 h incubation, the round coverslips were collected and washed three times with PBS (pH 7.4). Hoechst was used to label the nucleus. Fluorescence images were acquired using a laser confocal microscope (Leica SP8 STED 3X, Germany).

**Visualization of PB@ECM internalization in RAW264.7 cells**

RAW264.7 cells were seeded at 1 × 10^4^ cells per well in 6-well plates. After 24 h, the ECM, PB, and PB@ECM were added. After another 24 h, cells were collected and fixed with 2.5% glutaraldehyde for 12 h, followed by 2% osmic acid. Cells were then dehydrated, embedded, sectioned with an ultrathin slicer, and stained with 2% uranyl acetate. The sections were observed using TEM imaging.

**Observation of oxidative stress model**

RAW264.7 cells were seeded at 1 × 10^4^ cells per well in 6-well plates. After 24 h, ECM, PB, and PB@ECM were added. After another 24 h, 150 μM H_2_O_2_ was added and incubated for 1 h to induce oxidative stress. Then cells were collected and imaged using TEM.

**Isolation of mouse bone marrow neutrophils**

Bone marrow neutrophils were extracted from mice using a bone marrow neutrophil isolation kit (Solarbio, China). Briefly, the animals were sacrificed, and the femur and tibia were sterile extracted. The cartilage at both ends was cut to expose the red bone marrow cavity. A 1 mL sterile syringe with medium containing 10% standard FBS was used to flush the marrow cavity to get the bone marrow. A single-cell suspension at a concentration of 2 × 10^8^ - 1 × 10^9^ cells/mL was prepared. The suspension was centrifuged at 1000 g for 20~30 min at room temperature using a horizontal rotor. The lower neutrophils were transferred into a 15 mL clean centrifuge tube and washed with 10 mL PBS, followed by centrifugation at 250 g for 10 min. The supernatant was discarded, and the cell pellet was resuspended in 5 mL PBS, centrifuged at 250 g for another 10 min. The cell pellet was washed once more. The supernatant was discarded, and bone marrow neutrophils were obtained and cultured for follow-up experiments.

**Wound healing assay**

The migratory ability of the cells was evaluated by seeding 1 × 10^6^ cells into a six-well plate and culturing with DMEM for 24 h. Then a wound was made by manually scraping the cell monolayer with a 200 μL pipet tip. Cells were incubated for 48 h with FBS-free medium containing CXCL2, ECM, PB, or PB@ECM. Images were taken at the indicated times with an inverted microscope (Zeiss, Axio Vert.A1).

**Transwell assay**

The migratory ability of the cells was also evaluated using a 24-well transwell plate with a 5.0 µm Pore Polyester Membrane (Corning, no. 3421). The upper chamber contained RAW264.7 cells suspended in 150 μL DMEM with 10 % FBS or bone marrow neutrophil suspended in 150 μL RMPI1640 medium with 10 % FBS. The lower chamber contained 600 μL DMEM or RPMI1640 medium containing SB225002, CXCL2, ECM, PB, or PB@ECM. The transwell chamber was stained with 0.25 % crystal violet after 2 days of incubation at 37 °C. The transwell chambers were photographed with an inverted microscope (Zeiss, Axio Vert).

### Macrophage phenotype identification

RAW264.7 cells were seeded in 6-well plates and cultured for 24 h. The experimental groups were divided into Control, LPS, LPS + PB, LPS + ECM, and LPS + PB@ECM. Cells were treated with LPS (200 ng/mL) and nanozymes (50 μg/mL) for 24 h. After treatment, the cells were collected and labelled with APC-anti-mouse CD86 (Biolegend). The phenotypes of these labelled cells were identified using flow cytometry (CytoFLEX, Beckman). FlowJo v.10 was used for data analysis. The cell supernatant was collected for the subsequent enzyme-linked immunosorbent assay (ELISA) analysis.

### ELISA analysis

RAW264.7 cells were seeded in 6-well plates and cultured for 24 h. The experimental groups were divided into Control, H_2_O_2_, H_2_O_2_ + ECM, H_2_O_2_ + PB, and H_2_O_2_ + PB@ECM. Cells were treated with H_2_O_2_ (200 μM) and nanozymes (50 μg/mL ) for 24 h. After the treatment, the cell supernatants were collected and centrifuged (500 g, 5 min), and immediately frozen at −80 °C. The levels of inflammatory cytokines (IL-1β, IL-6, IFN-γ, and TNF-α) were measured according to the manufacturer’s instruction (Multi Sciences).

For measurement of CXCL2 binding, recombinant mouse CXCL2 (Cisbio Bioassays) was incubated with PB@CM, PB@ECM, or PB@ECM + SB225002 for 2 h at 37 °C. After centrifugation (16100 g, 10 min), CXCL2 concentration in the supernatant was quantified by mouse CXCL2 ELISA kits (ELK Biotechnology).

Mice colon samples were collected. The colon tissue (0.5 g) was placed in a 1.5 mL EP tube without enzyme, and 500 μL of RIPA lysis buffer (Servicebio, no. G2002) was added. The tissue was then homogenized using a SWE-FP high-speed tissue grinder (Servicebio). Afterward, the EP tube was incubated at 4 °C for 40 min, followed by centrifugation at 12000 rpm at 4 °C for 5 min. The supernatant was then collected and immediately frozen at −80 °C for the subsequent ELISA test. The levels of inflammatory cytokines (IL-1β, IL-6, IFN-γ, and TNF-α) and CXCL2 were measured by ELISA analysis.

### RNA-seq of mouse colon tissue

The colon tissues from the late-stage sample of AOM/DSS model were collected. Then the samples were subjected to transcriptome sequencing analysis by Novogene Co., Ltd. (Beijing, China). Whole-transcriptome profiles were generated using a NovaSeq platform (Illumina).

### Statistics and reproducibility

Quantitative data in this study were presented as the mean ± standard deviation (s.d.). All experiments were performed with at least three biological replicates. Statistical analysis between two groups was calculated using two-sided unpaired t-tests. One-way ANOVA followed by post-hoc correction was applied for multiple comparisons. A probability value less than 0.05 (*P* < 0.05) was regarded as statistically significant. NS, no significant difference, **P* < 0.05, ***P*< 0.01, ****P* < 0.001, *****P* < 0.0001. Statistical analyses were performed using GraphPad software.

### Supplementary Figures 1-35

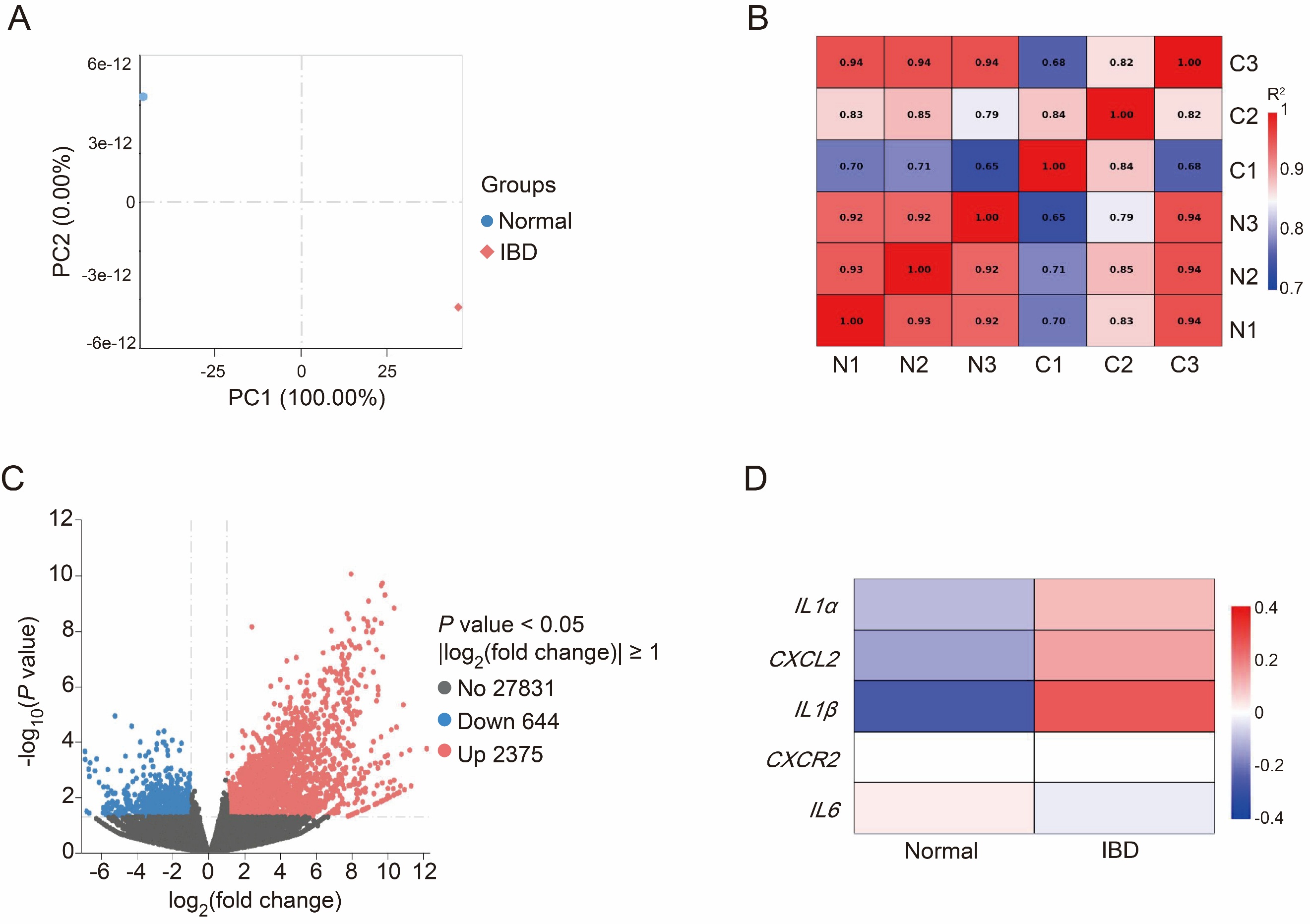

**Supplementary Figure 1.** **A**, Principal component analysis of transcriptional profiles from normal and IBD groups (n = 3). **B**, Correlation heatmap of normal and IBD groups. **C**, Volcano plots revealing differentially expressed genes between normal and IBD groups. **D**, Heatmap showing expression levels of *IL1α*, *CXCL2*, *IL1β*, *CXCR2*, and *IL6* between normal and CAC groups.

We performed transcriptome sequencing on colon tissues from patients with IBD or CAC, and healthy control to reveal their pathological features. The volcano plots showed that 2375 up-regulated and 644 down-regulated genes in IBD (Supplementary Figure 1C), and 956 up-regulated and 1110 down-regulated genes in CAC compared to healthy controls (Supplementary Figure 2C).

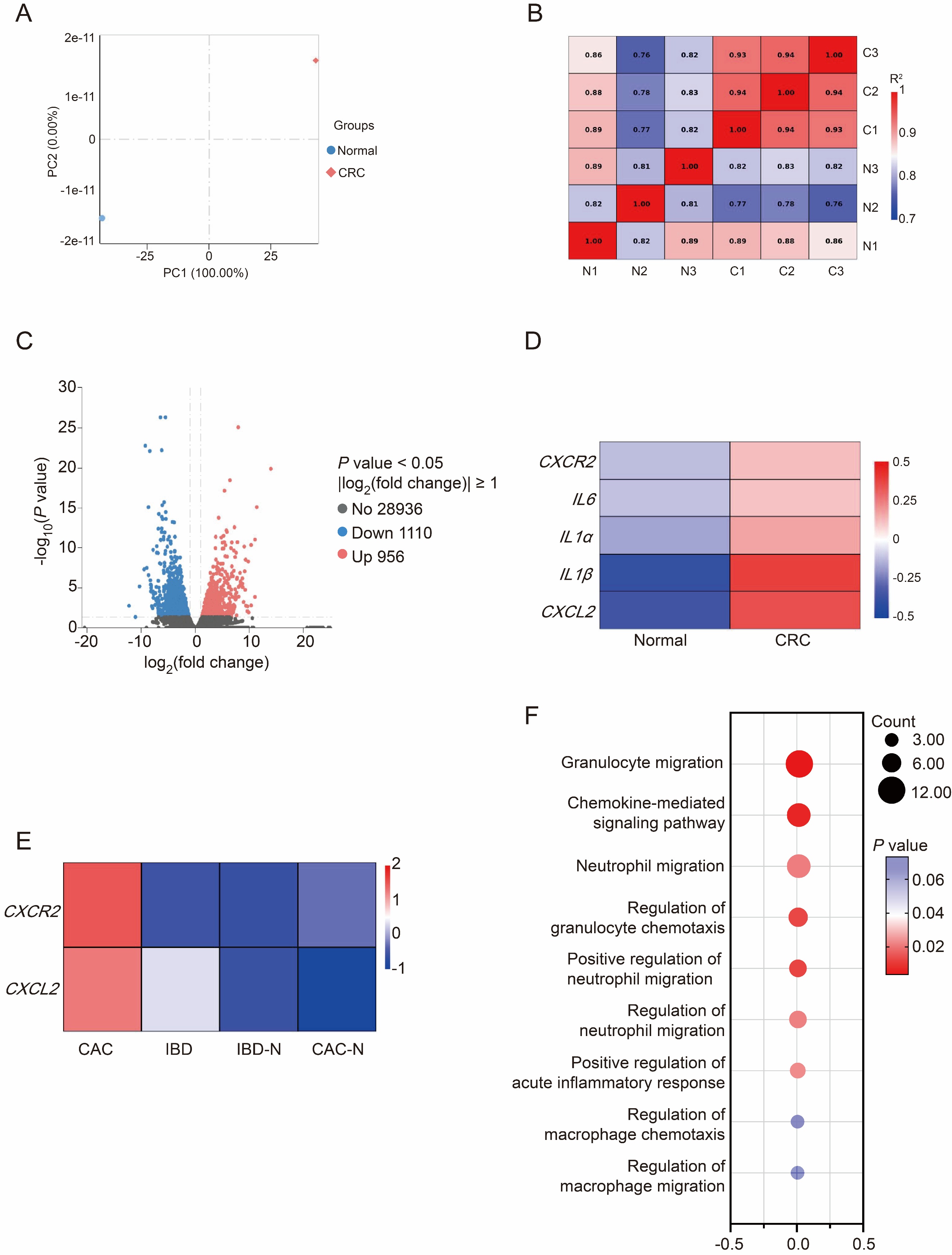

**Supplementary Figure 2.** **A**, Principal component analysis of transcriptional profiles from normal and CAC groups (n = 3). **B**, Correlation heatmap of normal and CAC groups. **C**, Volcano plots revealing differentially expressed genes between normal and CAC groups. **D**, Heatmap showing expression levels of *CXCR2*, *IL6*, *IL1α*, *IL1β*, and *CXCL2* between normal and CAC groups. **E,** Heat map of gene expression of CXCL2 and CXCR2 in patients (IBD-N, health control of IBD; CAC-N, health control of CAC). **F,** Gene Ontology (GO) enrichment analysis of genes up-regulated in CAC patients.

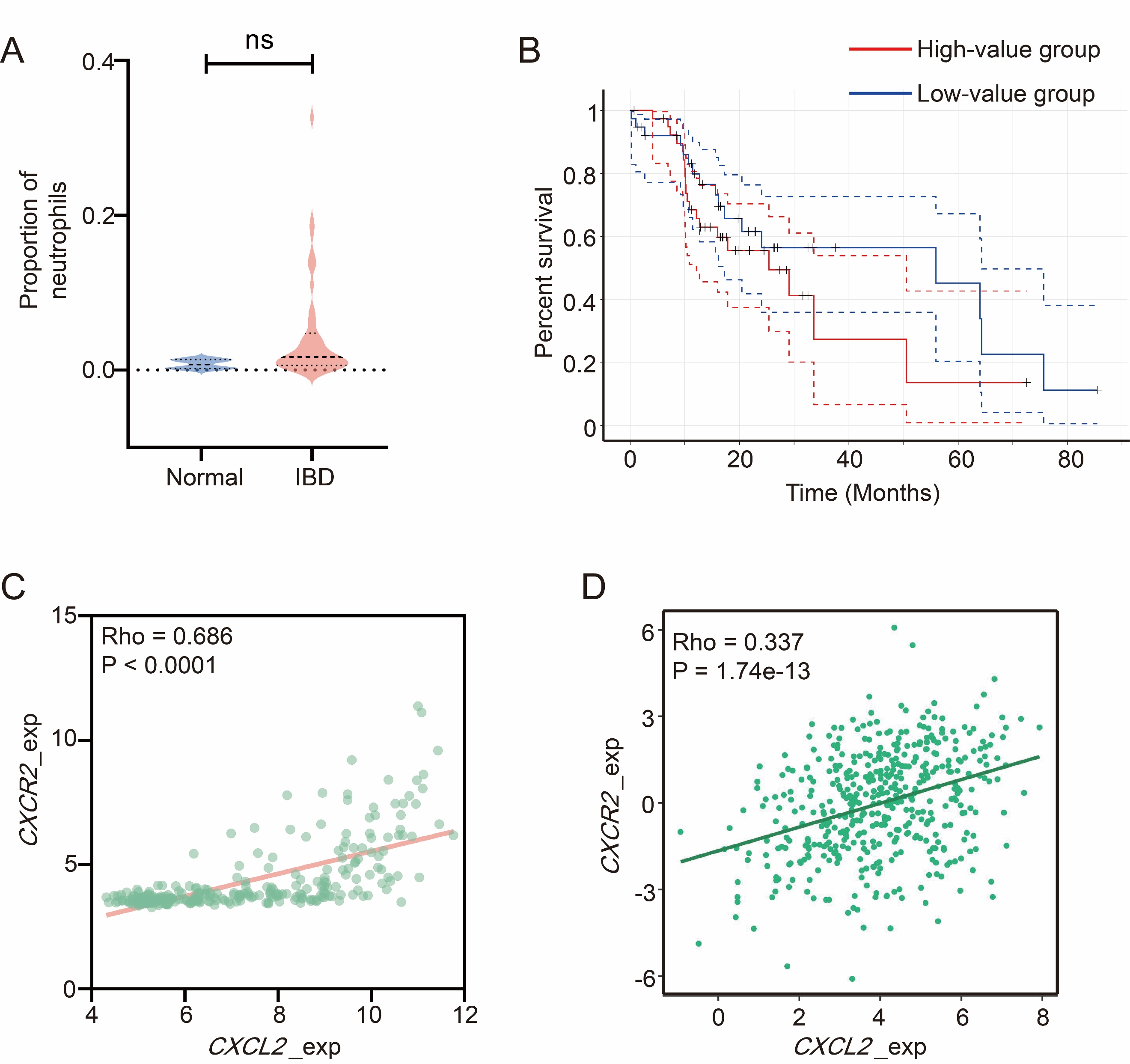

**Supplementary Figure 3.** **A,** Neutrophil infiltration in normal and IBD patient tissues (Normal = 4, IBD = 45). **B**, Kaplan–Meier analysis of relapse-free survival in COAD patients (GEPIA2021). **C**, Correlation between neutrophil levels and *CXCL2* expression in IBD patients (n = 265). **D**, Correlation between *CXCR2* and *CXCL2* expression in COAD patients (n = 459). NS, no significant difference.

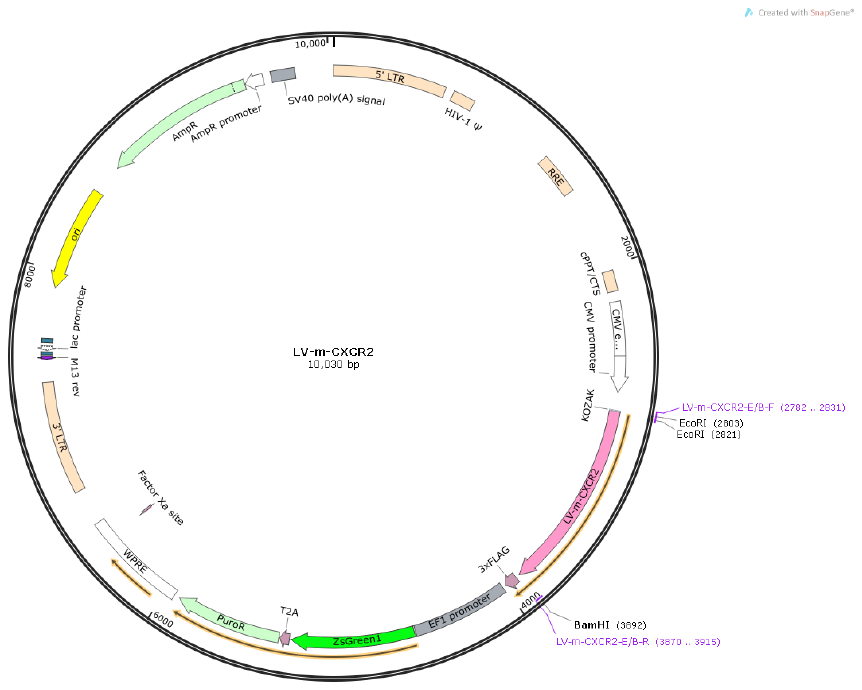

**Supplementary Figure 4.** Plasmid structure diagram, pHBLV-CMV-MCS-3FLAG-EF1-ZsGreen-T2A-PURO. (Plasmid structure diagram from Hanbio Biotechnology)

Zoanthus green fluorescent protein (ZsGreen) and a 3×Flag tag were used as dual reporters to facilitate monoclonal selection and visualization. CXCR2, a G protein-coupled receptor (GPCR), possesses intrinsic transmembrane anchoring capability, enabling its stable display on the engineered cell membranes^4^.

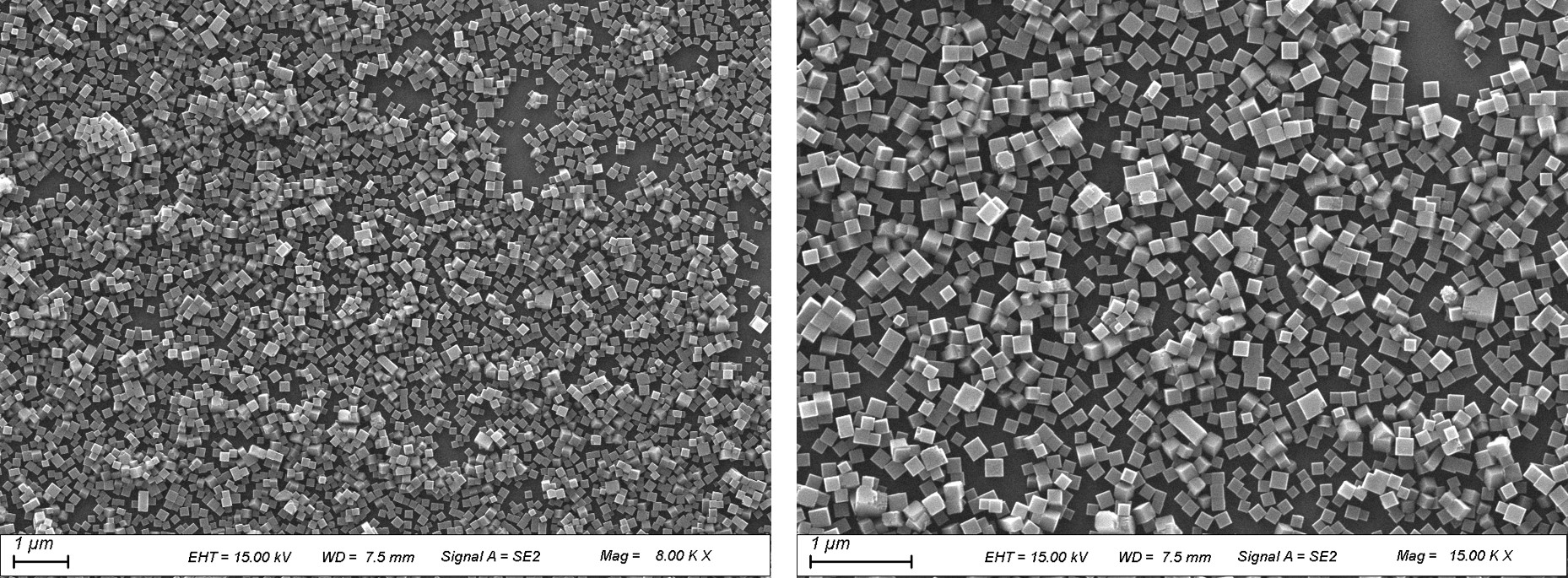

**Supplementary Figure 5.** SEM images of PB.

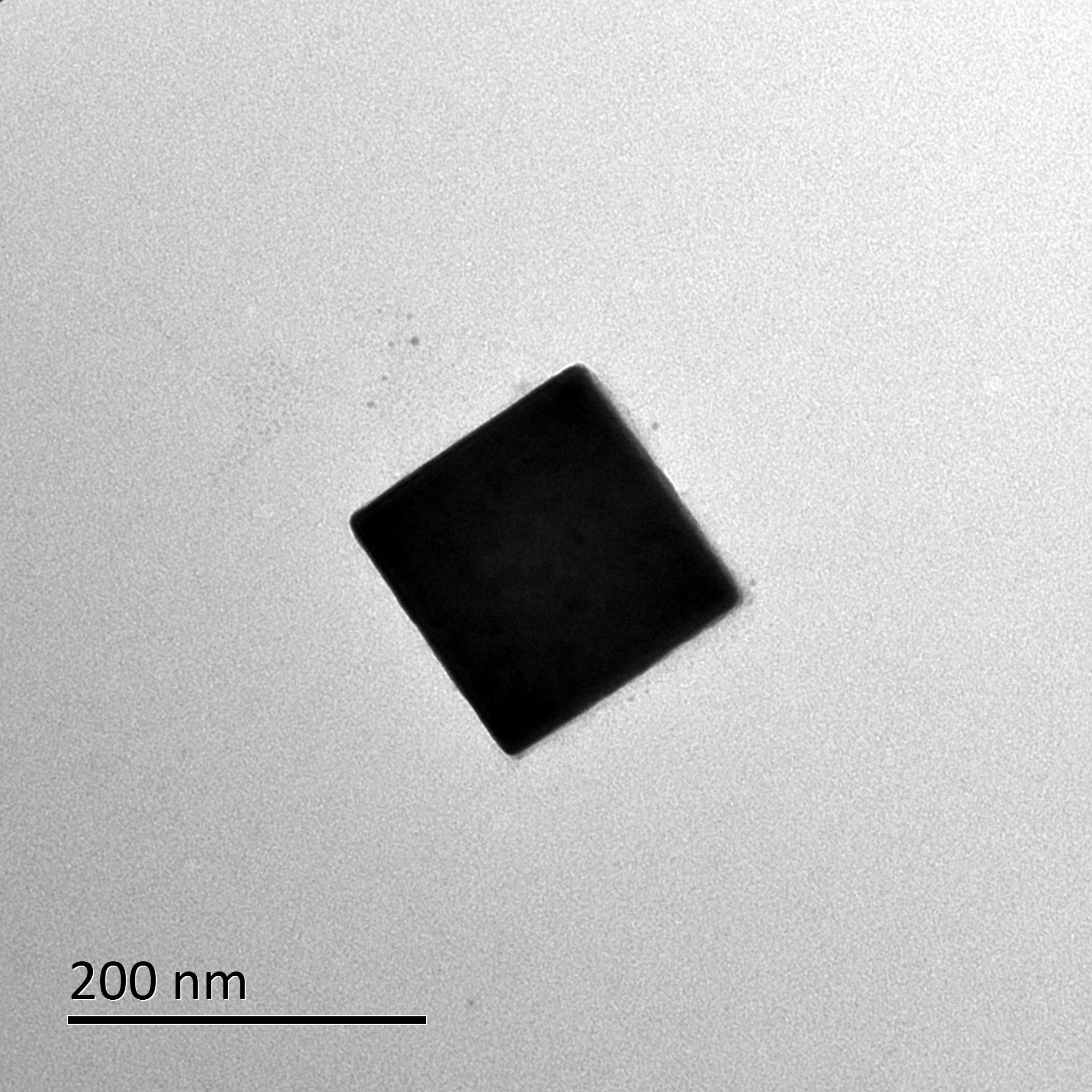

**Supplementary Figure 6.** TEM image of PB.

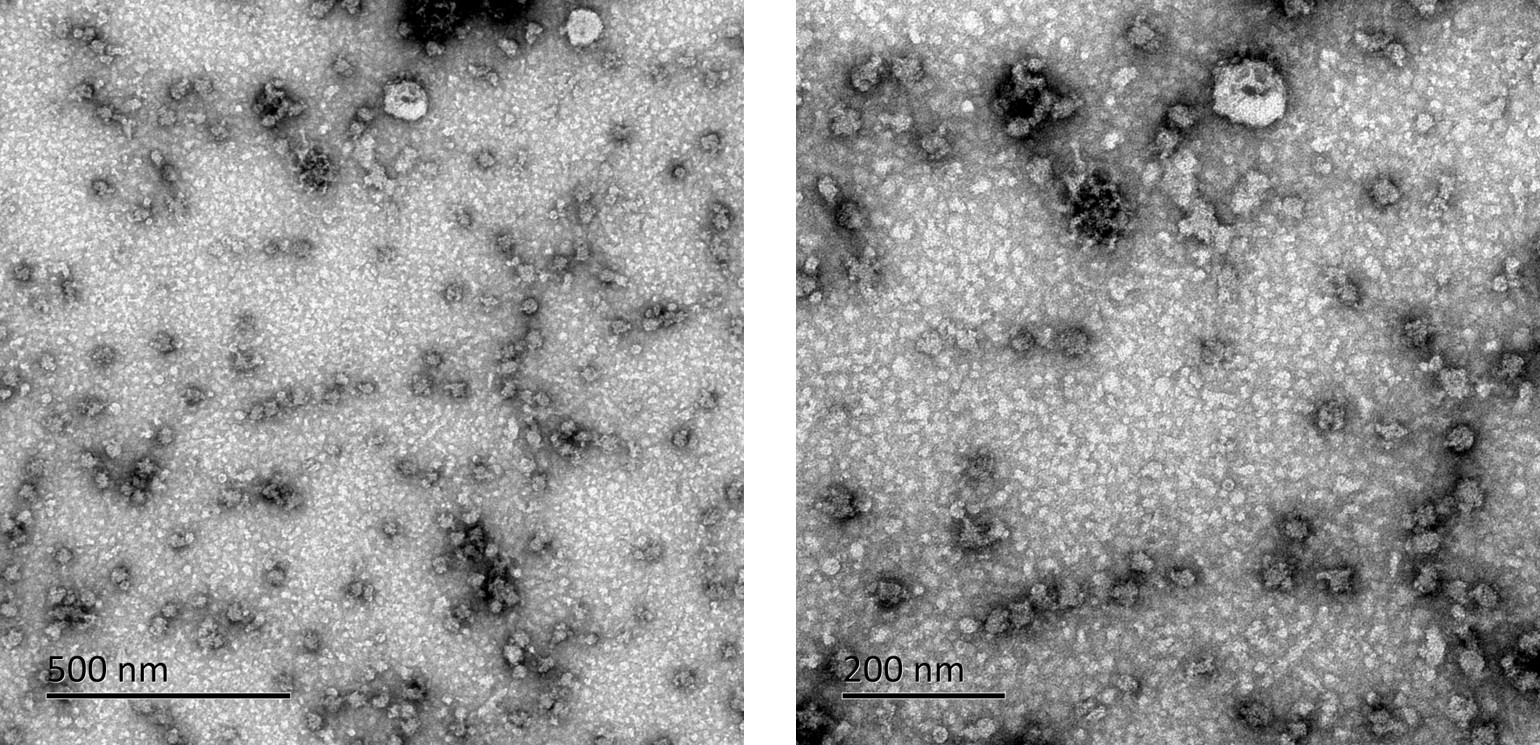

**Supplementary Figure 7.** TEM images of ECM.

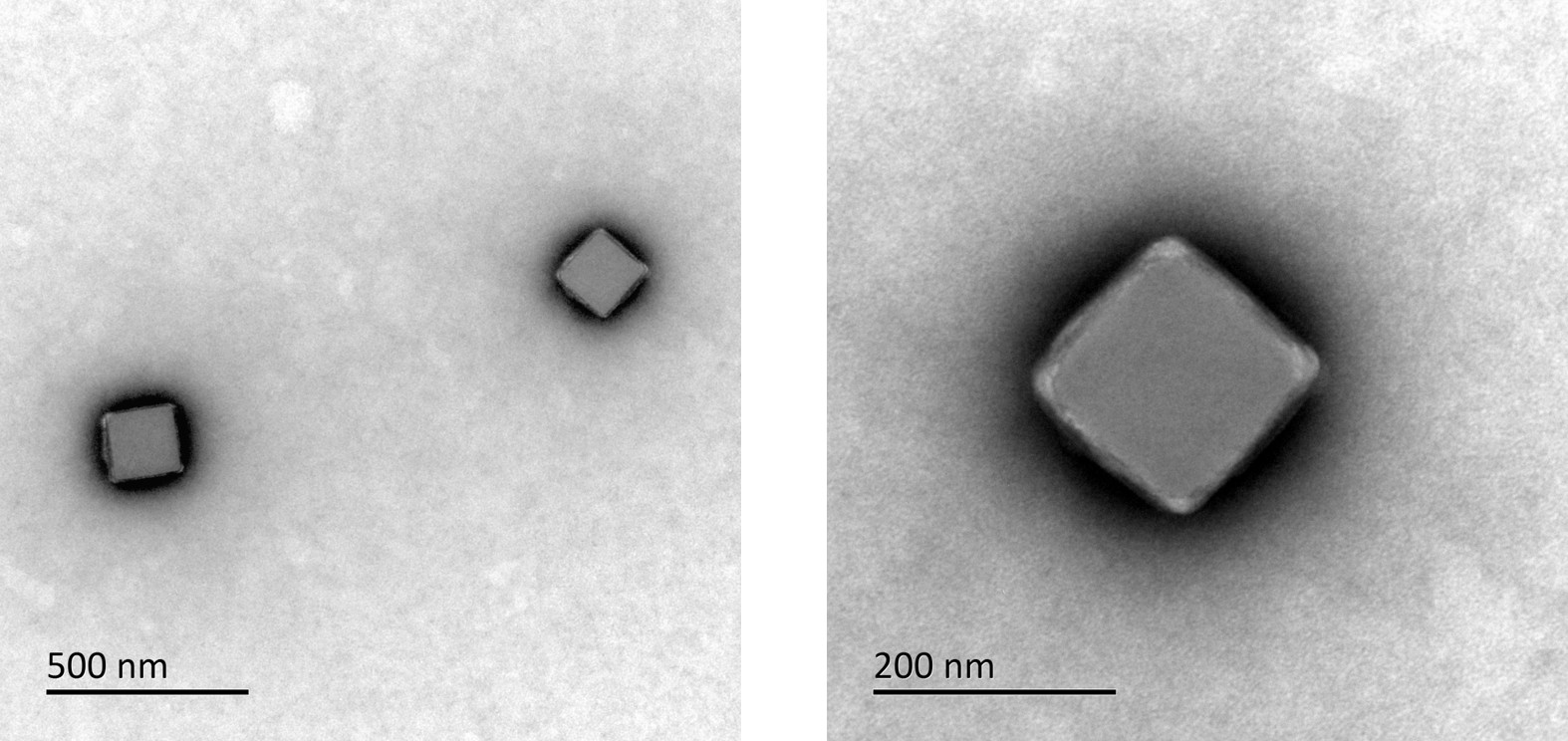

**Supplementary Figure 8.** TEM images of PB@ECM.

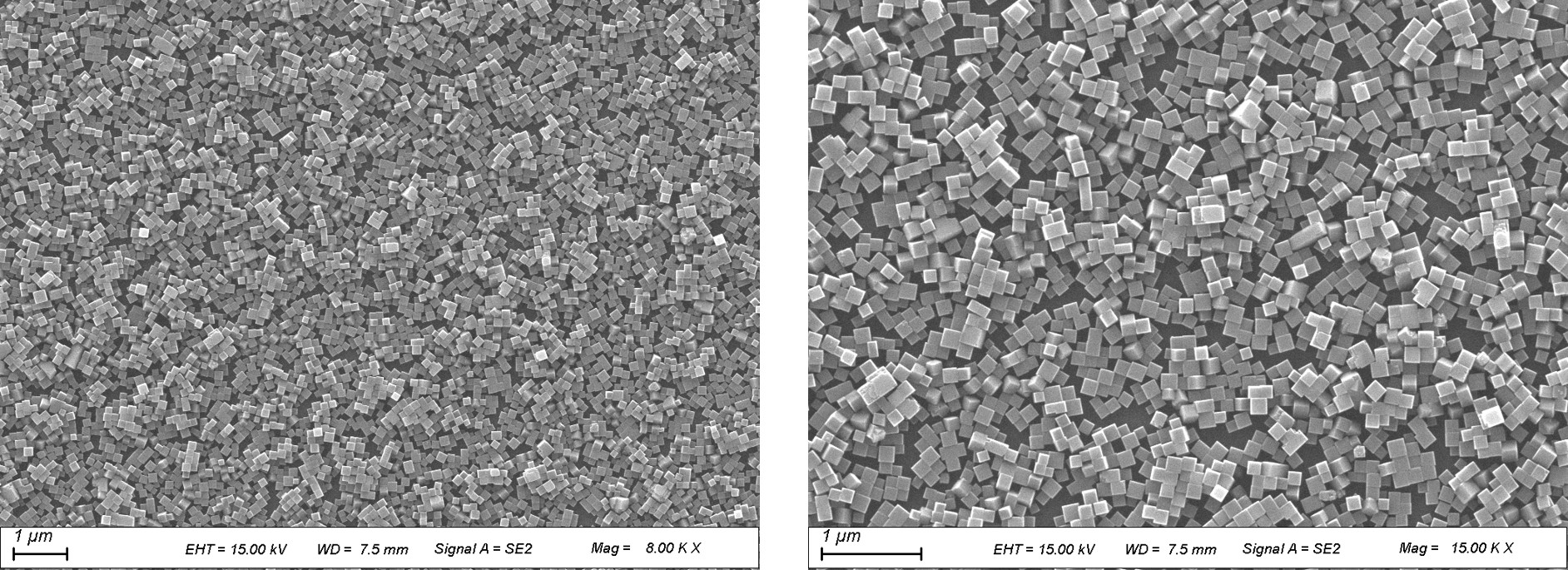

**Supplementary Figure 9.** SEM images of PB@ECM.

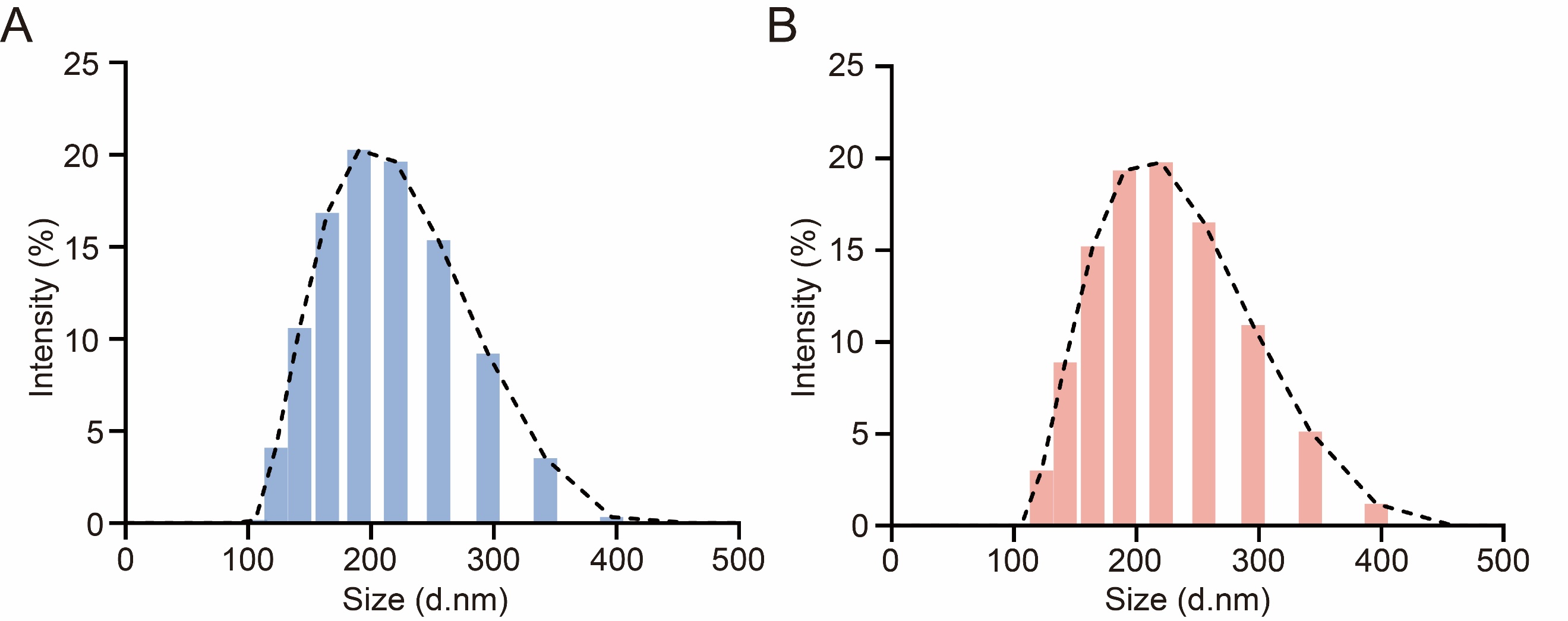

**Supplementary Figure 10.** Size distribution histograms of PB (**A**) and PB@ECM (**B**).

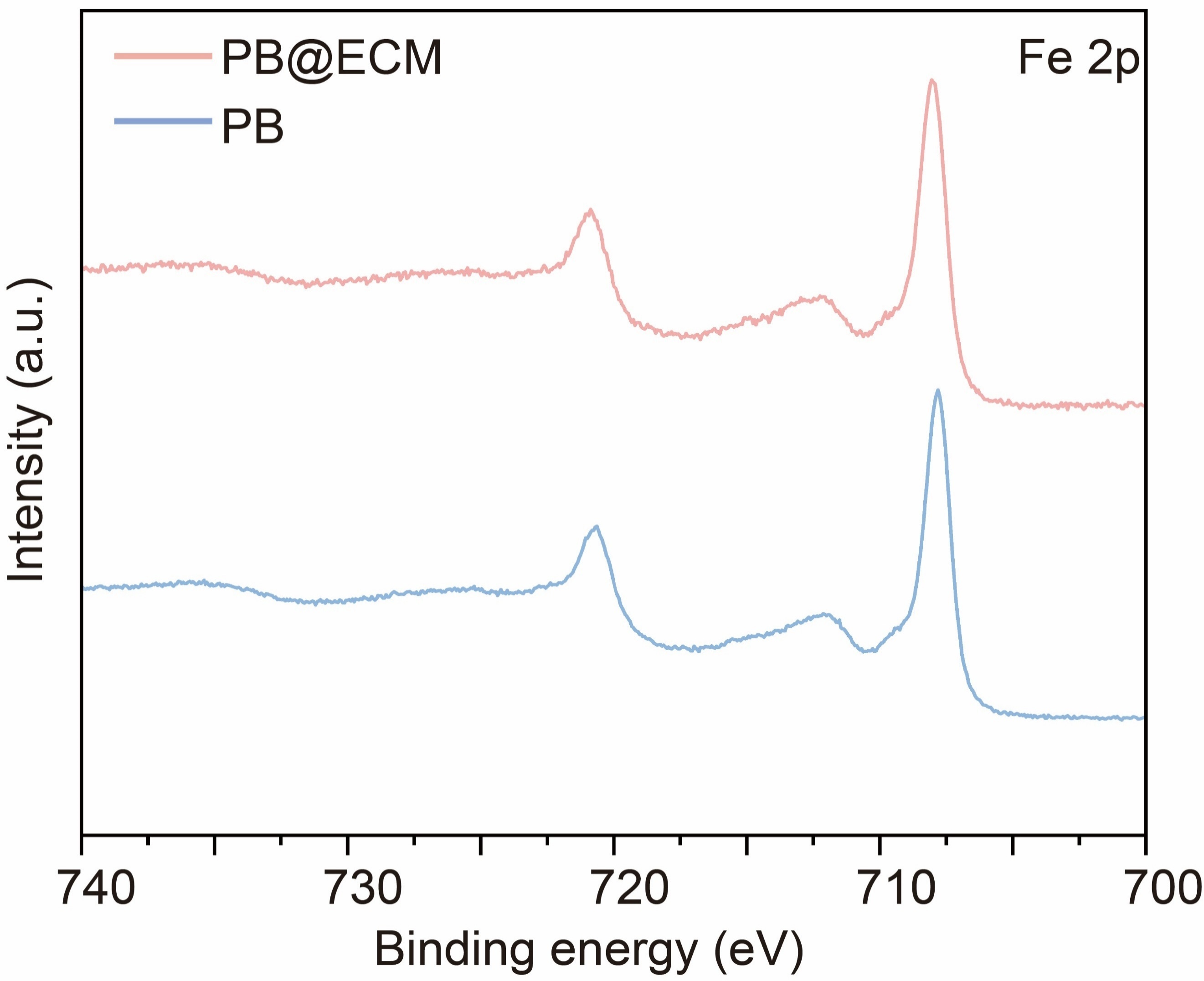

**Supplementary Figure 11.** XPS spectra of Fe 2p of PB and PB@ECM.

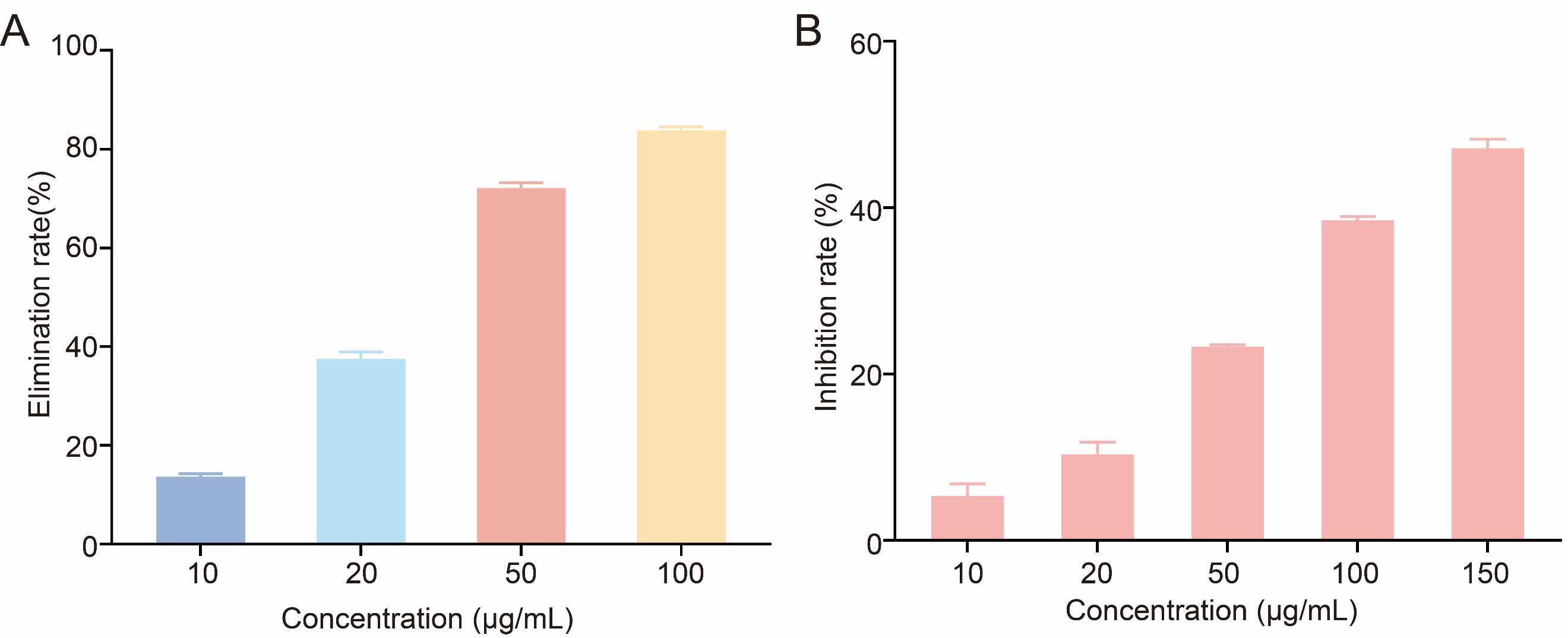

**Supplementary Figure 12. A**, Quantitative analysis of reaction of HE with X and XO in the presence of PB@ECM (n = 3) in Figure 4A. **B**, Dependence between inhibition rate of O_2_^•−^ and concentration of PB@ECM, measured using SOD-kit assay for SOD-like activity (n = 3). As a sensitive O_2_^•−^ indicator, HE could be oxidized by O_2_^•−^ to produce a wide fluorescence spectrum ranging from 550 nm to 650 nm (typically centered at 600 nm).

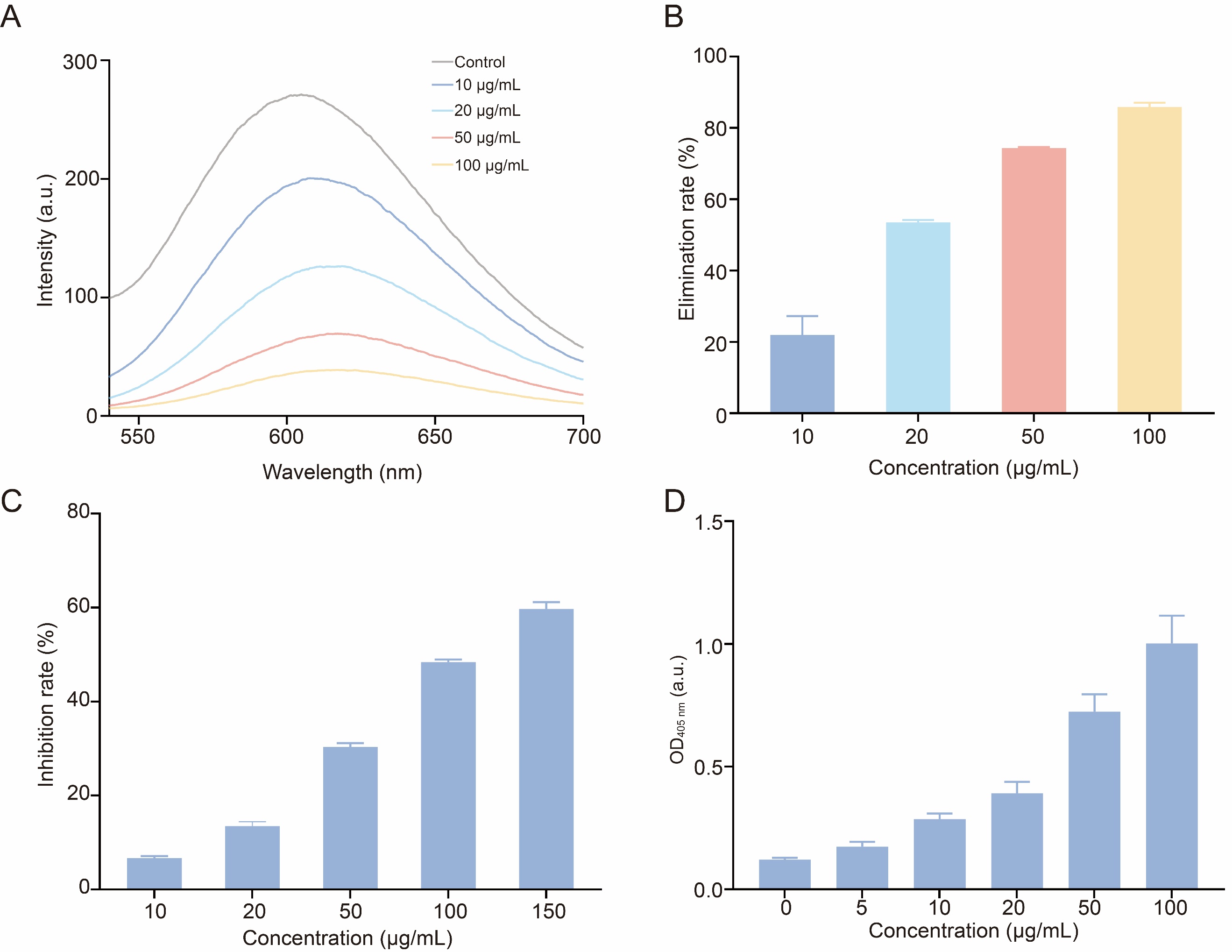

**Supplementary Figure 13.** Fluorescence spectra monitoring reaction of HE with X and XO in the absence and presence of PB (**A**) and corresponding quantitative analysis (**B**) (n = 3). **C**, Dependence between inhibition ratio of O_2_^•−^ and concentration of PB, measured using SOD-kit assay for SOD-like activity (n = 3). **D**, Dependence between OD_405 nm_ and concentration of PB, measured using dopamine-enabled assay for CAT-like activity (n = 3). In this assay, dopamine is oxidized by oxygen to form polydopamine, which exhibits a characteristic absorption peak at 405 nm.

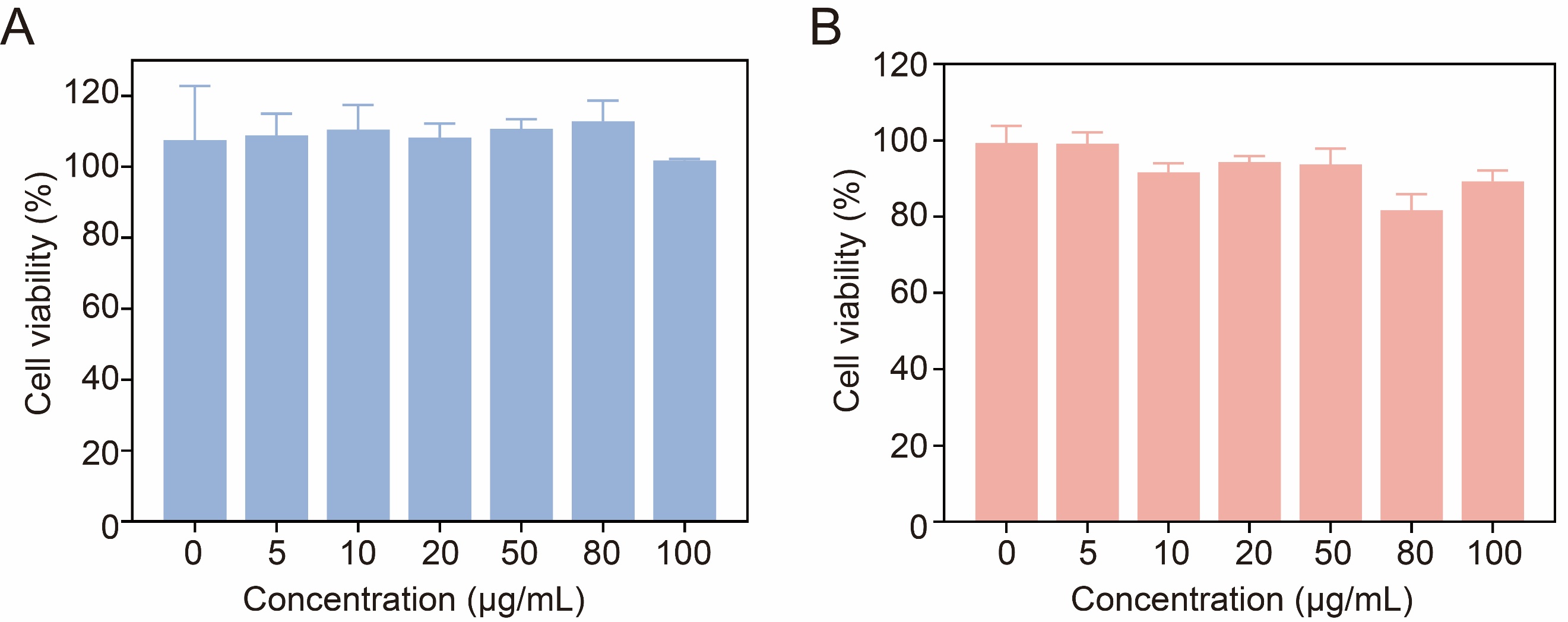

**Supplementary Figure 14.** Cytotoxicity of PB (**A**) and PB@ECM (**B**) evaluated using RAW264.7 cells as a model (n = 3).

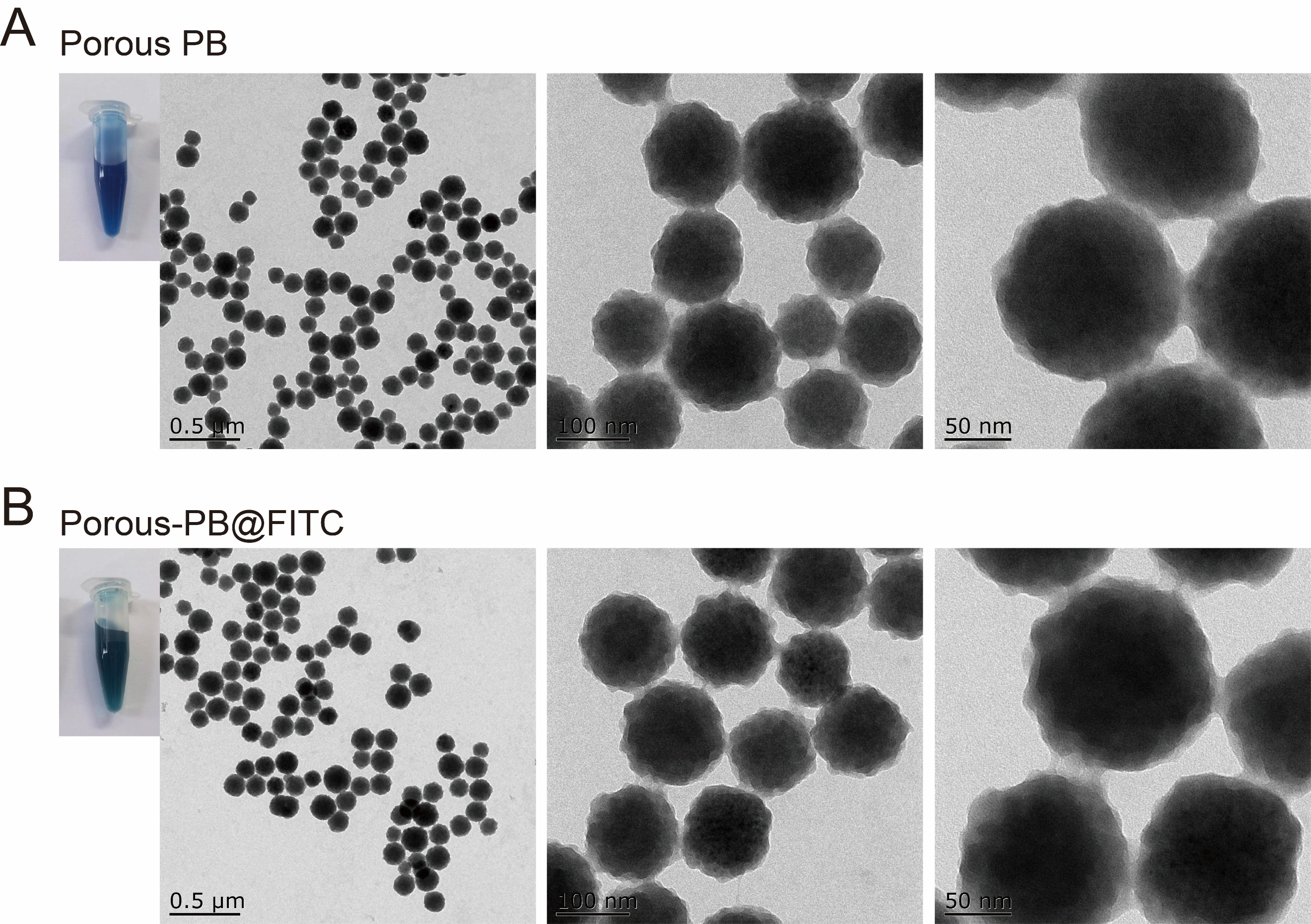

**Supplementary Figure 15.** Digital photos and TEM images of porous PB (**A**) and porous PB@FITC (**B**).

We synthesized porous PB by etching PB with hydrochloric acid to enlarge its pore size. The synthesized porous PB was loaded with FITC to obtain PB@FITC, which was used for *in vivo* targeted validation and cell internalization experiments. In other experiments, unetched PB was utilized.

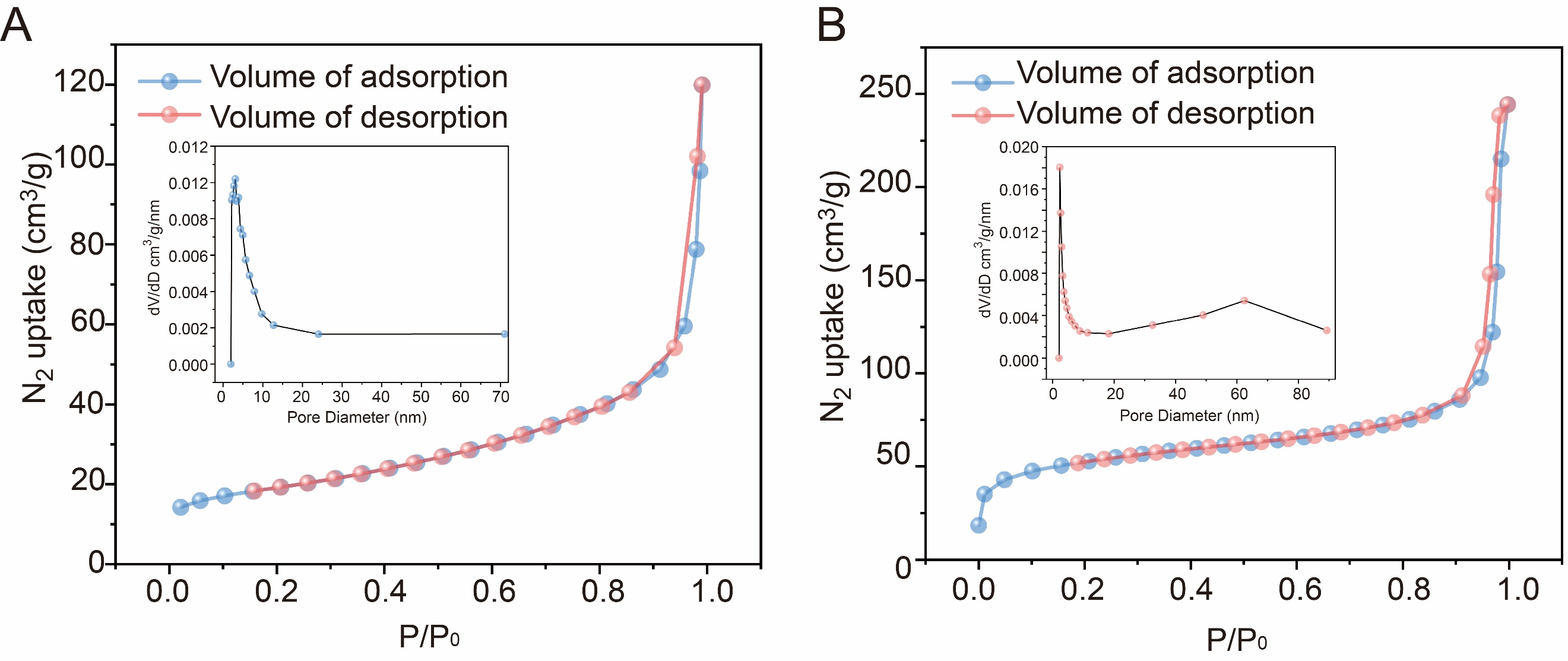

**Supplementary Figure 16.** Nitrogen adsorption–desorption isotherms of PB (**A**) and porous PB (**B**). Insets: corresponding pore size distribution curves.

The Brunauer–Emmett–Teller (BET) analysis was used to evaluate changes in surface area and porosity before and after acid etching. Acid treatment led to an increase in the pore diameter of PB (from ~5.4 nm for PB to ~58.3 nm for porous PB). Mesoporous pore diameters have appeared. Furthermore, the BET surface area increased from 68.2 m^2^/g for PB to189.2 m^2^/g for porous PB, confirming the successful synthesis of porous PB.

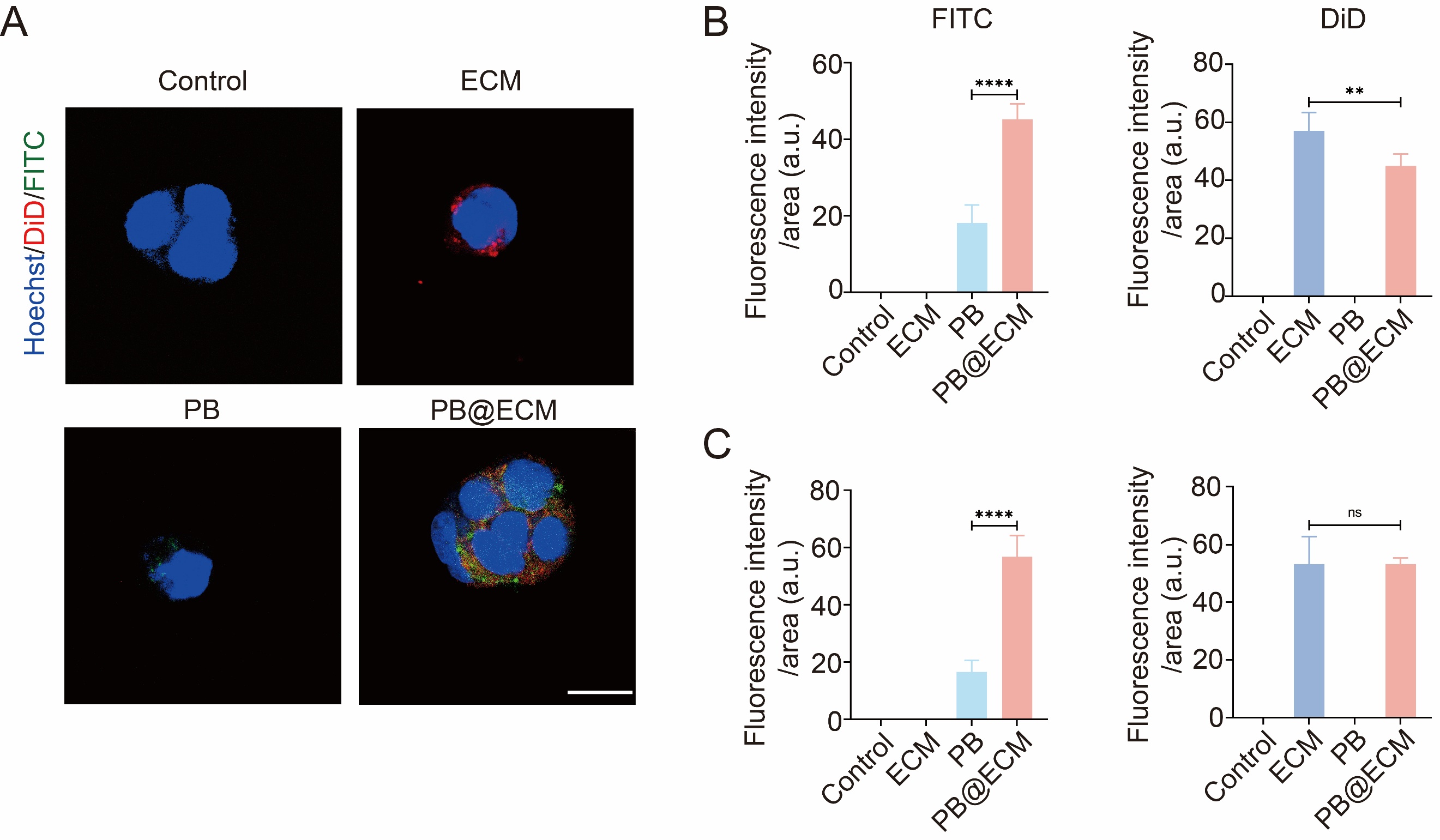

**Supplementary Figure 17. A**, Representative fluorescence images showing uptake of ECM, PB, and PB@ECM by Caco-2 cells. Scale bar, 20 μm. **B**, Quantification of intracellular fluorescence of FITC and DiD in RAW264.7 cells after indicated treatments (n =3). **C**, Quantification of intracellular fluorescence of FITC and DiD in Caco-2 cells after indicated treatments (n =3). FITC, FITC-labeled PB; DiD, DID-ECM; ns, no significant difference; ***P* < 0.01; *****P* < 0.0001.

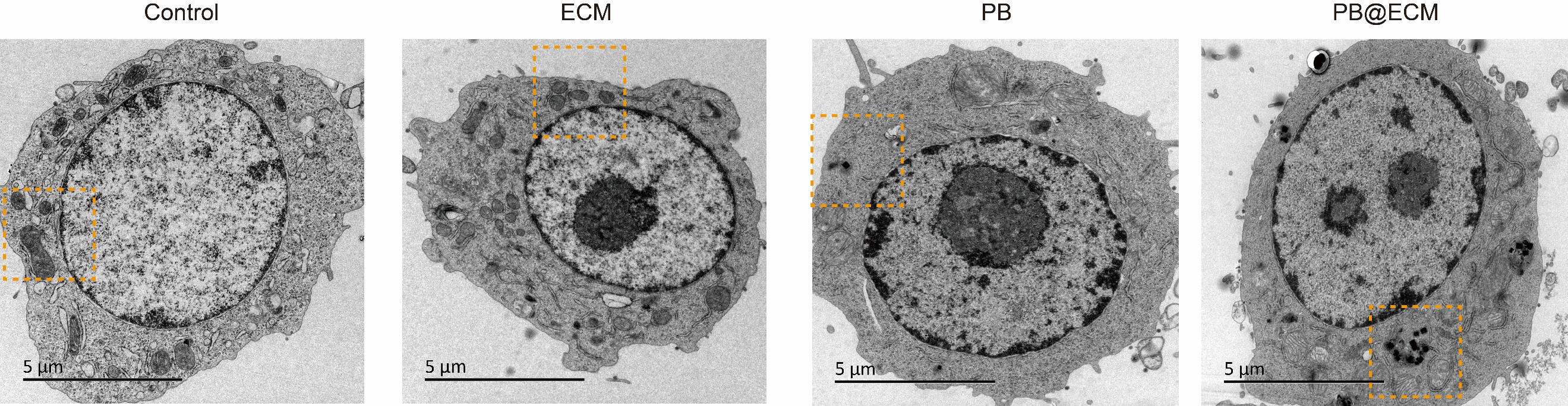

**Supplementary Figure 18.** Electron microscopic images of cells after indicated treatments. Scale bar, 5 μm.

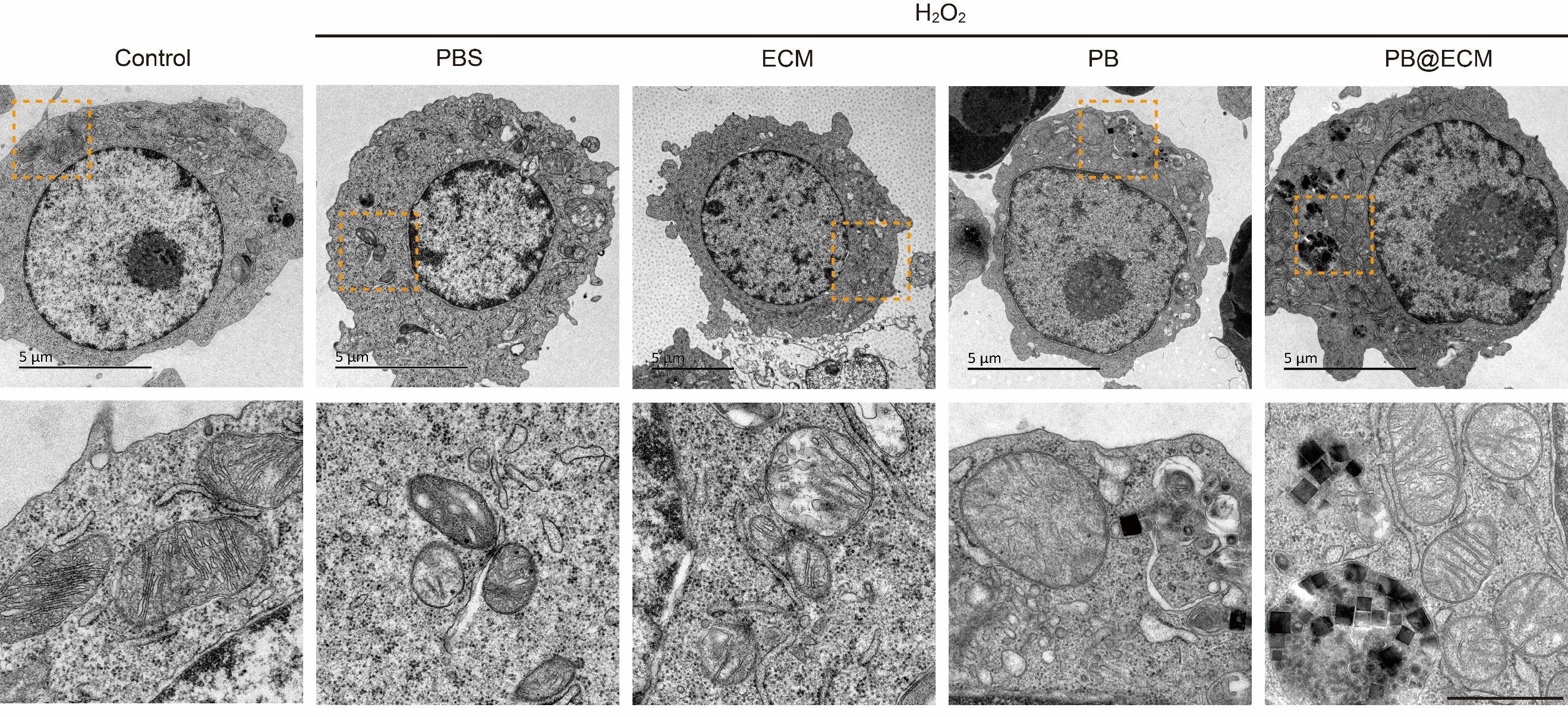

**Supplementary Figure 19.** Electron microscopic images of cells treated with or without hydrogen peroxide, followed by PBS, ECM, PB, or PB@ECM. Scale bars, 5 μm (upper) and 1 μm (lower).

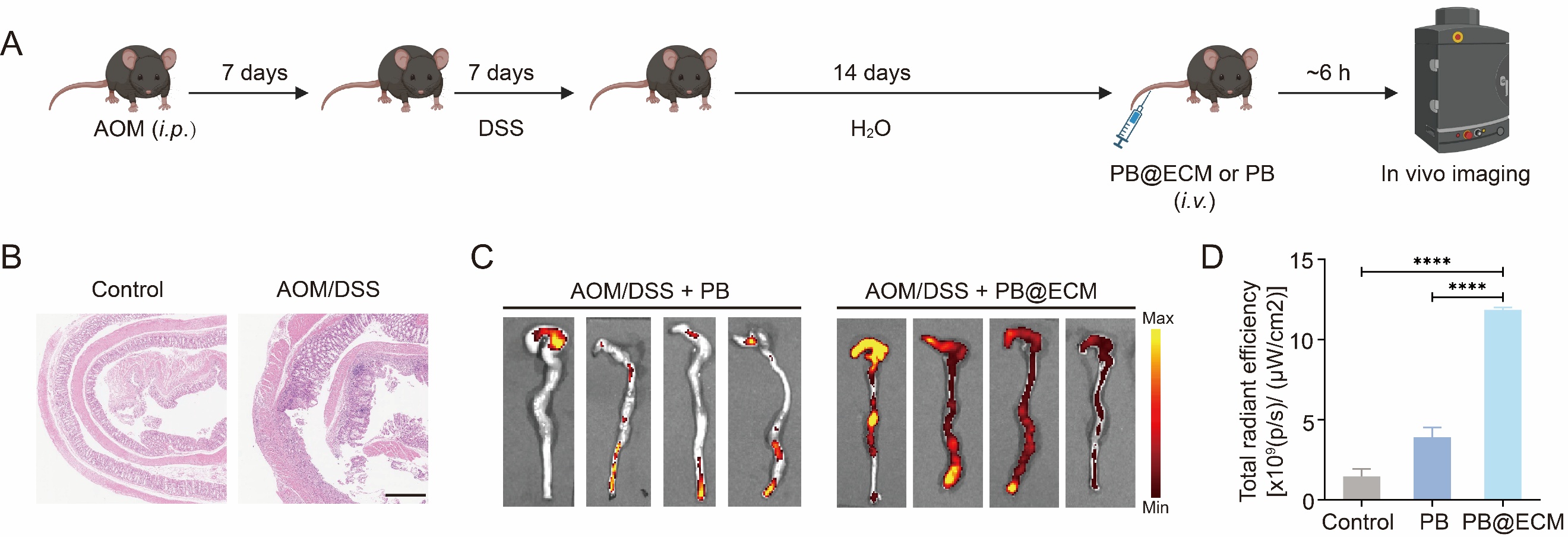

**Supplementary Figure 20. A**, Schematic illustration of targeted experiment using PB@ECM in AOM/DSS mice. Mice were sacrificed 6 h after intravenous injection, and intestines were dissected and imaged using an IVIS imaging system. **B**, Representative H&E-stained colonic sections of mice from control and AOM/DSS groups. Scale bar, 400 μm. **C**, Fluorescence images of colons from AOM/DSS + PB and AOM/DSS + PB@ECM groups. **D**, Quantitative analysis of fluorescence intensity in panel C by Living Image software (n = 4). *****P* < 0.0001.

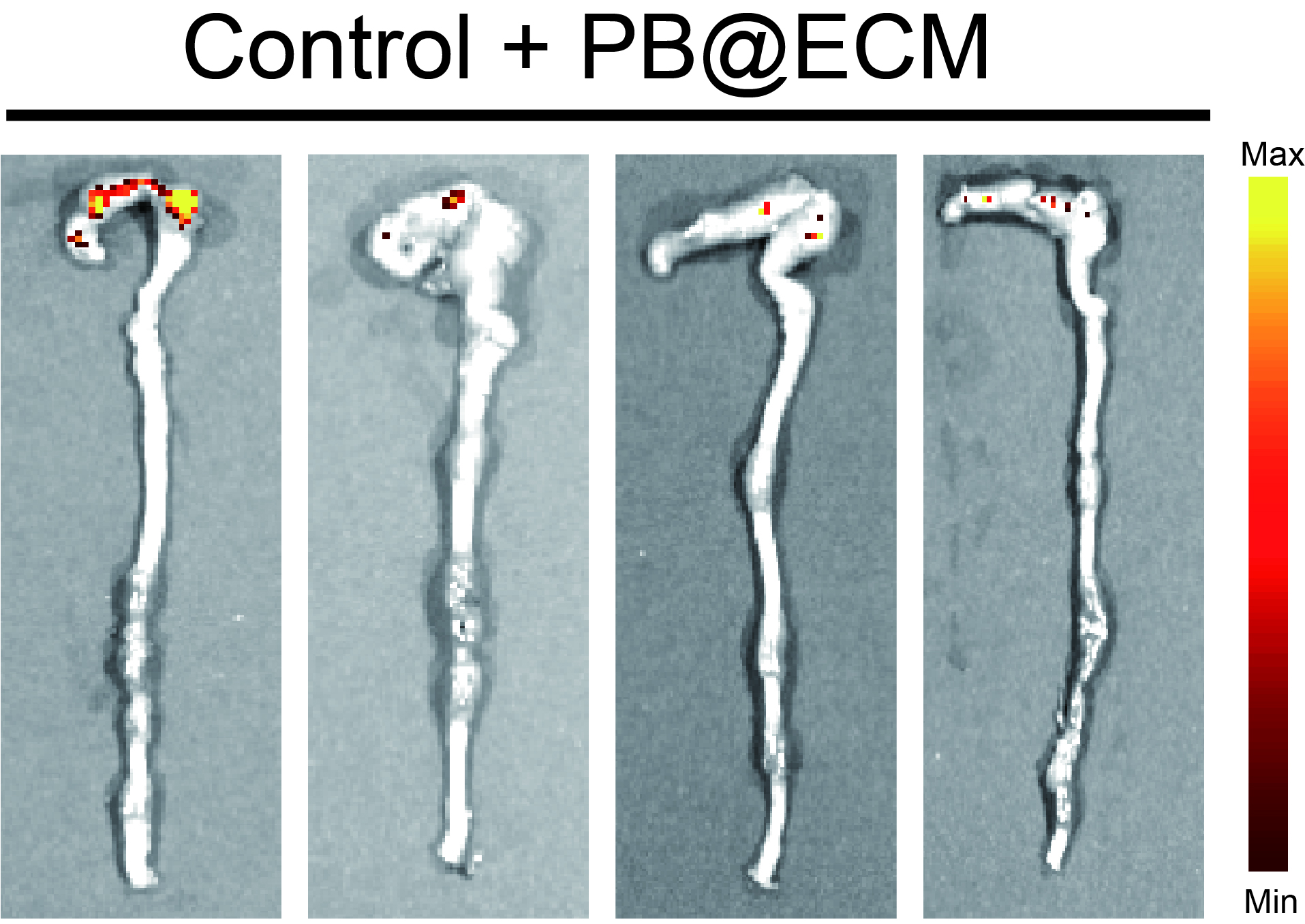

**Supplementary Figure 21.** Fluorescence images of colon in Control + PB@ECM group.

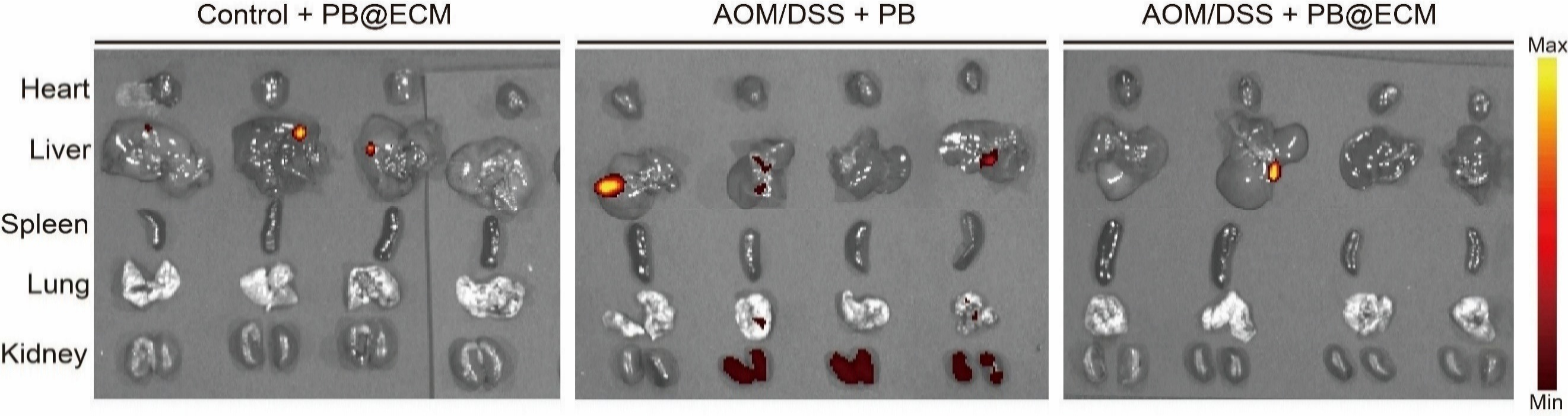

**Supplementary Figure 22.** Fluorescence images of heart, liver, spleen, lung, and kidney after indicated treatments: Control + PB@ECM, AOM/DSS + PB, and AOM/DSS + PB@ECM groups.

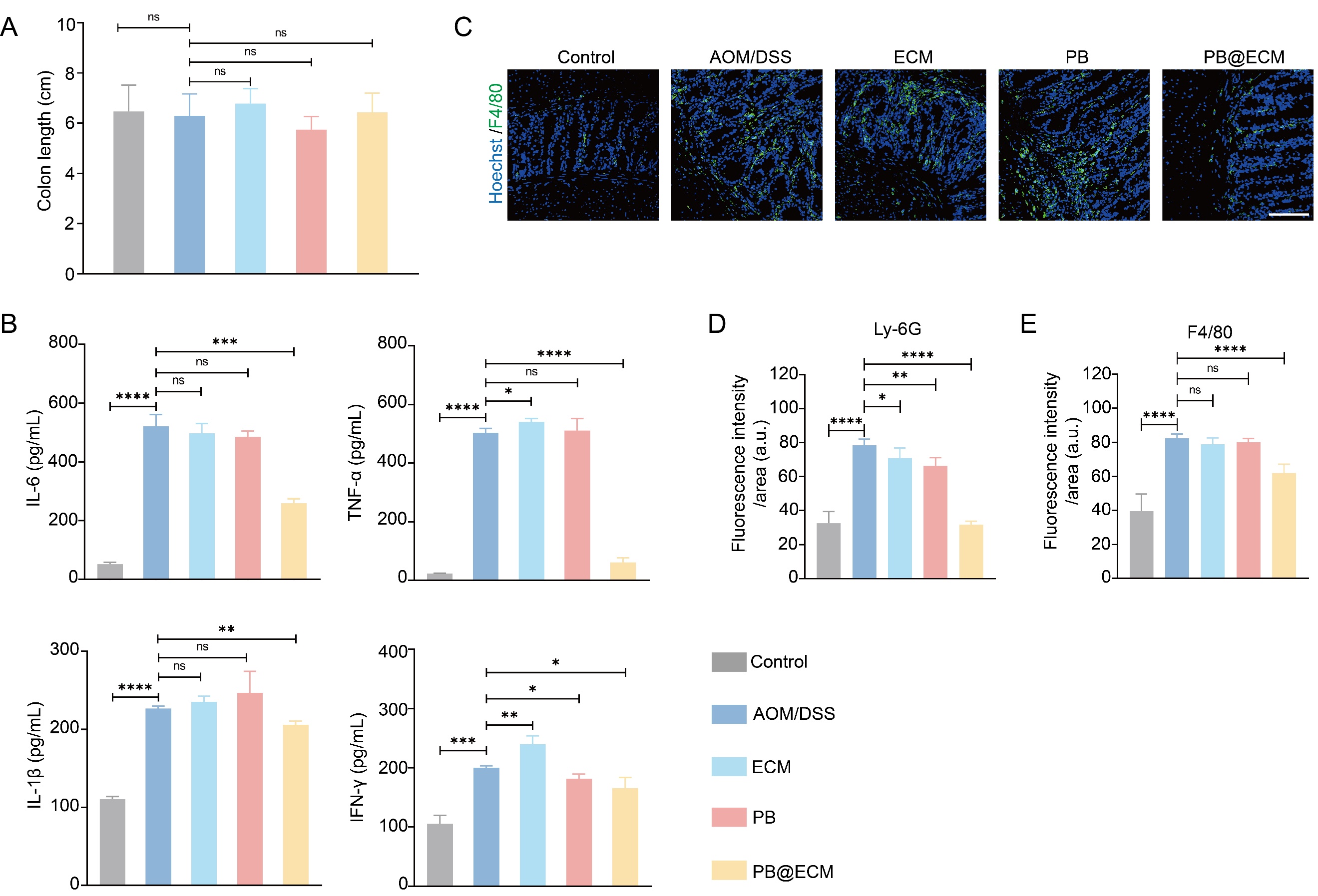

**Supplementary Figure 23. A**, Corresponding colon lengths in indicated groups at early stage (n = 5). **B**, Levels of typical inflammatory cytokines (IL-6, TNF-α, IL-1β, and IFN-γ) in colons of mice on day 28 after indicated treatments at early stage (n = 3). **C**, Representative confocal images of macrophages (F4/80) in colon tissue at early stage. Green, macrophages stained with F4/80; blue, nuclei stained with Hoechst. Scale bar, 100 μm. **D**, Quantitative analysis of Ly-6G immunofluorescence staining (n = 4). **E**, Quantitative analysis of F4/80 immunofluorescence staining (n = 4). Ns, no significant difference; **P* < 0.05; ** *P* < 0.01; ****P* < 0.001; *****P* < 0.0001.

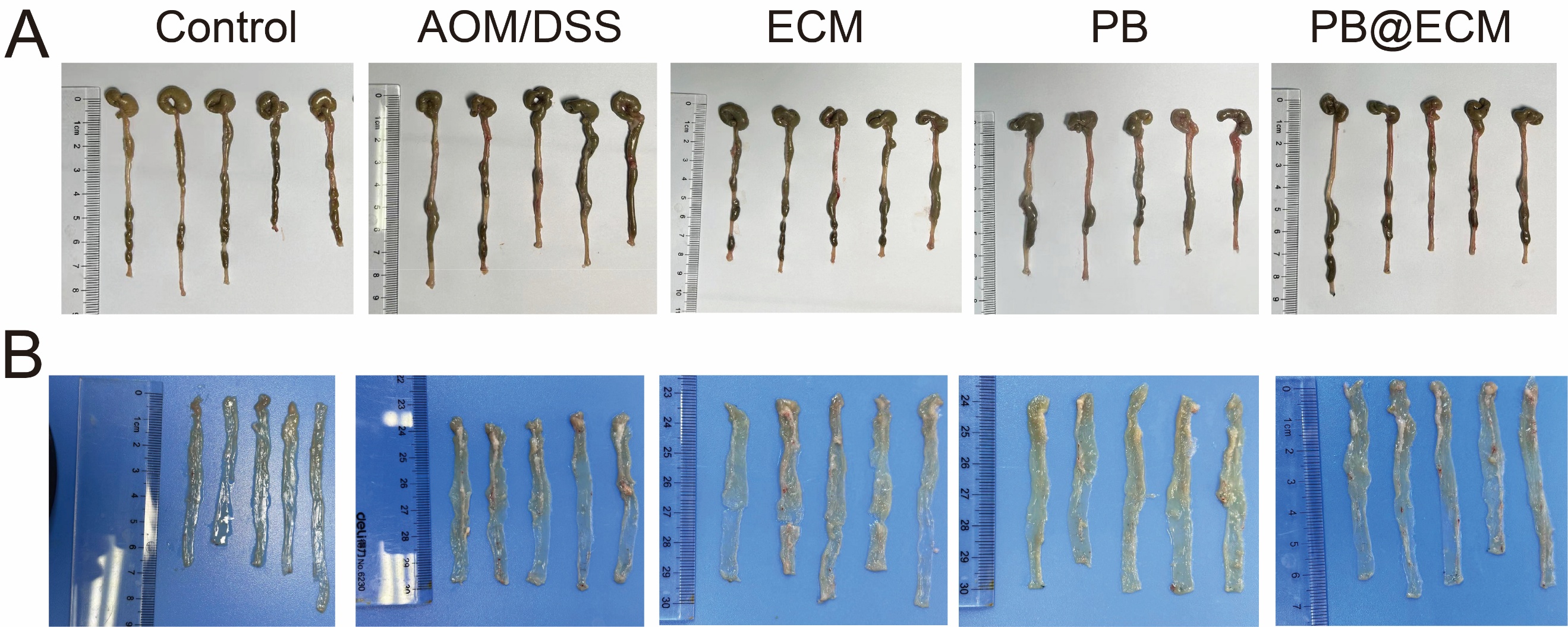

**Supplementary Figure 24.** Representative images of colons (**A**) and anatomical colon (**B**) of indicated groups at early stage.

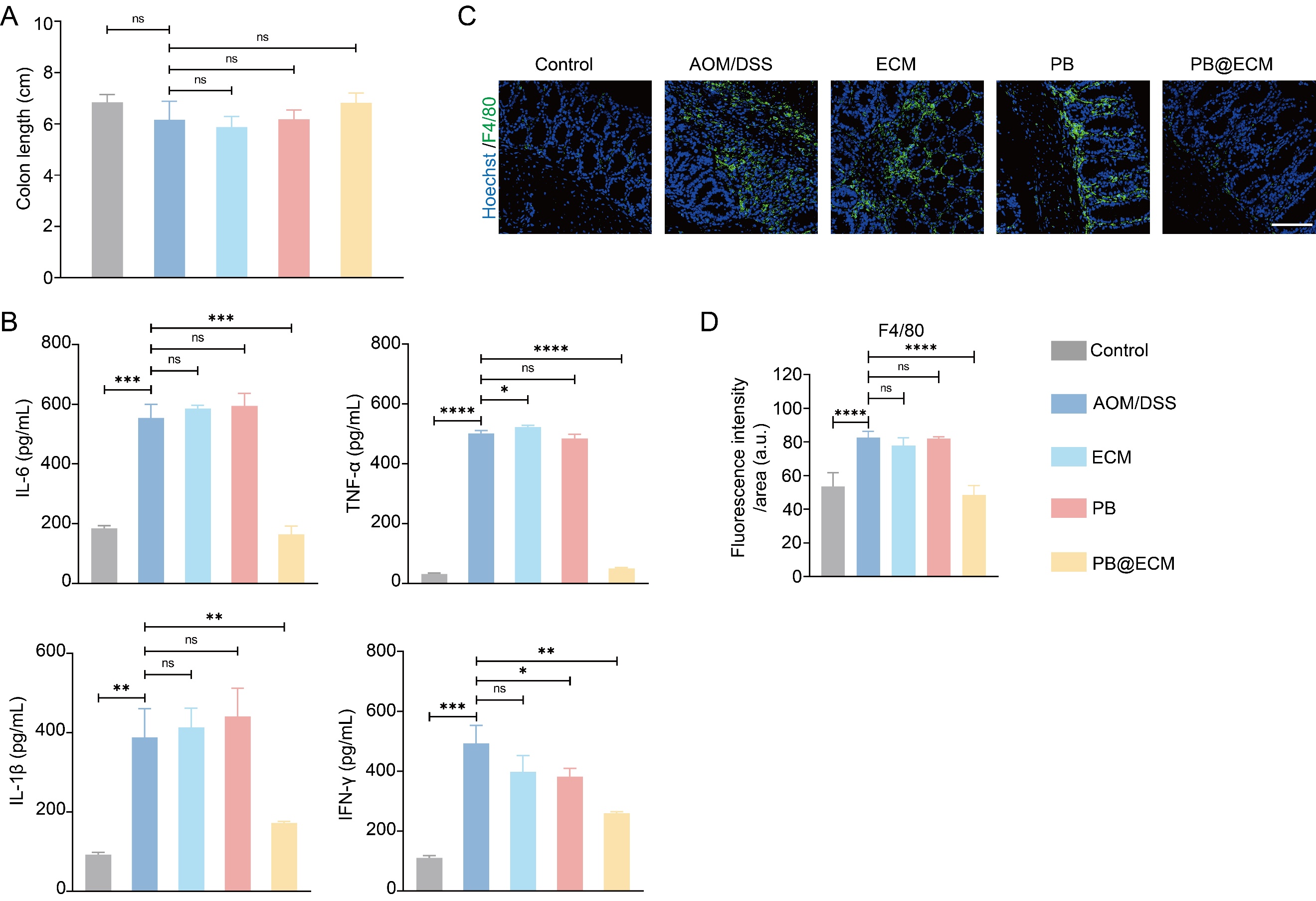

**Supplementary Figure 25. A**, Corresponding colon lengths in indicated groups at intermediate stage (n = 5). **B**, Levels of typical inflammatory cytokines (IL-6, TNF-α, IL-1β, and IFN-γ) in colons of mice on day 49 after indicated treatments at intermediate stage (n = 3). **C**, Representative confocal images of macrophages (F4/80) in colon tissue at intermediate stage. Green, macrophages stained with F4/80; blue, nuclei stained with Hoechst. Scale bar, 100 μm. **D**, Quantitative analysis of F4/80 immunofluorescence staining (n = 4). NS, no significant difference; **P*< 0.05; ***P*< 0.01; ****P* < 0.001; *****P*< 0.0001.

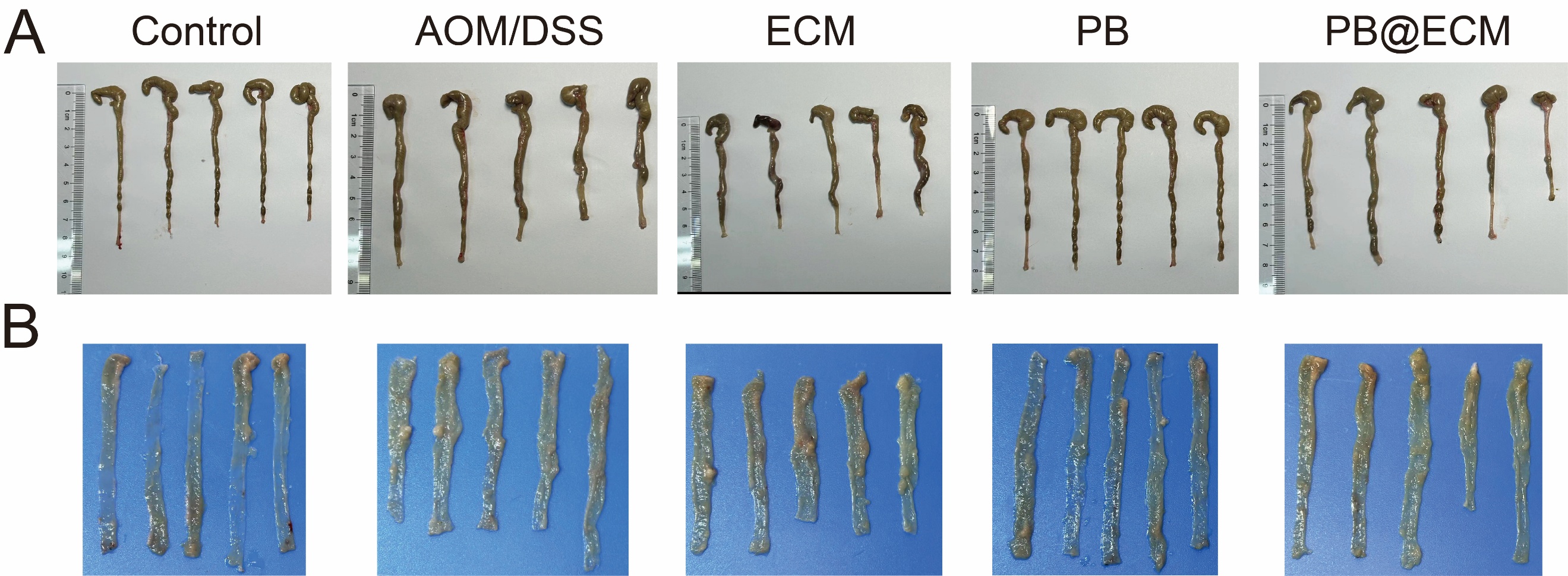

**Supplementary Figure 26.** Representative images of colons (**A**) and anatomical colon (**B**) of indicated groups at intermediate stage.

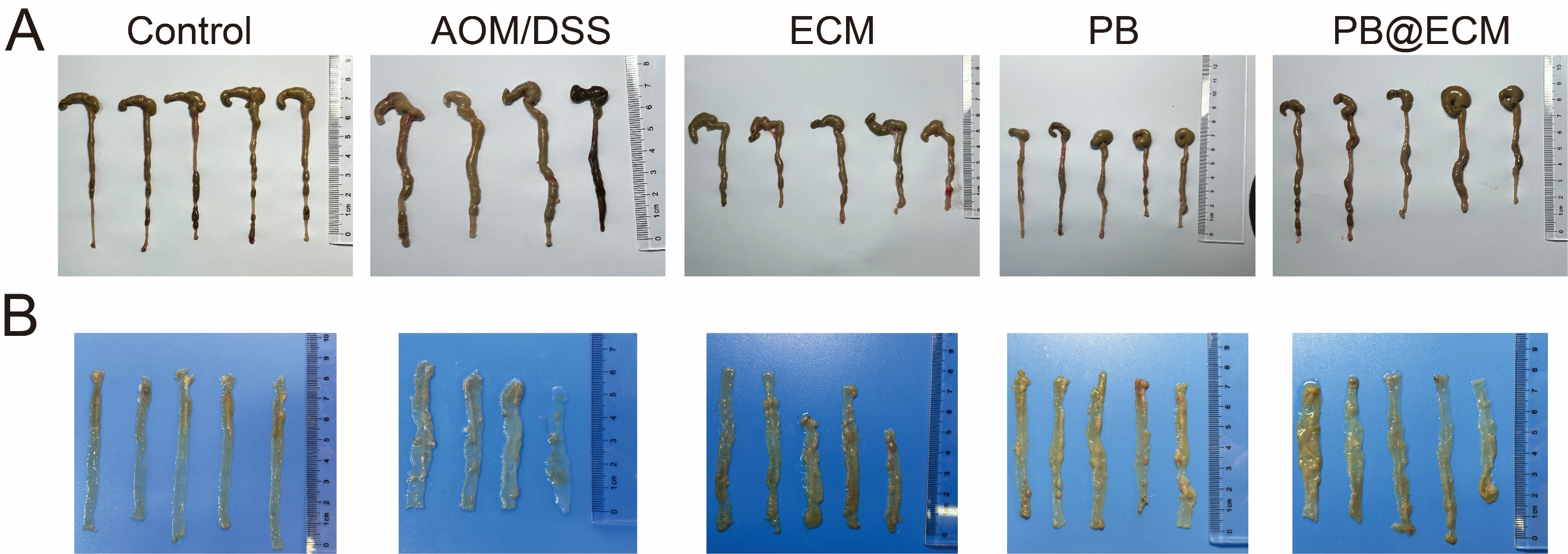

**Supplementary Figure 27.** Representative images of colons (**A**) and anatomical colon (**B**) of indicated groups at late stage. Note: one mouse in the AOM/DSS group died.

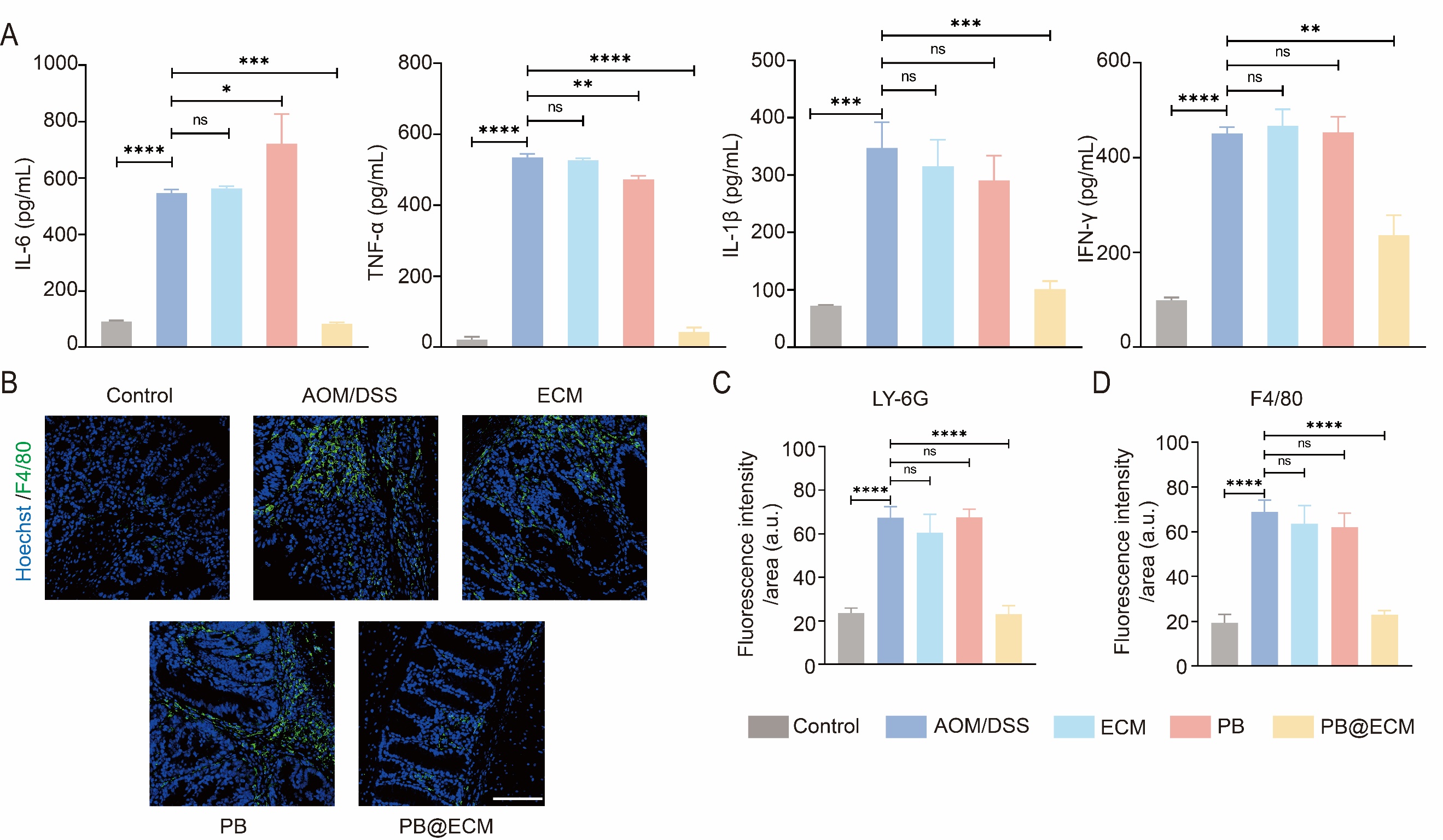

**Supplementary Figure 28. A**, Levels of typical inflammatory cytokines (IL-6, TNF-α, IL-1β, and IFN-γ) in colons of mice on day 70 after indicated treatments at late stage (n = 3). **B**, Representative confocal images of macrophages (F4/80) in colon tissue at intermediate stage. Green, macrophages stained with F4/80; blue, nuclei stained with Hoechst. Scale bar, 100 μm. **C**, Quantitative analysis of Ly-6G immunofluorescence staining (n = 4). **D**, Quantitative analysis of F4/80immunofluorescence staining (n = 4). Ns, no significant difference; **P* < 0.05; ***P* < 0.01; ****P* < 0.001; *****P* < 0.0001.

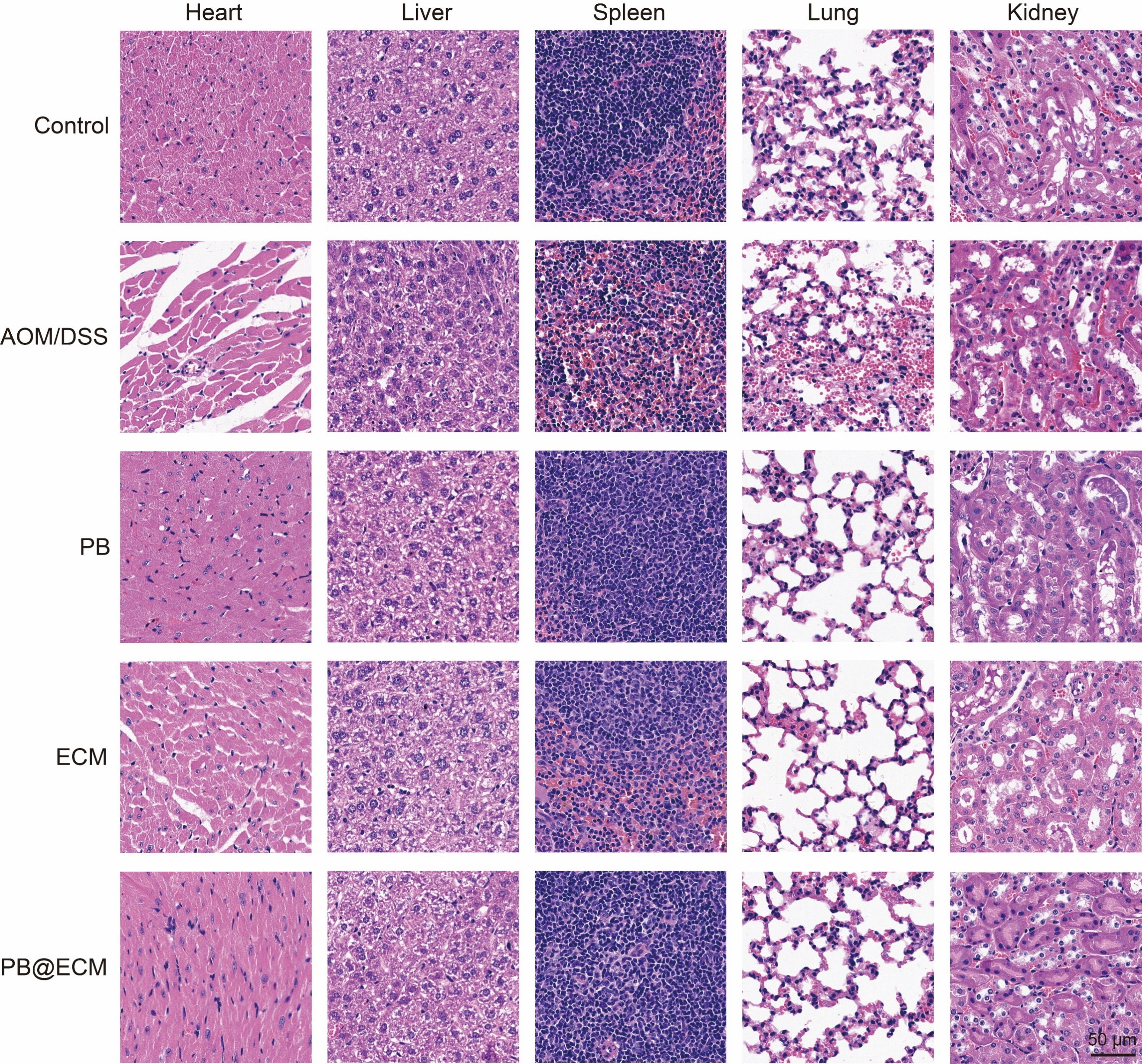

**Supplementary Figure 29.** H&E-stained images of main organs from early stage mice after indicated treatments.

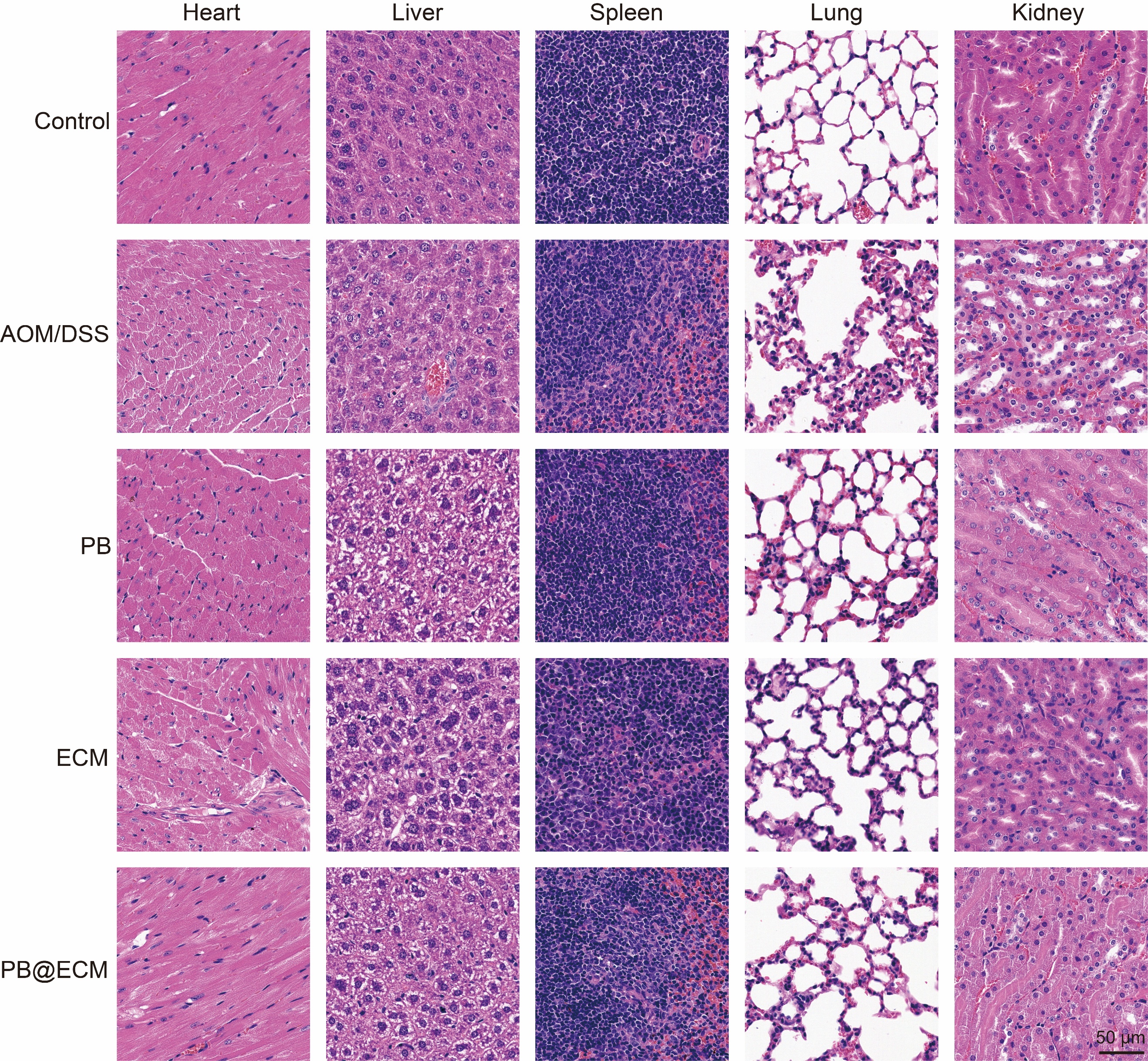

**Supplementary Figure 30.** H&E-stained images of main organs from intermediate stage mice after indicated treatments.

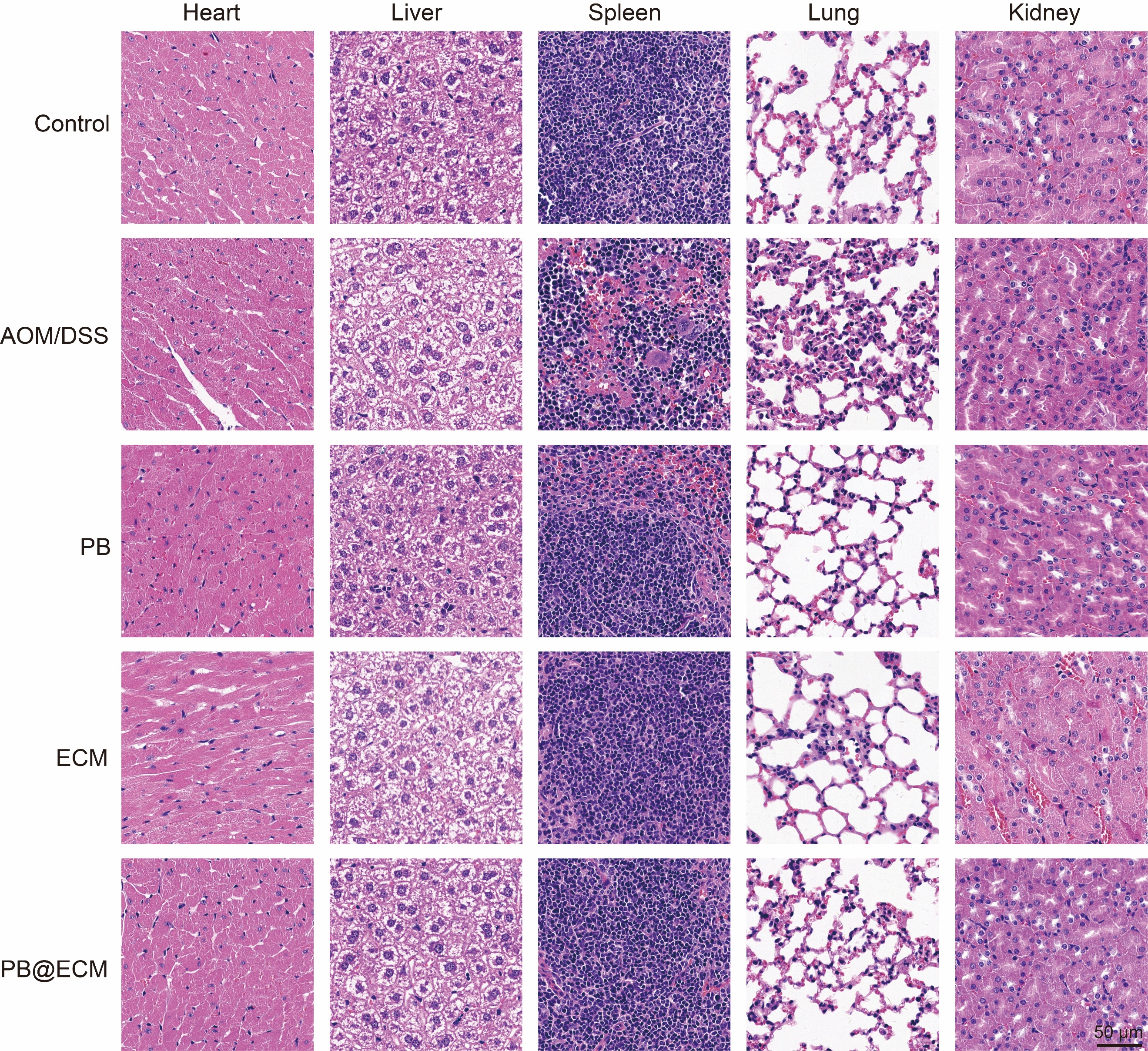

**Supplementary Figure 31.** H&E-stained images of main organs from late stage mice after indicated treatments.

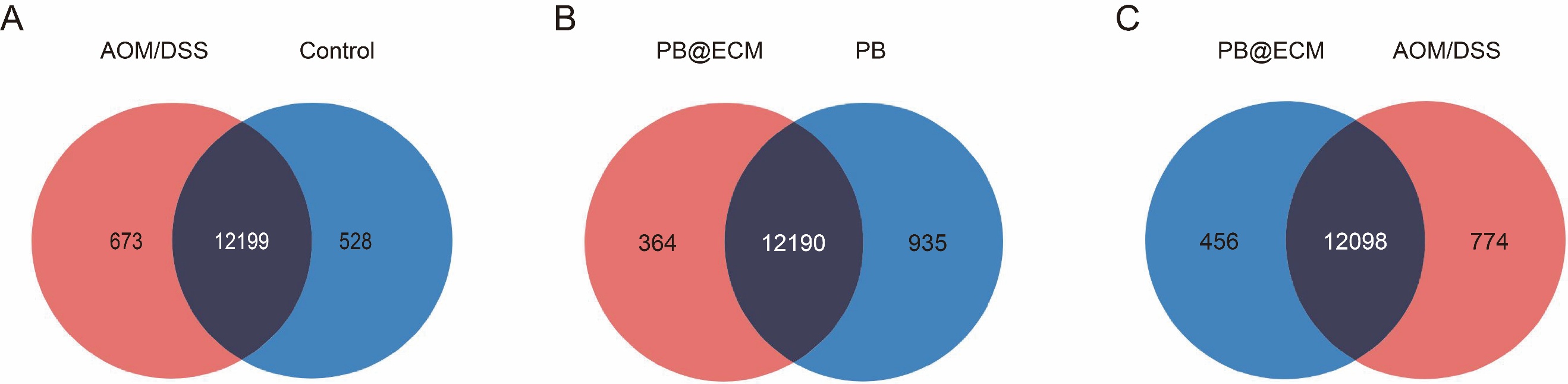

**Supplementary Figure 32.** Venn diagrams of AOM/DSS *vs.* Control (**A**), PB@ECM *vs.* PB (**B**), and PB@ECM *vs.* AOM/DSS groups (**C**).

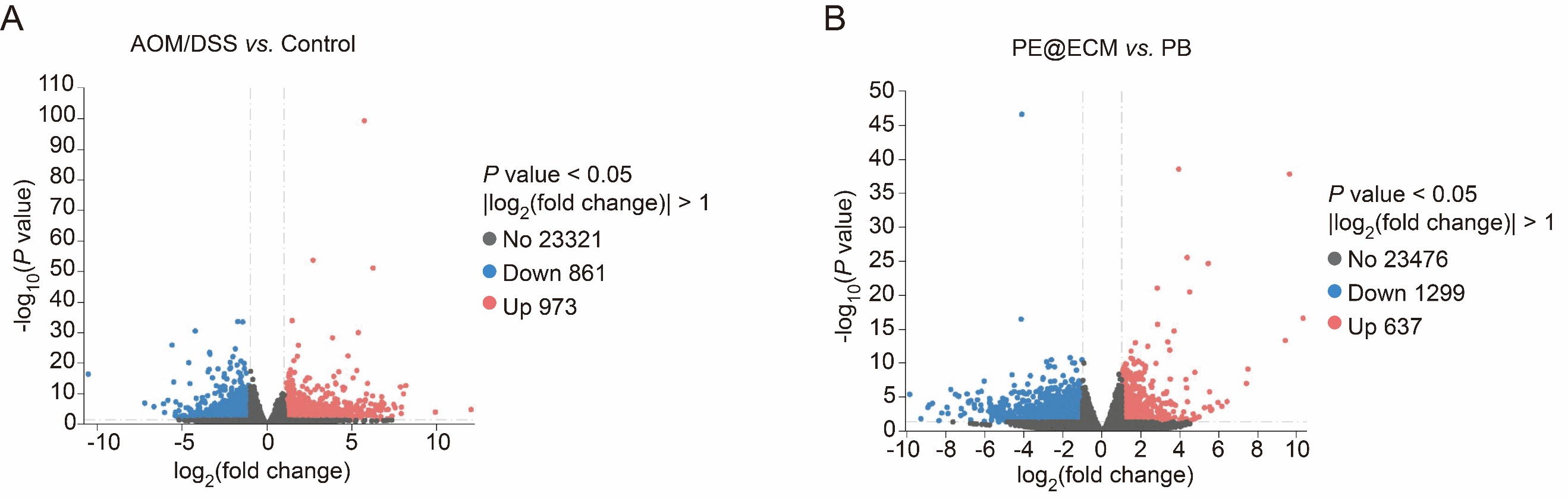

**Supplementary Figure 33.** Volcano plots of differentially expressed genes analysis (DEGs) between AOM/DSS and Control groups (**A**), and between PB@ECM and PB groups (**B**).

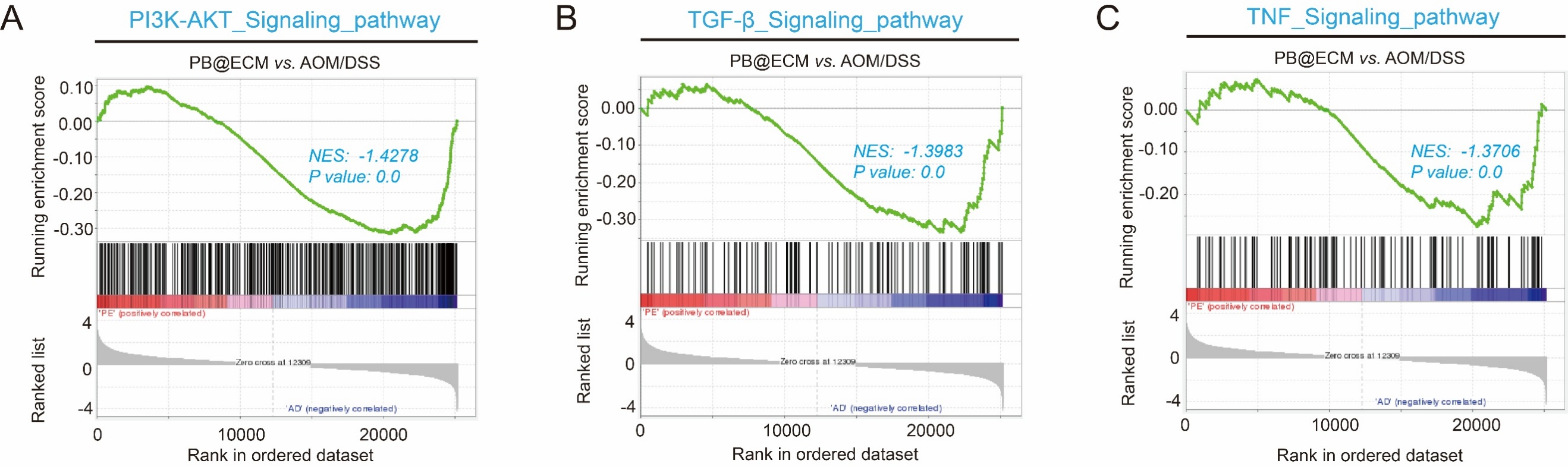

**Supplementary Figure 34.** Gene set enrichment analysis (GSEA) plots showing gene changes of PI3K-AKT (**A**), TGF-β (**B**), and TNF (**C**) in PB@ECM group compared with AOM/DSS group. PI3K-AKT, phosphatidylinositol 3-kinase/Akt pathway; TGF, transforming growth factor; TNF, tumor necrosis factor.

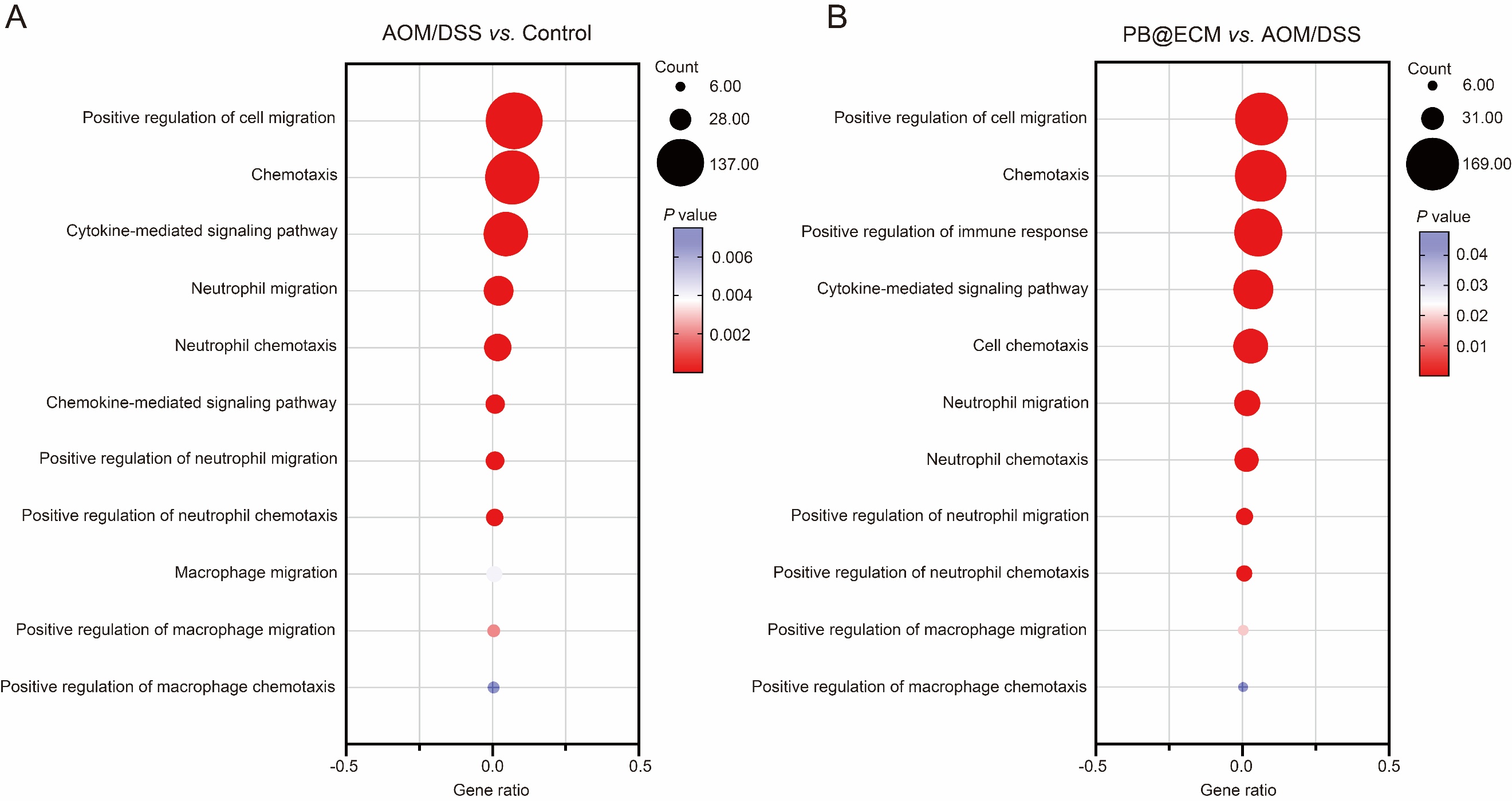

**Supplementary Figure 35.** Enriched gene ontology analysis (GO) terms of genes significantly upregulated in AOM/DSS group compared with Control group (**A**) and significantly downregulated in PB@ECM group compared with AOM/DSS group (**B**).

### Supplementary Tables 1 - 5

**Supplementary Table 1. IBD Patients' clinical information.**

| Patient | Gender | Age | Source |
| --- | --- | --- | --- |
| IBD1 | Female | 23 | Jinling Hospital |
| IBD2 | Female | 51 | Jinling Hospital |
| IBD3 | Male | 37 | Jinling Hospital |
| Normal1 | Female | 23 | Jinling Hospital |
| Normal2 | Female | 51 | Jinling Hospital |
| Normal3 | Male | 37 | Jinling Hospital |

**Supplementary Table 2. CAC Patients' clinical information.**

| Patient | Gender | Age | Source |
| --- | --- | --- | --- |
| CAC1 | Female | 58 | Jinling Hospital |
| CAC2 | Male | 30 | Jinling Hospital |
| CAC3 | Female | 61 | Jinling Hospital |
| Normal1 | Female | 58 | Jinling Hospital |
| Normal2 | Male | 30 | Jinling Hospital |
| Normal3 | Female | 61 | Jinling Hospital |

**Supplementary Table 3. Target gene sequence (*Cxcr2*) used for plasmid construction.**

| Target gene | Sequence (5’-3’) |
| --- | --- |
| *Cxcr2* | atgggagaattcaaggtggataagttcaacattgaagatttcttcagtggagatcttgatattttcaattatagctctggcatgccctctattctgccagatgctgtcccatgccactcagagaacctggaaatcaacagttatgctgtggttgtaatatacgtcctggtgactctgctgagccttgtggggaactccttggtgatgctggtcatcttatacaaccggagcacctgctctgtcaccgatgtctacctgctgaacctggccattgctgacctgttctttgccctgaccttgcctgtctgggctgcatctaaagtaaatggatggacttttggctcaaccctgtgcaagatattctcatacgtgaaggaggtaccttctacagcagtgttctgctactagcctgcatcagcatggaccgctacctggccattgtacatgccacaagtacactgatccagaagagacacttggtcaagtttgtgtgcatagccatgtggttactatcagtaattctggccctgcccatcttaattctacgaaatcctgttaaggtaaacctttctaccttagtctgctatgaggatgtaggtaacaatacatcccgtttgagggtcgtactgcgtatcctgcctcagacttttggcttcctcgtgccgctgctcatcatgctgttctgctacgggttcacactgcgcaccctctttaaggcccacatggggcagaagcaccgggccatgcgggtcatcttcgctgtcgtccttgtcttcctgctctgctggctgccctacaacctggttctgttcacagacaccctcatgagaaccaagctgatcaaggagacctgtgagcgccgcgatgacattgacaaggccttgaatgctacggagattcttggcttcctccacagctgccttaaccccatcatctatgcctttattggccagaaatttcgccatggacttctcaagatcatggctacttatggccttgtcagcaaggagttcttagccaaggagggaaggccttcttttgttagctcgtcttcagcaaacacctctactaccctctaa |

**Supplementary Table 4. Primers used to qRT-PCR.**

| Mouse | Forward primer (5’-3’) | Reverse primer (5’-3’) |
| --- | --- | --- |
| *Cxcr2* | CCCTGCCCATCTTAATTCTACG | AGGAAGCCAAAAGTCTGAGG |
| *β-Actin* | GGTGTGATGGTGGGAATGGG | ACGGTTGGCCTTAGGGTTCAG |

**Supplementary Table 5. Antibody information.**

| Antibody | Source | Application |
| --- | --- | --- |
| CXCR2 | Affinity Biosciences | WB (1:200) |
| F4/80 | Servicebio | IF (1:100) |
| Ly-6G | Servicebio | IF (1:100) |
